# Transport engineering for improving production and secretion of valuable alkaloids in *Escherichia coli*

**DOI:** 10.1101/2021.02.01.429260

**Authors:** Yasuyuki Yamada, Miya Urui, Hidehiro Oki, Kai Inoue, Haruyuki Matsui, Yoshito Ikeda, Akira Nakagawa, Fumihiko Sato, Hiromichi Minami, Nobukazu Shitan

**Author notes:** Correspondence: Nobukazu Shitan, (NS).

## Abstract

Metabolic engineering of microorganisms to produce specialized plant metabolites has been established. However, these methods are limited by low productivity and the intracellular accumulation of metabolites. Here, we aimed to use transport engineering for producing reticuline, an important intermediate in the alkaloid biosynthetic pathway. We established a reticuline-producing *Escherichia coli* strain and introduced a multidrug and toxic compound extrusion transporter, *Arabidopsis* AtDTX1, into it. AtDTX1 was selected due to its suitable expression in *E. coli* and its reticuline-transport activity. Expression of AtDTX1 significantly enhanced reticuline production by 11-fold; produced reticuline was secreted into the medium. AtDTX1 expression conferred high plasmid stability, and up- or downregulated genes associated with biological processes including metabolic pathways for reticuline biosynthesis, leading to a high production and secretion of reticuline. The successful application of a transporter for alkaloid production suggests that the transport engineering approach may improve the biosynthesis of specialized metabolites via metabolic engineering.

## Introduction

Specialized plant secondary metabolites perform diverse functions owing to the variety in their chemical structures. Many specialized metabolites have been used as medicinal resources ^1^. However, meeting the commercial demand for these metabolites is difficult owing to their low concentration in plant cells, danger of extinction of plants, and cost-ineffective methods of chemical synthesis. To circumvent these problems, biosynthetic enzymes for generating these useful metabolites have been studied, and the genes for the corresponding enzymes have been isolated. Further progress in this field has enabled the production of useful compounds by introducing biosynthetic genes into microorganisms such as *Escherichia coli* (*E. coli*) and yeast by metabolic engineering or synthetic biology ^2,3,4^. For example, artemisinic acid, the precursor of anti-malarial drug artemisinin ^5^; thebaine ^6,7^, an important opiate; cannabinoid ^8^, a potential medical compound; colchicine ^9^, a medicine used for treating gout; and tropane alkaloids ^10^, which act as neurotransmitter inhibitors, have been produced in microorganisms.

Although microbial production of central and specialized metabolites is now possible, growth retardation and low productivity have been reported in certain cases, probably due to the cytotoxicity of the substrates or products, or due to negative feedback of the biosynthetic enzymes. For example, yeast cells expressing a plant prenyltransferase and producing a prenylated flavonoid showed decreased growth and production of the end product when the cells were subjected to high concentration of naringenin, a flavonoid substrate ^11^. In another case, yeast stress-responsive genes, including multidrug transporter genes, were highly upregulated in yeast cells induced for artemisinic acid production ^12^. Furthermore, most metabolites accumulate in the cells, which necessitate their extraction from cells and purification from various cellular metabolites. Therefore, additional approaches for alleviating these problems are necessary.

One approach involves the use of transporters that efflux metabolites from cells. Introduction of transporters in metabolite-producing microorganisms might increase production via removal of negative feedback inhibition and enable efficient recovery from the medium. Several examples of successful transport engineering have been reported for central metabolites such as alcohols ^13^, alkanes ^14^, and fatty alcohols ^15^. However, for specialized metabolites, most of which are valuable as medicinal resources and are toxic to microorganisms owing to their strong biological activities, reports regarding the use of efflux pumps are scarce. This is probably due to the dearth of knowledge regarding plant transporters for specialized metabolites. However, certain researchers have isolated transporters implicated in the transport of specialized metabolites ^16,17,18,19,20,21^, which include ATP-binding cassette (ABC) transporters that transport substrates using the energy obtained from ATP hydrolysis ^22^; multidrug and toxic compound extrusion (MATE) transporters, which efflux substrates as proton antiporters ^23,24^; nitrate transporter 1/peptide transporter family (NPF) members, which import substrates as proton symporters ^25^; and purine permease (PUP) members, which import substrates as proton symporters (Supplementary Fig. 1) ^26^. We have studied the transport mechanisms of alkaloids and characterized the function of several transporters using microorganisms; for example, the ABCB-type ABC transporters are responsible for berberine translocation in *Coptis japonica* ^27,28^ and four MATE transporters and one PUP transporter are required for nicotine transport in *Nicotiana tabacum* ^29,30,31,32^. In addition, other groups have reported several transporters that transport various alkaloids ^19, 20^. Furthermore, using metabolic engineering, we have established *E. coli* that can produce reticuline ^33^, an important intermediate for various benzylisoquinoline alkaloids such as morphine and berberine, via three engineered pathways, 1) an L-tyrosine-overproducing pathway via glycolysis, pentose phosphate pathway, and shikimic acid pathway; 2) a pathway producing dopamine from L-tyrosine along with the tetrahydrobiopterin (BH_4_)-synthesis pathway; 3) a reticuline-producing pathway from dopamine (Fig. 1). Therefore, we hypothesized that the production and secretion of specialized metabolites such as reticuline can be enhanced by combining transport engineering with metabolic engineering in *E. coli* using the information available regarding alkaloid transporters.

**Fig. 1.**
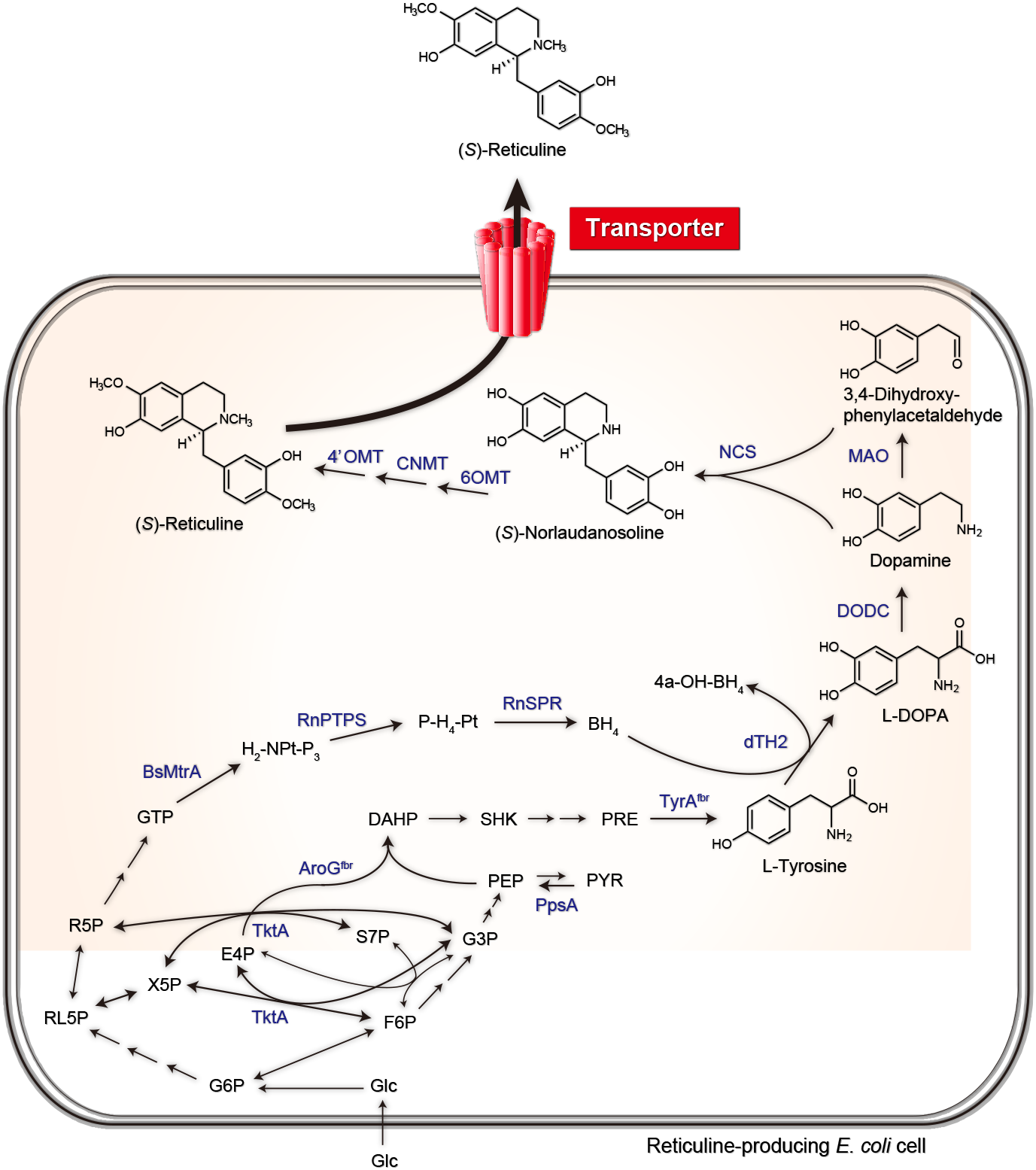
Simplified scheme showing the biosynthetic pathway in *E. coli* leading to high production of reticuline, including the reticuline transportation step. Reticuline is synthesized from simple carbon sources, i.e., glucose or sucrose, via the sequential action of enzymes dTH2, DODC, MAO, NCS, 6OMT, CNMT, and 4′OMT. As an efflux transporter of reticuline, AtDTX1 was introduced in this reticuline-producing *E. coli* strain. Abbreviations of the compounds and enzymes are as follows: 4a-H-BH_4_, 4a-hydroxytetrahydrobiopterin; AroG^fbr^, 2-dehydro-3-deoxyphosphoheptonate aldolase; BH4, tetrahydrobiopterin; BsMtrA, GTP cyclohydrolases I; CNMT, coclaurine *N*-methyltransferase; dTH2, tyrosine hydroxylase; DAHP, 3-deoxy-D-arabino-heptulosonate-7-phosphate; DODC, L-DOPA decarboxylase; E4P, D-erythrose 4-phosphate; F6P, D-fructose 6-phosphate; G3P, D-glyceraldehyde 3-phosphate; G6P, D-glucose 6-phosphate; H_2_-NPt-P_3_, 7,8-dihydroneopterin triphosphate; MAO, monoamine oxidase; 4′OMT, 3′-hydroxy-*N*-methyl-(*S*)-coclaurine 4′-*O*-methyltransferase; 6OMT, norcoclaurine 6-*O*-methyltransferase; PEP, phosphoenolpyruvate; P-H4-Pt, 6-pyruvoyltetrahydropterin; PpsA, phosphoenolpyruvate synthetase; PRE, prephenate; PYR, pyruvate; R5P, D-ribose 5-phosphate; RL5P, D-ribulose 5-phosphate; RnPTPS, 6-pyruvoyltetrahydropterin synthase; RnSPR, sepiapterin reductase; S7P, D-sedoheptulose 7-phosphate; SHK, shikimate; TktA, transketolase; TyrA^fbr^, chorismate mutase-prephenate dehydrogenase (feedback-resistant); and X5P, D-xylose 5-phosphate.

In this study, we aimed to investigate the effect of introducing a MATE transporter, AtDTX1 from *Arabidopsis*, selected due to its expression and transport activity in *E. coli*, on reticuline production and efflux in alkaloid-producing *E. coli* (Fig. 1). Our observations suggested that the combination of transport engineering and metabolic engineering is a powerful tool that can be used for enhancing the productivity of specialized plant metabolites in microorganisms. This technology would lead to high production and a stable supply of useful pharmaceutical compounds in the future.

## Results

### Selection of appropriate transporters for transport engineering

For applying transport engineering to alkaloid production, appropriate transporters involved in transporting specialized metabolites must be selected ^16,18,20^. First, we focused on the MATE family of plant transporters known to transport specialized metabolites ^20,23^, as members of this protein family efflux substrates from the cytosol (Supplementary Fig. 1) and their microbial expression is well studied. Among these, we focused on AtDTX1 from *Arabidopsis thaliana* and NtJAT1 from *Nicotiana tabacum* (Supplementary Fig. 2). AtDTX1 transports plant-derived alkaloids such as berberine and palmatine, whose chemical structures are relatively similar to that of reticuline (Supplementary Fig. 3). AtDTX1 was isolated from a functional screen using *E. coli* ^34^, indicating that it is well expressed in *E. coli* NtJAT1 is expressed well in *Saccharomyces cerevisiae* and localizes to the plasma membrane, where it shows substrate specificity for alkaloids such as berberine (Supplementary Fig. 2) ^29^. Therefore, we investigated the expression and reticuline transport activities of AtDTX1 and NtJAT1 in *E. coli* BL21(DE3).

### Expression and reticuline transport activity of AtDTX1 and NtJAT1 in *E. coli*

We investigated whether AtDTX1 and NtJAT1 are adequately expressed and are able to efflux reticuline in *E. coli* Each cDNA was subcloned into pCOLADuet-1, a low copy plasmid, and introduced in *E. coli* BL21(DE3). The transformants and those harboring only the vector control were incubated in Luria Bertani (LB) medium, and isopropyl β-d-1-thiogalactopyranoside (IPTG) (final concentration, 1 mM) was added at OD_600_ = 0.6 to induce the expression of each transporter. A 37 kDa immunoreactive band was detected in the membrane preparation of *E. coli* transformed with *AtDTX1* cDNA using His-antibody and also in Coomassie Brilliant Blue (CBB) staining. This band was absent in the membranes of the vector control cells (Fig. 2a), indicating that AtDTX1 was expressed in *E. coli*. In contrast, a weak band of around 37 kDa was observed for NtJAT1 using the anti-NtJAT1 antibody (Fig. 2b). Next, BL21(DE3) cells expressing AtDTX1 or NtJAT1, or control cells were incubated in LB medium containing reticuline (final concentration, 250 µM), and the intracellular reticuline content was quantitatively analyzed using ultra performance liquid chromatography-mass spectrometry (UPLC-MS). Cells expressing AtDTX1 or NtJAT1 accumulated significantly less content of reticuline than the control cells (Fig. 3). These data suggested that both MATE transporters were expressed at the plasma membrane of *E. coli* cells and effluxed reticuline.

**Fig. 2.**
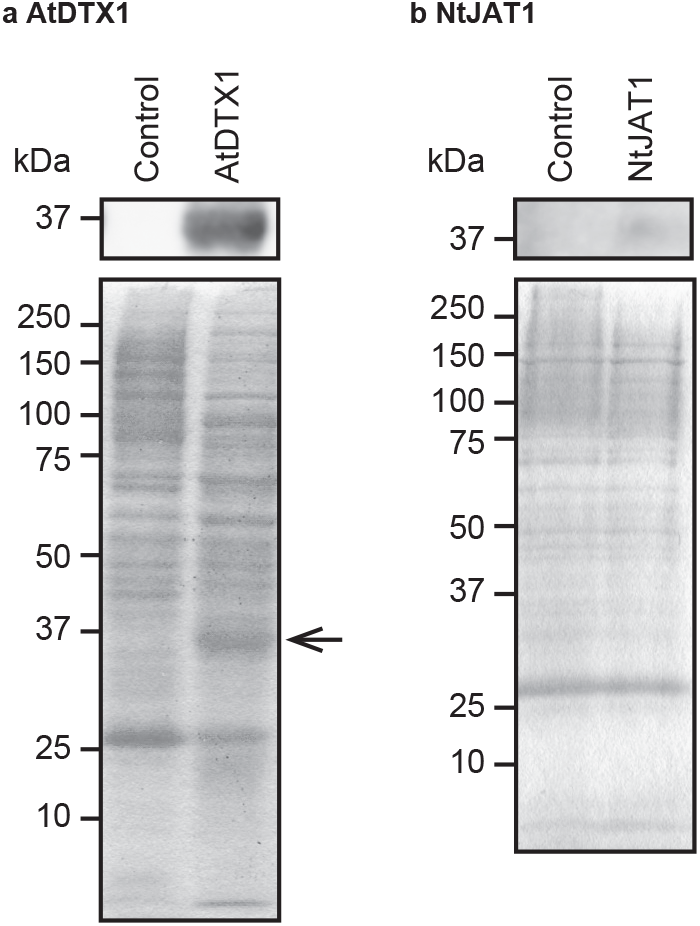
Expression of AtDTX1 and NtJAT1 in *E. coli* BL21(DE3). Expression of MATE transporter was induced by adding IPTG (1 mM) and incubating for 3.5 h. Membrane proteins (10 μg per lane) of *E. coli* expressing AtDTX1, NtJAT1, or vector control were extracted, separated via sodium dodecyl sulfate-polyacrylamide gel electrophoresis, and blotted onto a polyvinylidene difluoride membrane. The membrane was probed with anti-His antibodies against AtDTX1 or anti-NtJAT1 antibodies against NtJAT1. The position of AtDTX1 is marked by an arrowhead.

**Fig. 3.**
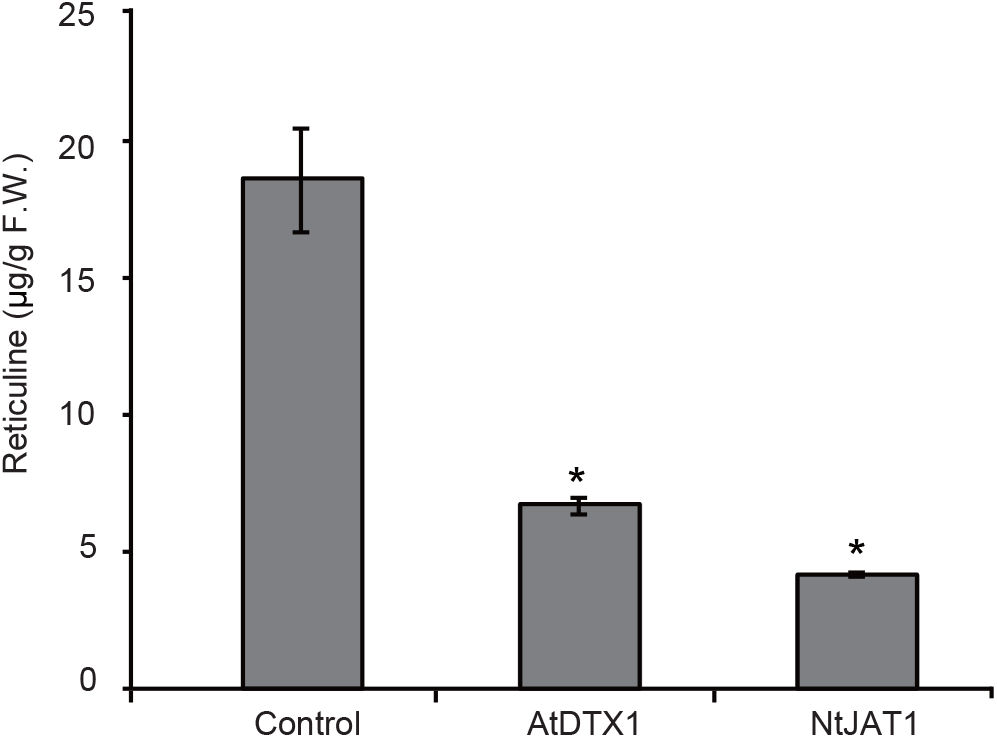
Reticuline transport activity of AtDTX1 or NtJAT1 in *E. coli* BL21(DE3). AtDTX1 or NtJAT1 expression was induced by adding IPTG (1 mM) and incubating for 3 h. Control, or AtDTX1- or NtJAT1-expressing *E. coli* BL21 (DE3) cells were resuspended and cultured in LB medium containing reticuline (250 μM) for 6 h. Results show mean ± standard deviation (SD) of triplicates. Asterisks indicate statistically significant difference compared to the control (ANOVA Bonferroni test; \**P* < 0.01).

### Determination of the conditions for transporter expression and cell culture

During the reticuline transport assay, we observed that the growth of NtJAT1-expressing *E. coli* cells was significantly inhibited. As the expression of membrane proteins tends to reduce cell growth ^13^, the growth rate of *E. coli* cells expressing AtDTX1 or NtJAT1 was determined. In the presence of IPTG (0.1 mM), AtDTX1 expression in BL21(DE3) cells retarded growth slightly, whereas NtJAT1 significantly inhibited growth (Supplementary Fig. 4). As growth defect is undesirable for alkaloid production, only AtDTX1 was used for further analysis.

We introduced AtDTX1 or the control vector in reticuline-producing cells. Reticuline-producing cells were established in this study from the BL21(DE3) strain via the introduction of four vectors with 14 genes related to reticuline biosynthesis as described previously ^33^. Although a different combination of vectors and genes was used (Supplementary Table 1), this compound was produced from glucose, a simple carbon source (Fig. 1). Reticuline-producing cells grew more slowly than the BL21(DE3) strain, probably because these cells harbored multiple vectors and were cultured in the presence of five antibiotics (tetracycline, ampicillin, spectinomycin, chloramphenicol, and kanamycin). AtDTX1 expression did not significantly affect the proliferation of *E. coli* in the presence of 0.1 mM IPTG (Supplementary Fig. 5). A previous report ^35^ has shown that reticuline is produced efficiently using modified LB medium containing K_2_HPO_4_, KH_2_PO_4_, and glycerol (0.4%). Hence, we decided to investigate the reticuline production of these cells using modified LB medium in the presence of 0.1 mM IPTG.

### AtDTX1 significantly improved reticuline production and secretion in the medium

We determined the effect of AtDTX1 expression on cell growth and reticuline production under the conditions described above. At OD_600_ = 0.6, IPTG (0.1 mM) was added to the medium to induce the biosynthetic enzymes for reticuline synthesis and AtDTX1. AtDTX1-expressing cells grew almost similarly as the control cells during the exponential phase; however, they entered the stationary phase earlier than the control cells (Fig. 4). Reticuline levels in the cells and medium were monitored quantitatively using UPLC-MS analysis (Supplementary Fig. 6). Time course analysis (from 0 to 72 h) showed that cellular reticuline content in AtDTX1-expressing *E. coli* cells was significantly higher than that of the vector control (Fig. 5a). At 12 h, the reticuline content in AtDTX1-expressing cells reached 17.7 μg/g fresh weight, which was 3.5-fold higher than that of the control cells (5.1 μg/g fresh weight). Reticuline was detected in the medium of AtDTX1-expressing cells (0.23 mg/L at 8 h) earlier than in the medium of control cells (0.13 mg/L at 12 h). Reticuline content in AtDTX1-expressing cells sharply increased from 12 h to 24 h, and at 24 h, the difference in reticuline content between the two cell lines was 11-fold (14.4 mg/L in AtDTX1-expressing cells and 1.3 mg/L in control cells); this difference was maintained for 72 h (Fig. 5b). These results indicated that AtDTX1 expression significantly enhanced reticuline production and secretion in the medium.

**Fig. 4.**
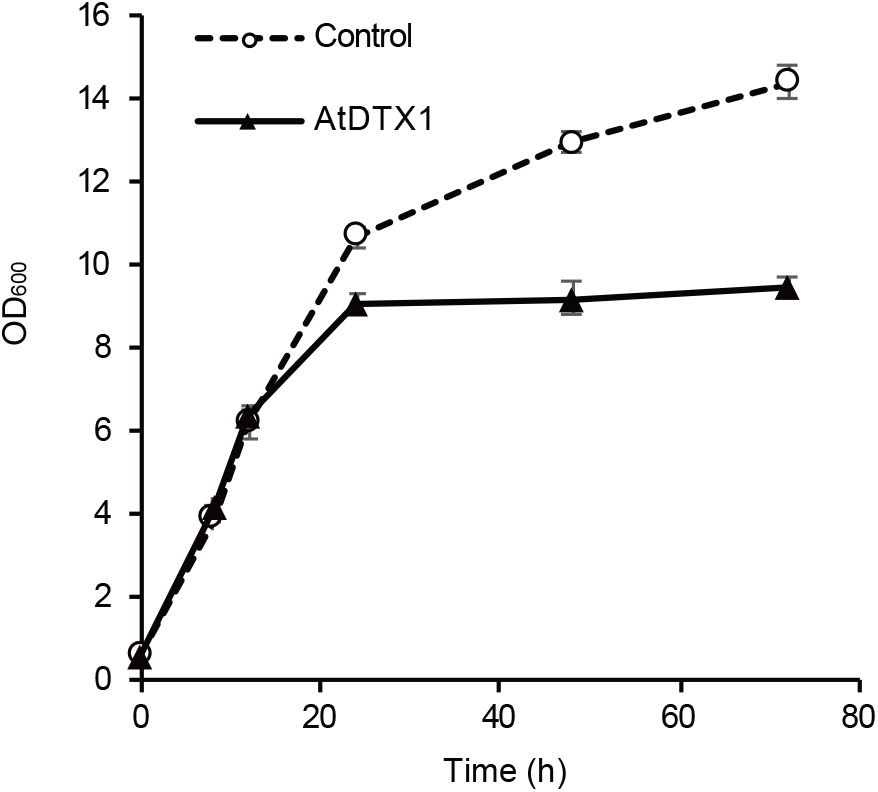
Growth of reticuline-producing *E. coli* after induction of AtDTX1 by IPTG. Growth was evaluated by measuring the optical density at 600 nm. Control (dashed line) and AtDTX1-expressing (solid line) *E. coli* cells were cultured in modified LB medium containing 9.4 g/L K_2_HPO_4_, 2.2 g/L KH_2_PO_4_, 0.4% glycerol, and antibiotics. IPTG (0.1 mM final concentration) was added at OD_600_ = 0.6 (time = 0 h) and sampled at the times indicated. Results show mean ± standard deviation (SD) of triplicates.

**Fig. 5.**
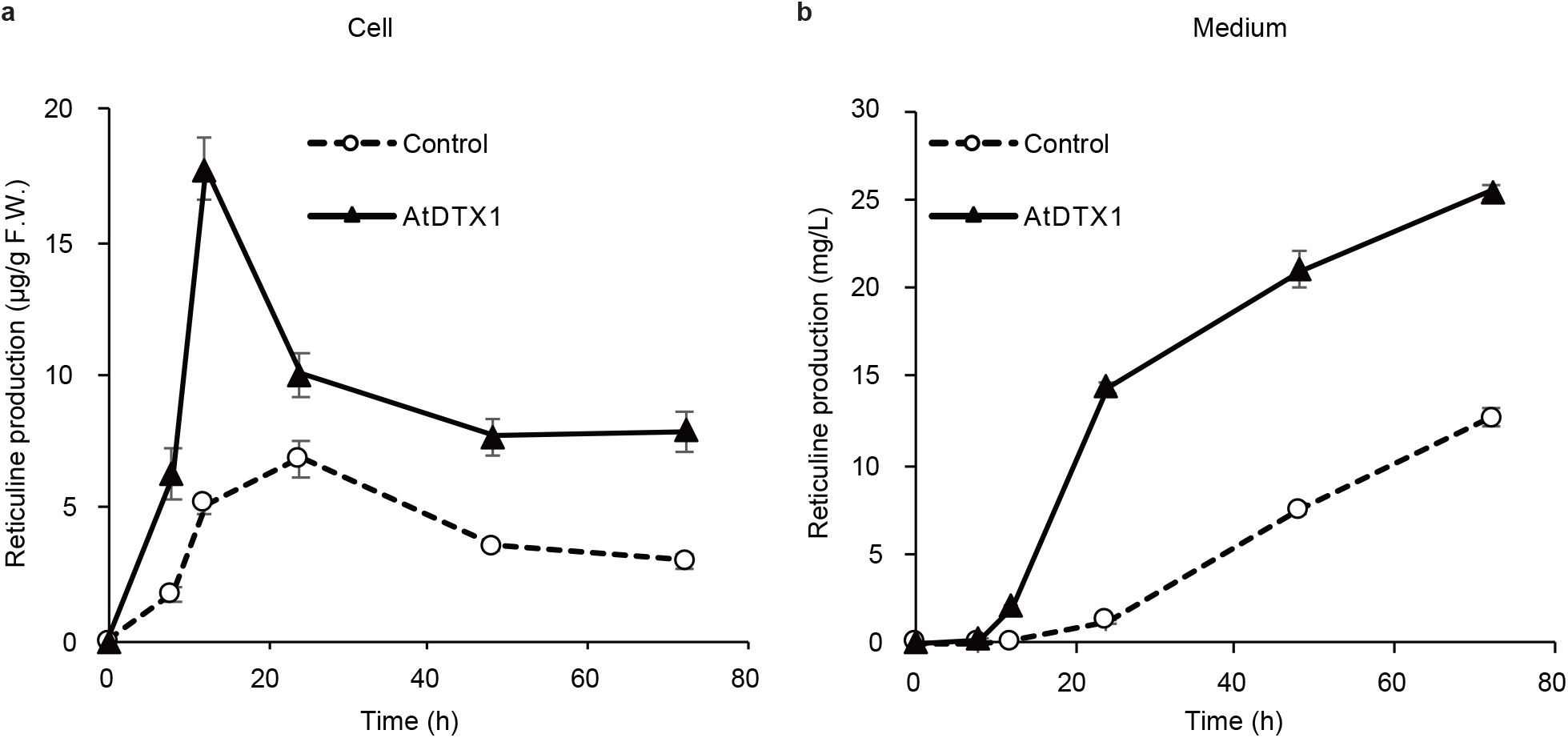
Reticuline production in *E. coli* cells and medium. Time-dependent production of (*S*)-reticuline in *E. coli* cells (a) and medium (b). Control (dashed line) and AtDTX1-expressing (solid line) *E. coli* cells were incubated in modified LB medium. IPTG (final concentration 0.1 mM) was added at OD_600_ = 0.6 and sampled at the times indicated. Results show mean ± standard deviation (SD) of triplicates.

### Plasmid stability and transcriptomic analysis of AtDTX1-expressing *E. coli* cells

A previous study regarding metabolic engineering of yeast cells for the biosynthesis of artemisinic acid reported that the end product, artemisinic acid, lowered plasmid stability and the productivity of this compound ^12^. Therefore, to understand the mechanism underlying the enhancement of reticuline production due to AtDTX1 expression, we assessed plasmid stability in reticuline-producing cells. At 4 h, both cells (AtDTX1-expressing and control) maintained almost similar plasmid numbers. However, at 24 h, the AtDTX1-expressing cells maintained plasmids in 91.7% of the cells, which was significantly higher than the proportion of the control cells (64.3%) (Fig. 6, Supplementary Fig. 7).

**Fig. 6.**
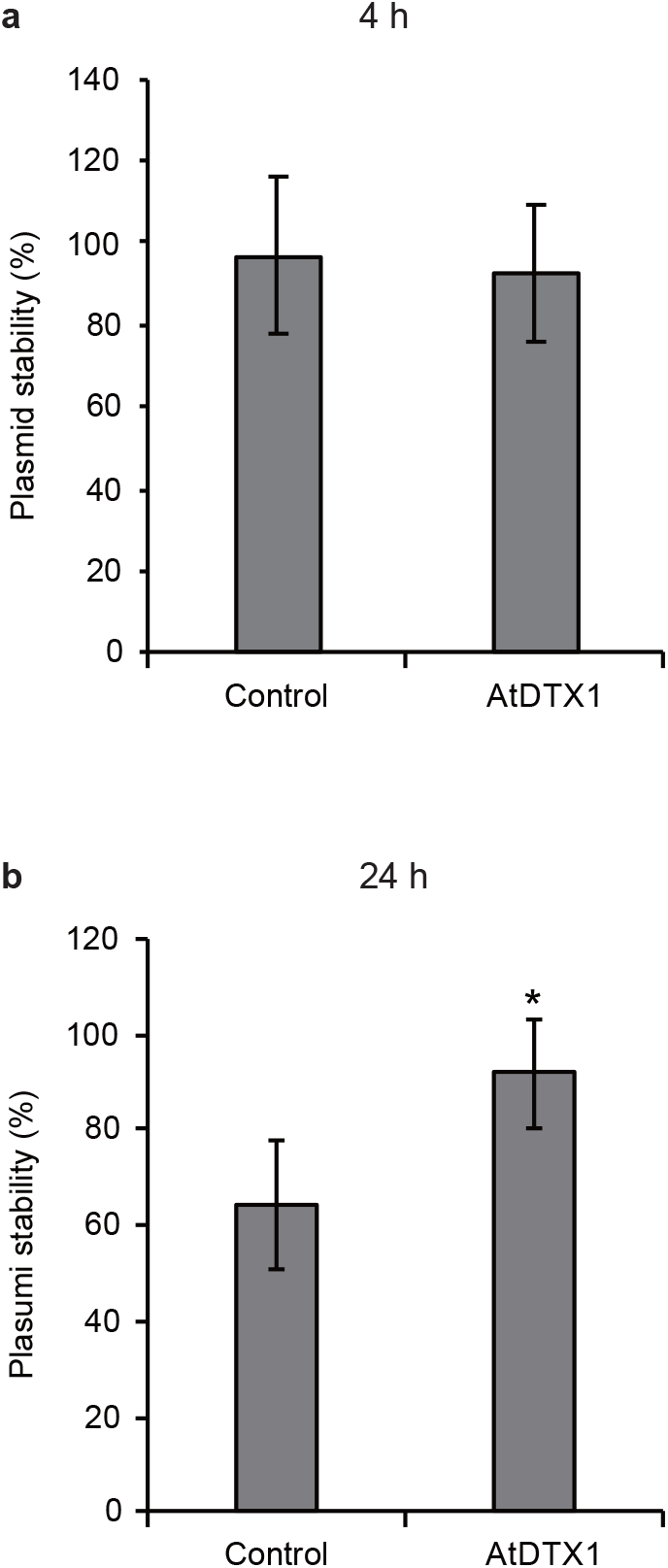
Plasmid stability of reticuline-producing *E. coli* cells. Plasmid stabilities of reticuline-producing cells harboring either control vector or pCOLADuet1_AtDTX1 grown in modified LB medium at 4, and 24 h after adding IPTG. Results show mean ± standard deviation (SD) (n = 9). Asterisks indicate statistically significant difference compared to the control (Student’s t-test; * *P* < 0.01).

Next, the changes in the transcriptome of the cells were investigated. We performed RNA-seq (Supplementary Data 1) and analyzed the differentially expressed genes (DEGs) using edgeR, which showed that compared to the vector control cells at each time point (0, 8, 12, 24, and 48 h), 1,393 genes were induced or suppressed in AtDTX1-expressing cells, with |fold change| ≥ 2 (Supplementary Data 2-7). These genes exhibited varying expression levels relative to that of the control (Supplementary Fig. 8). The number of DEGs increased in a time-dependent manner (Fig. 7a). Gene Ontology (GO) enrichment analysis indicated that several subcategories were enriched in the categories “biological process”, “cellular process”, and “molecular function”, and revealed the difference between the number of upregulated and downregulated genes (Fig. 7b-d). The number of genes that were categorized into metabolic process (GO:0008152), cellular process (GO:0009987), response to stimulus (GO:0050896), localization (GO:0051179), membrane (GO:0016020), cell part (GO: 0044464), catalytic activity (GO:0003824), transport activity (GO:0005215), and binding (GO 0005488) was high. Several genes that were categorized into detoxification (GO:0098754) and antioxidant activity (GO:0016209) were upregulated in AtDTX1-expressing cells at 12, 24, and 48 h, whereas many genes categorized into localization (GO:0051179), membrane (GO:0016020), protein-containing complex (GO:0032991), and transporter activity (GO:0005215) were downregulated (Fig. 7b-d). In addition, we also identified biological pathways regulated in AtDTX1-expressing cells by Kyoto Encyclopedia of Genes and Genomes (KEGG) analysis using the pathway of *E. coli* K-12 MG1655 (Table 1, Supplementary Table 2, and Supplementary Fig. 9, 10). In “Global and overview maps” (Table 1), the number of genes that were categorized into metabolic pathways (01100), biosynthesis of secondary metabolites (01110), microbial metabolism in diverse environments (01120), carbon metabolism (01200), biosynthesis of amino acids (01230), and biosynthesis of cofactors (01240) was high. In contrast, the number of genes related to 2-oxocarboxylic acid metabolism (01210), fatty acid metabolism (01212), and degradation of aromatic compounds (01220) was low. In detailed categories, many genes in carbohydrate metabolism, energy metabolism, nucleotide metabolism, amino acid metabolism, and membrane transport were altered (Supplementary Table 2). Interestingly, at 12 h, the time of maximum difference of cellular reticuline production was observed; several genes of pentose phosphate pathway (00030) were upregulated (Supplementary Table 2, and Supplementary Fig. 9). This pathway is involved in reticuline production by supplying various metabolites for reticuline, such as D-glyceraldehyde 3-phosphate (G3P), D-ribose 5-phosphate (R5P), and D-erythrose 4-phosphate (E4P) (Fig. 1). This induction may be caused by AtDTX1-dependent secretion of reticuline, the end-product, from the cell. Furthermore, methionine biosynthesis from homoserine was highly upregulated (Supplementary Fig. 9). The enhancement of methionine biosynthesis is also supported by the increased biosynthesis of 5-methyl-tetrahydrofolate (5-Methyl-THF), a methyl donor for methionine synthesis via homocysteine methyltransferase (Supplementary Fig. 9). Methionine is used for the biosynthesis of S-adenosyl-L-methionine (SAM), methyl donor for reticuline biosynthesis via three methyl transferases, that is, 4’OMT, CNMT, and 6OMT (Fig. 1, 8). Some methyl transferases, such as 4’OMT and 6OMT, have been reported to be inhibited by end-product or related compounds—berberine or norreticuline ^36,37^. Therefore, reticuline efflux from the cytosol via AtDTX1 may have relieved the negative feedback on methyltransferases and methionine biosynthesis was enhanced to supply sufficient amount of SAM.

**Table 1.** The number of genes induced or suppressed in KEGG pathway of “Global and overview maps” in AtDTX1-expressing cells.

| Metabolism |  | (h) | 0 | 8 | 12 | 24 | 48 |
| --- | --- | --- | --- | --- | --- | --- | --- |
| Global and overview maps | 01100 Metabolic pathways | Up | 1 | 23 | 67 | 57 | 51 |
|  |  | down | 6 | 8 | 32 | 75 | 104 |
|  | 01110 Biosynthesis of secondary metabolites | Up | 0 | 9 | 32 | 18 | 15 |
|  |  | down | 3 | 0 | 13 | 36 | 45 |
|  | 01120 Microbial metabolism in diverse environments | Up | 0 | 6 | 19 | 10 | 12 |
|  |  | down | 2 | 5 | 14 | 26 | 25 |
|  | 01200 Carbon metabolism | Up | 0 | 7 | 15 | 4 | 5 |
|  |  | down | 0 | 0 | 7 | 8 | 8 |
|  | 01210 2-Oxocarboxylic acid metabolism | Up | 0 | 2 | 3 | 0 | 0 |
|  |  | down | 0 | 0 | 0 | 0 | 2 |
|  | 01212 Fatty acid metabolism | Up | 0 | 0 | 0 | 0 | 0 |
|  |  | down | 0 | 0 | 0 | 2 | 1 |
|  | 01230 Biosynthesis of amino acids | Up | 0 | 5 | 15 | 6 | 5 |
|  |  | down | 0 | 0 | 0 | 14 | 20 |
|  | 01240 Biosynthesis of cofactors | Up | 0 | 0 | 6 | 15 | 8 |
|  |  | down | 0 | 1 | 4 | 10 | 6 |
|  | 01220 Degradation of aromatic compounds | Up | 0 | 1 | 2 | 0 | 2 |
|  |  | down | 0 | 0 | 1 | 4 | 1 |

**Fig. 7.**
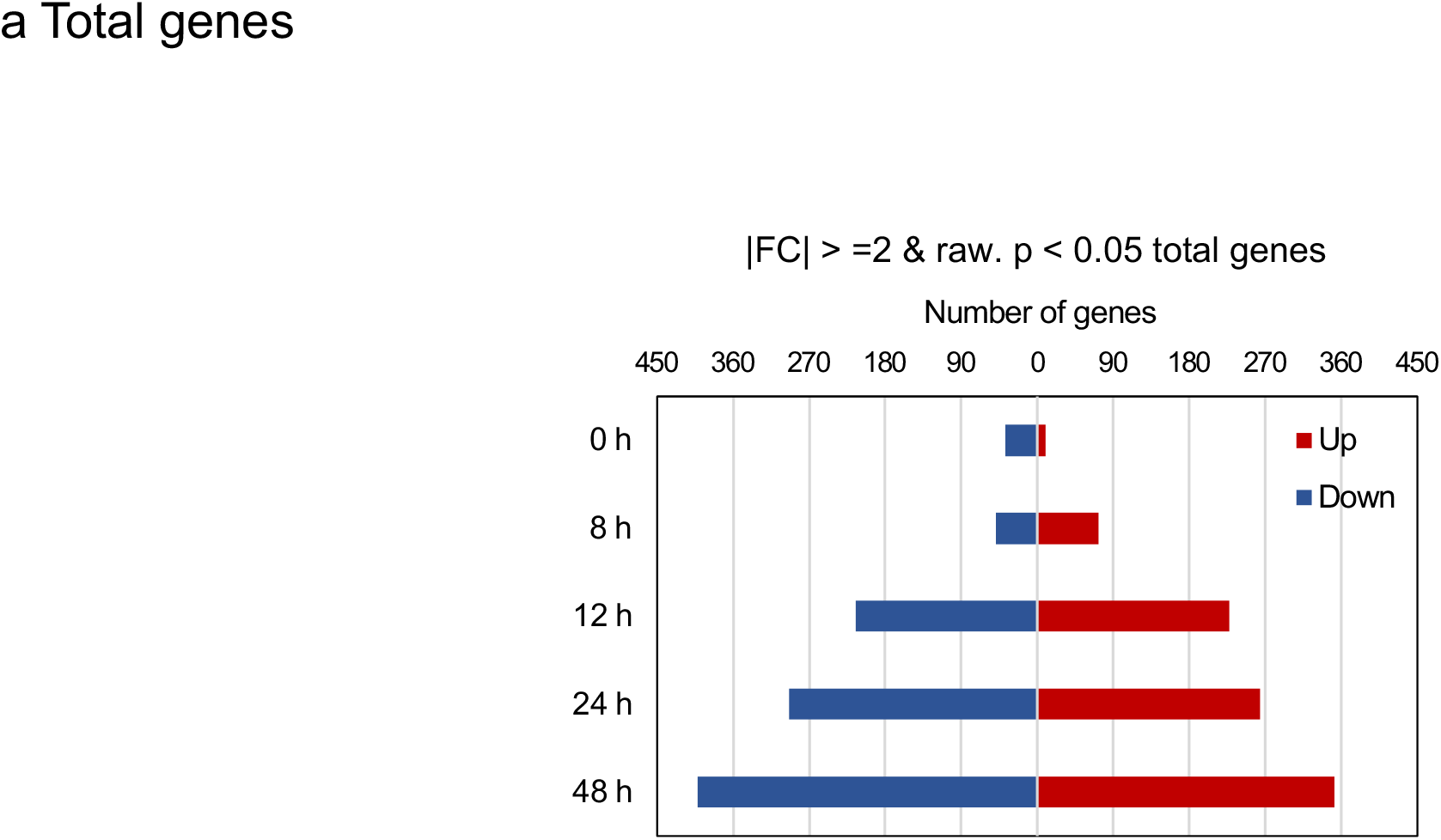
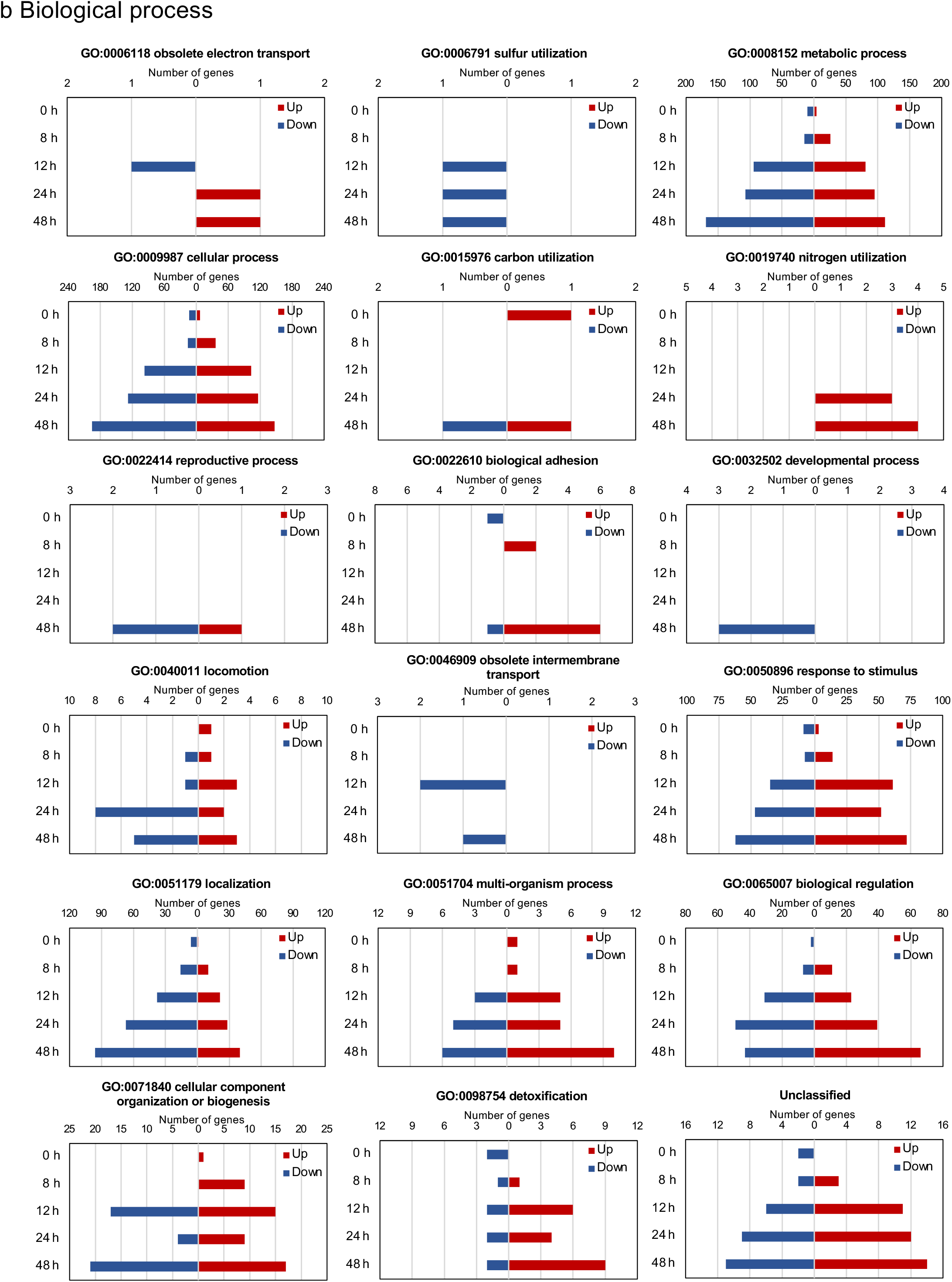
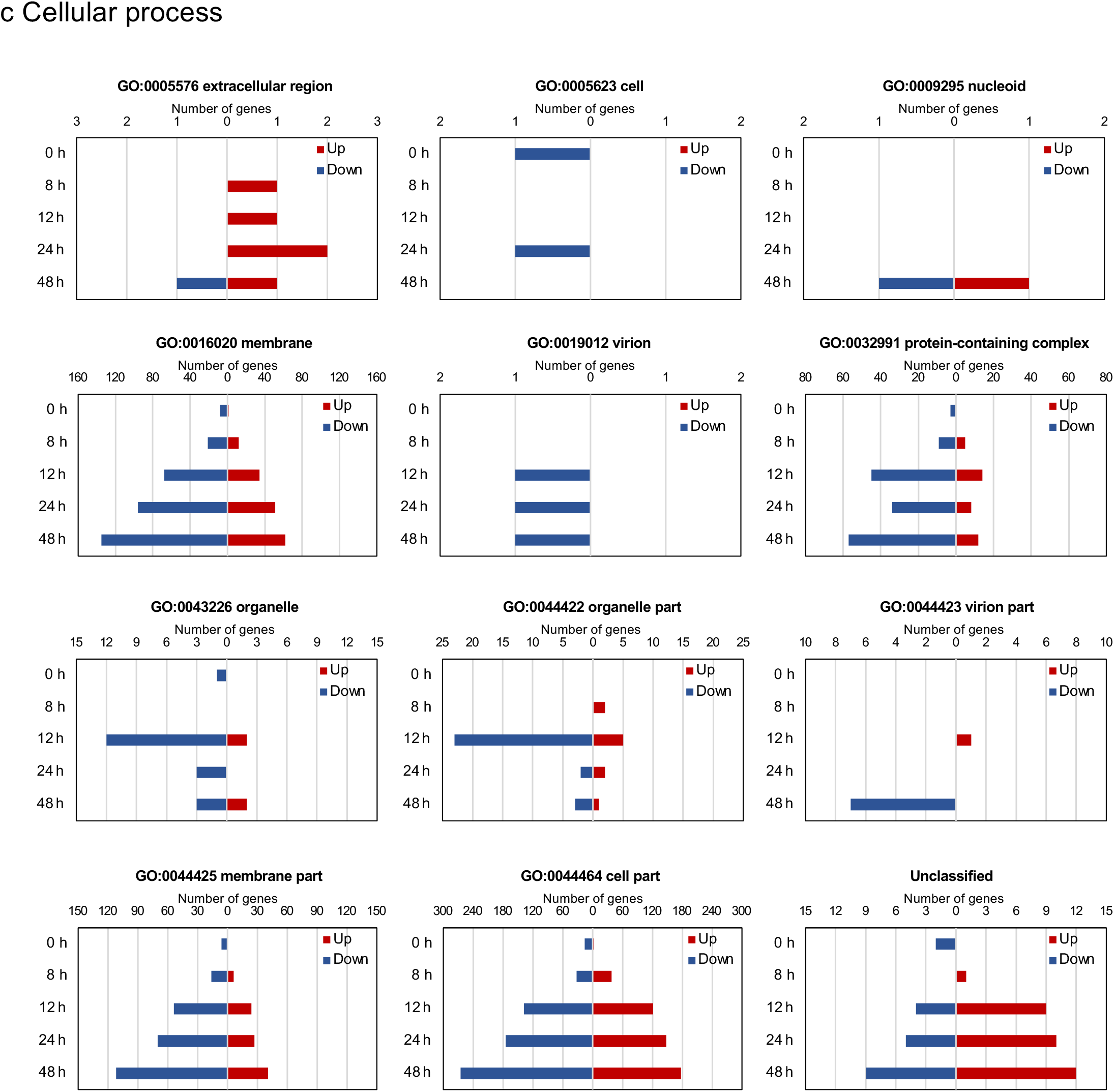
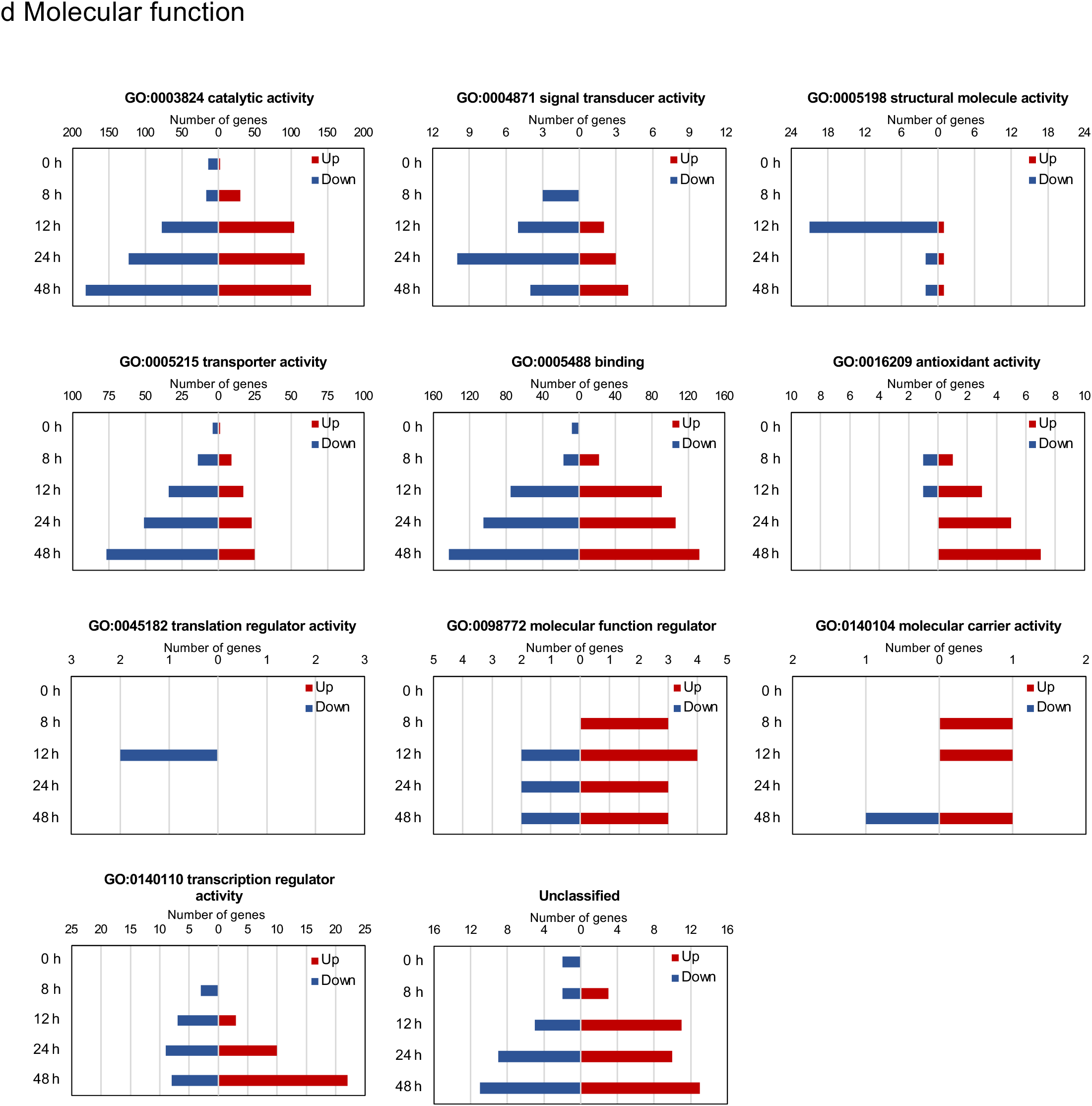
Gene Ontology categories of differentially expressed genes (DEGs) in AtDTX1-expressing cells compared to those in vector control cells at the same time points. DEGs were categorized into total (a), biological process (b), cellular process (c) and molecular function (d) based on |fc| ≥ 2 & exactTest raw *P*-value < 0.05. The number of upregulated (red bars) and downregulated (blue bars) genes are shown separately.

These data suggested that AtDTX1 expression conferred high plasmid stability in the cells, significantly affected cellular processes, and induced the biosynthetic flow in the cell (Fig. 8). Overall, these changes led to the production of high amounts of reticuline and its efflux into the medium.

**Fig. 8.**
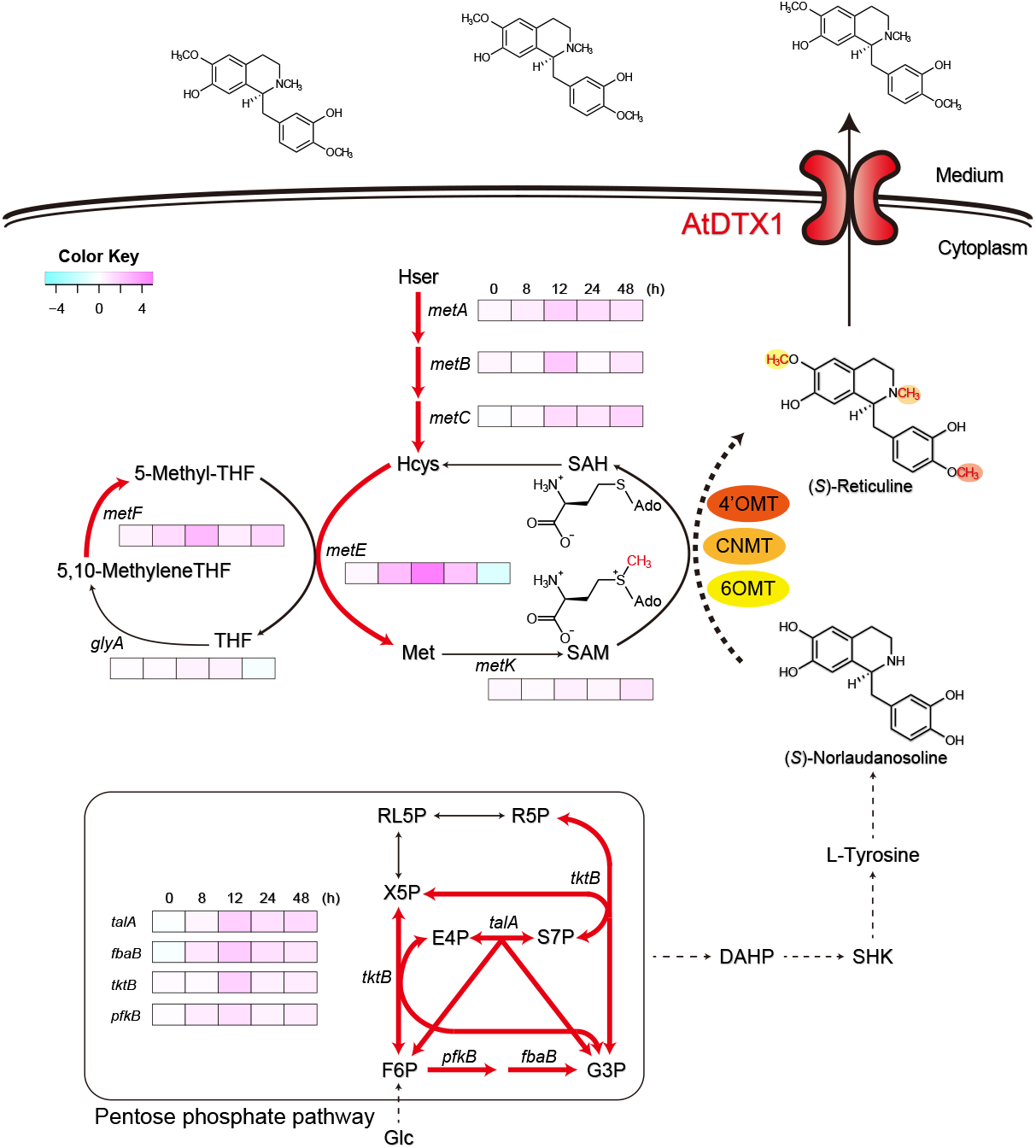
Expression pattern of endogenous genes involved in reticuline biosynthesis. Summary of gene expression changes in AtDTX1-expressing *E. coli* cells. Red arrows indicate reaction steps involved in reticuline biosynthetic genes that were up-regulated in AtDTX1-expressing cells with fold change ≥ 2 at 12 h. Heatmap (logFC) shows the expression pattern. Dotted lines represent multiple steps. Abbreviations are as follows: *fba*B fructose-bisphosphate aldolase class 1; *glyA* serine hydroxymethyltransferase; Hcys, homocysteine; Hser, homoserine; Met, methionine; *metA* homoserine transsuccinylase; *metB* cystathionine gamma-synthase; *metC* Cystathionine β-lyase; *metE* cobalamin-independent homocysteine transmethylase; metF 5,10-methylenetetrahydrofolate reductase; metK methionine adenosyltransferase; pfkB ATP-dependent 6-phosphofructokinase isozyme 2; SAH, S-adenosylhomocysteine; SAM, S-adenosylmethionine; *talA* transaldolase A; *tktB* transketolase 2; and THF, tetrahydrofolate.

## Discussion

Reports regarding the practical application of transporters for the production of specialized metabolites are scarce, as little has been known regarding plant transporters for specialized metabolites. In this study, we aimed to utilize transport engineering for the synthesis and efflux of plant specialized metabolites from *E. coli*. To the best of our knowledge, this is the first study in which a plant alkaloid transporter was used for microbial production, with the product subsequently effluxed into the medium. Heterologous expression of an efflux transporter, AtDTX1, in reticuline-producing *E. coli* increased reticuline secretion by nearly 11-fold (Fig. 5).

Selection of an appropriate transporter is important for transport engineering. In this study, we focused on the MATE family. This is because NPF and PUP transport substrates inward as proton co-transport, which is not appropriate for studying product efflux from the cytosol to the medium (Supplementary Fig. 1). Some plant ABC transporters show efflux activity. However, most plant ABC transporters, especially those involved in alkaloid transport (ABCB-type), function as a single polypeptide consisting of two transmembrane domains (TMDs) and two nucleotide binding domains (NBDs) (called full-size) ^22^, whereas most prokaryotic ABC transporters express TMD and NBD as separate proteins, which work as multi-complexes, or as half-size homo-or hetero-dimers consisting of one TMD and one NBD ^38^. Therefore, full-size plant ABC transporters might not function efficiently in *E. coli* and knowledge required for the functional analysis of plant ABCB transporters in *E. coli* is limited. In contrast, MATE transporters efflux substrates via a proton antiport mechanism, and the structure of plant MATE transporters are similar to those of microbial MATE transporters, as both contain 12 transmembrane helixes. Therefore, we selected a MATE transporter in this study.

Plant MATE transporters efflux diverse substrates, such as citrate, hormones, and specialized metabolites, and their substrate specificities are related to the homology of their amino acid sequences to some extent. In the phylogenetic relationship analysis (Supplementary Fig. 11), many transporters in clade I transport specialized metabolites; for example, AtTT12 and *Medicago truncatula* MATE1 transport proanthocyanidin, MtMATE2 and *Vitis vinifera* AM1 transport anthocyanin, and NtJAT1 and *Coptis japonica* MATE1 transport alkaloids ^20,23^. Transporters in clade II are implicated in plant development, such as in leaf senescence; however, the substrates of these transporters are not yet known, with the exception of AtDTX50, which transports abscisic acid (ABA), an important plant hormone ^24^. AtEDS5 of clade III is localized in plastids and transports salicylic acid. Most clade IV transporters transport citrate and are involved in Fe translocation or Al^3+^ detoxification ^23^.

Although many plant transporters of specialized metabolites have recently been identified and characterized ^16,18,19^, knowledge regarding their substrate specificities and expression in microorganisms is still limited. Hence, whether a selected transporter can transport a product in microorganisms should be determined, and novel transporters for specific products have to be identified in some cases. In addition, for utilizing plant transporters for the microbial production of specialized metabolites, transporter expression and activity in the host microorganism should be assessed. In this study, AtDTX1 was selected based on the results of a previous study ^34^ demonstrating the function and alkaloid transport profiles in *E. coli*. AtDTX1 was expressed well in *E. coli* and showed transport activity for reticuline (Fig. 2 and 3). In contrast, NtJAT1 expression was weak (Fig. 2) and hindered cell growth significantly (Supplementary Fig. 4), and was hence not used further in this study. However, NtJAT1 is best suited for the transport engineering of yeast cells for producing alkaloids, as its expression and substrate specificity for alkaloids have been characterized in yeast cells ^29^.

The expression level and time required for the induction of the transporter should be carefully determined. Overexpression of the transporter, similar to that of other membrane proteins, inhibits cell growth ^13^. Thus, for *n*-butanol production using transport engineering, transporter expression was optimized to balance its toxicity while ensuring efficient production ^13^. In this study, we suppressed the basal expression level of the transporter using a low copy plasmid, pCOLADuet-1. Nevertheless, significant growth inhibition was observed when expression was induced by high concentrations of IPTG (1 mM) at the lag phase of *E. coli* (OD_600_ = 0.1) (data not shown). Therefore, low concentration of IPTG (0.1 mM) was added after the cells grew to some extent (OD_600_ = 0.6). This experimental condition negligibly affected cell growth in reticuline-producing cells (Fig. 4) and resulted in high production and secretion of reticuline (Fig. 5).

Several studies have reported using transport engineering for the production of central metabolites and some specialized metabolites such as codeine and tropane alkaloids ^10,39^; the transporters used for codeine and tropane production play roles in the transport of biosynthetic intermediates from the medium to the cytosol or from the cytosol to the vacuolar lumen. However, these studies have mainly focused on the increase in production after transporter expression, and only few studies have analyzed the genetic changes in the engineered cells. In this study, we investigated the intracellular alterations based on plasmid stability and transcriptomic analysis (Fig. 6~8, Supplementary Figure 7~10, Supplementary Data 1~7). The expression of several genes involved in metabolism, especially categorized into metabolic pathways (01100), microbial metabolism in diverse environment (01120), biosynthesis of amino acids (01230), sulfur metabolism (00920), purine metabolism (00230), pyrimidine metabolism (00240), were highly regulated. Interestingly, other cellular processes, such as ribosome (03010), ABC transporters (02010), and two-component system (02020) were also highly regulated, suggesting diverse effects of AtDTX1 expression in cellular function. It is noted that metabolic pathways relatively related to reticuline biosynthesis were altered. In addition to the pentose phosphate pathway, the metabolism of several amino acids other than methionine was affected; for instance, the genes for the biosynthesis of valine, leucine, and isoleucine, were upregulated at 12 h. The downregulation of the gene for tryptophan biosynthesis at 24 h might lead to higher production of tyrosine (Supplementary Fig. 10). AtDTX1-dependent reticuline efflux might cause the decrease of biosynthetic intermediates like G3P, R5P, E4P, and tyrosine, and SAM, methyl donor for methyltransferase, which possibly resulted in the upregulation of related cellular metabolism to compensate for the supply of those compounds (Fig. 1). These alterations would have led to not only secretion to the medium but also high production in the cells (Fig. 5).

In contrast, AtDTX1 expression seems to induce stress in the cells. In the GO analysis, many genes categorized into response to stimulus (GO:0050896), detoxification (GO:0098754), antioxidant activity (GO:0016209) were upregulated (Fig. 7). Analysis of each gene revealed that genes, such as those encoding the universal stress protein UspG (locus tag: ECD_RS02910, 4.6-fold at 24 h) and stress-induced protein YchH (ECD_RS06300, 33-fold at 24 h) were highly upregulated (Supplementary Data 6). As reticuline did not significantly inhibit the growth of *E. coli* (Supplementary Fig. 12), these stress-related genes might be induced by AtDTX1 expression. GO analysis also suggested that other genes, for example, those encoding lipopolysaccharide core heptose (II) kinase RfaY (ECD_RS18200, −15.5-fold at 24 h), zinc resistance sensor/chaperone ZraP (ECD_RS20375, 154-fold at 24 h), and iron-sulfur cluster repair protein YtfE (ECD_RS21495, 35.4-fold at 24 h), were also highly up- or downregulated in AtDTX1-producing cells (Supplementary Data 6); however, whether these genes are implicated in reticuline production remains unclear. Surprisingly, plasmid stability was high in AtDTX1-expressing cells (Fig. 6), suggesting that the percentage of reticuline-producing cells was high in the culture medium. These findings indicated that the expression of the transporter AtDTX1, in this case, induced diverse changes in the cells, and the integrated effect of these alterations increased the production of the desired compound.

Application of metabolic engineering to efflux transporters enables the high production and efficient recovery of valuable specialized metabolites. Some synthesized metabolites accumulate *in vivo*; therefore, procedures for the extraction and purification of metabolites are necessary, which renders some microbial productions commercially unfeasible. Combining transport engineering with metabolic engineering not only improves the production of valuable metabolites, but is also beneficial for the efficient recovery of the products. Although the knowledge about transporters for transport engineering is still limited and the identification of the optimal transporters for the desired metabolites would be necessary, further progress in this field will enhance the production of medicinal resources from plants.

In conclusion, we successfully developed the first *E. coli* transport engineering platform that enables high production and secretion of a valuable alkaloid. The results of the present study are of considerable significance, as *E. coli* is a standard microorganism for industrial-scale production of plant medicinal resources ^2,3,7^. The combined platform of metabolic engineering and transporter engineering described here will provide opportunities for low-cost production of various valuable alkaloids. The use of this technology for rapid mass production of useful plant metabolites will contribute to the health and welfare of society.

## Methods

### Chemicals

(S)-reticuline was purified as described previously ^33^ and was used as a standard. The other chemicals used in this study were purchased from FujiFilm Wako Pure Chemical Corporation (Osaka, Japan) or Nacalai Tesque (Kyoto, Japan).

### Construction of pCOLADuet-1_AtDTX1 and NtJAT1

Total RNA was isolated from *A. thaliana* seedlings (ecotype Columbia) with the RNeasy Plant Mini-kit (Qiagen, Hilden, Germany), according to the manufacturer’s instructions. The RNA was reverse transcribed using SuperScript III reverse transcriptase (Invitrogen, CA), followed by incubation with RNaseH (Invitrogen, CA). The full-length cDNA of *AtDTX1* (At2g04040) was amplified using reverse transcription-polymerase chain reaction (RT-PCR) from the cDNA. The primer sequences used are as follows: AtDTX1_infusion2fw, 5’-<u>ACCACAGCCAGGATCC</u>GATGGAGGAGCCATTTCTTC-3’ and AtDTX1_infusion2rv, 5’-<u>AAGCATTATGCGGCCGCTTAAGCCAATC</u>TGTTTTCAGT-3’, where the underlined sequences indicate additional sequences for in-fusion cloning. The PCR product was subcloned between the *BamHI* and *NotI* sites of the pCOLADuet-1 (Novagen, USA) multiple cloning site (MCS)-1, which contains a six-amino acid His-tag for the creation of a N-terminal fusion using the In-Fusion HD cloning kit (Clontech, USA).

The full-length cDNA of *NtJAT1* (accession no. AM991692) was amplified from the cDNA of *N. tabacum*, using RT-PCR. The primer sequences are as follows: NtJAT1_infusionfw, 5’-<u>AAGGAGATATACATATG</u>GTAGAGGAGTTGCCACAG-3’ and NtJAT1_infusionrv, 5’-<u>TTACCAGACTCGAGGGTACC</u>CTAAGATCTTCCTTCGTGTA-3’, where the underlined sequences indicate additional sequences for in-fusion cloning. The PCR product was subcloned between the *Nde*I and *Kpn*I sites of the pCOLADuet-1 (Novagen, USA) MCS-2 using the In-Fusion HD cloning kit.

### Reticuline-producing *E. coli* cells

The genes encoding for reticuline biosynthesis were cloned in plasmids, as reported previously ^33^, with different combination of genes and vectors, which are summarized in Supplementary Table 1. These plasmids were introduced into *E. coli* BL21(DE3), yielding strain AN2104, which produces reticuline from a simple carbon source (Fig. 1). pCOLADuet-1 (vector control), pCOLADuet-1_AtDTX1, or pCOLADuet-1_NtJAT1 were introduced into *E. coli* BL21(DE3) or AN2104.

### Expression of AtDTX1 or NtJAT1 in *E. coli*

BL21(DE3) cells harboring pCOLADuet-1, pCOLADuet-1_AtDTX1, or pCOLADuet-1_NtJAT1 were cultured overnight in liquid LB medium containing 50 mg/L kanamycin (Nacalai Tesque) at 30°C with shaking at 200 rpm. Overnight cultures of OD_600_ = 0.1 were inoculated in 90 mL LB medium containing 50 mg/L kanamycin and grown at 30°C. IPTG (1 mM) was added when the OD_600_ of the cultures reached 0.6 and were incubated for 3 h. Next, the cells were harvested via centrifugation, re-suspended in 0.5 mL 1× phosphate buffered saline, and disrupted via sonication. Nine cycles of 30 seconds of sonication and 30 seconds of rest on ice were performed. The mixture was centrifuged at 3,000 × g for 15 min, and the supernatant was centrifuged at 20,000 × g for 20 min to pellet the membrane proteins. The membrane proteins were then denatured, subjected to electrophoresis on 10% sodium dodecyl sulfate-polyacrylamide gel, and transferred to a Immobilon polyvinylidene difluoride membrane (Millipore, Tokyo, Japan). The membrane was treated with BlockingOne (Nacalai Tesque) and incubated with anti-His monoclonal antibody (MBL, Japan) against His-AtDTX1 or anti-NtJAT1^29^ against NtJAT1. The immunoreactive band was visualized using Chemi-Lumi One Super (Nacalai Tesque).

### Reticuline transport of AtDTX1 and NtJAT1 in *E. coli* cells

BL21(DE3) cells harboring pCOLADuet-1, pCOLADuet-1_AtDTX1, or pCOLADuet-1_NtJAT1 were cultured as described above. IPTG (1 mM) was added when OD_600_ of the cultures reached 0.6, followed by incubation for 3 h. The cells were harvested, suspended in LB medium containing 50 mg/L kanamycin, 1 mM IPTG, and 250 μM reticuline at OD_600_ = 0.7, and incubated at 30°C with shaking at 200 rpm for 6 h. Next, the cells were harvested and washed with LB medium, and the reticuline content in the cells was quantified as described below.

### Growth of *E. coli* BL21(DE3) cells expressing AtDTX1 or NtJAT1

*E. coli* BL21(DE3) cells harboring either pCOLADuet-1, pCOLADuet-1_AtDTX1, or pCOLADuet-1_NtJAT1 were cultured overnight in liquid LB medium containing kanamycin at 30°C with shaking at 200 rpm. The overnight cultures were inoculated in 80 mL LB medium containing antibiotics as described above and grown at 25°C. When the OD_600_ of the cultures reached approximately 0.6, each culture was divided into three 20-mL cultures in 100-mL baffled shake flasks, followed by the addition of IPTG (0.1 mM final concentration) for induction; the samples were further cultured at 25°C with shaking at 200 rpm. Growth was evaluated by the measurement of the optical density at 600 nm.

### Growth of *E. coli* reticuline-producing cells expressing AtDTX1

Reticuline-producing *E. coli* harboring pCOLADuet-1 or pCOLADuet-1_AtDTX1 were cultured overnight in liquid LB medium containing 80 mg/L ampicillin (Sigma-Aldrich, St. Louis, MO, USA), 2 mg/L tetracycline (Nacalai Tesque), 100 mg/L spectinomycin (Nacalai Tesque), 30 mg/L chloramphenicol (Nacalai Tesque), and 50 mg/L kanamycin at 30°C with shaking at 200 rpm. The overnight grown cultures were treated as described above. Growth was evaluated by measurement of the optical density at 600 nm.

### Reticuline production

Reticuline-producing *E. coli* cells (AN2104 strain) harboring pCOLADuet-1 empty vector or pCOLADuet-1_AtDTX1 were cultured overnight in liquid LB medium containing antibiotics (tetracycline, spectinomycin, chloramphenicol, ampicillin, and kanamycin) at 30°C with shaking at 200 rpm. The overnight cultures were inoculated into 150 mL LB medium containing 9.4 g/L K_2_HPO_4_, 2.2 g/L KH_2_PO_4_, 0.4% glycerol, and antibiotics as described above, and grown at 30°C. Each culture was divided into three 45-mL cultures in 300-mL baffled shake flasks when the OD_600_ of the cultures reached approximately 0.6, following which 5 mL of 30% glucose (3% final concentration) and IPTG (0.1 mM final concentration) were added for induction; the samples were further cultured at 25°C with shaking at 150 rpm. The samples were harvested 8, 12, 24, 48, and 72 h after induction.

### UPLC-MS analysis for detection and quantification of reticuline

The culture samples were separated into supernatants (medium) and pellets (cell) via centrifugation. Trichloroacetate (2% final concentration) was added to the supernatants for precipitating the proteins, following which the supernatants were analyzed using an ACQUITY UPLC system with QDa mass detector (Waters Corp., Milford, MA, USA) after filtering using a 0.45 μm cosmospin filter (Nacalai Tesque). The pellets were incubated overnight with 15 μL/mg fresh weight (FW) methanol containing 0.1 N HCl. After centrifugation and filtration, the supernatants were also analyzed.

UPLC was performed using a CORTECS UPLC C18 column (2.1 × 100 mm, 1.6 μm; Waters Corp.) and operated at 40°C. The mobile phase A consisted of an aqueous solution of 0.01% acetic acid, while mobile phase B consisted of acetonitrile containing 0.01% acetic acid. Gradient elution was performed as follows: 0–9 min, 5–40% B; 9–12 min, 40–50% B; 12–15 min, 50–5% B. The flow rate and injection volume were set at 0.3 mL/min and 2 μL, respectively.

The QDa conditions were set as follows: cone voltage of 15 V, capillary voltage of 0.8 kV, and source temperature of 600°C. Reticuline (m/z = 330) was detected using single ion recording mode and identified by directly comparing its retention time and fragmentation spectrum (50 V cone voltage) with that of pure reticuline. The amount of reticuline was quantified using a standard curve.

### Plasmid stability

Cell viability was estimated by counting the number of colony forming units (CFU). Reticuline-producing *E. coli* cultures at 0, 4, and 24 h after IPTG treatment were diluted 100,000-fold, 1,000,000-fold, and 10,000-fold, respectively, following which 50 μL of the diluted samples were plated on LB plates containing five antibiotics or on antibiotic-free LB plates. CFUs were determined after 24–36 h of growth at 25℃. Plasmid stability was calculated by comparing the CFU on antibiotic-free plates with the CFU on antibiotic-containing plates.

### RNA-seq analysis

Reticuline-producing *E. coli* cells were harvested at 0, 8, 12, 24, and 48 h after IPTG treatment and total RNA was extracted from *E. coli* cells using an RNeasy mini kit (Qiagen) after lysis with lysozyme (FujiFilm Wako) and proteinase K (Invitrogen, Carlsbad, CA, USA). RNA sequencing and DEG analysis was carried out by Macrogen Japan Corp. (Kyoto, Japan) as follows; RNA sequencing with paired-end 101-bp reads was performed using an Illumina NovaSeq 6000 system. The quality check of the raw sequences and trimming were performed using FastQC (http://www.bioinformatics.babraham.ac.uk/projects/fastqc/) and Trimmomatic 0.38 (http://www.usadellab.org/cms/?page=trimmomatic), respectively. After the read mapping using GCF_000022665.1_ASM2266v1 as a reference genome and Bowtie 1.1.2 (http://bowtie-bio.sourceforge.net/index.shtml), expression profiling was performed using the HTSeq version 0.10.0 (http://www-huber.embl.de/users/anders/HTSeq/doc/overview.html). DEG analysis was performed on comparison pairs (AtDTX1 vs VC1) using edgeR as per the following workflow; 1) read count value of known genes obtained through the HTseq were used as the original raw data. 2) Low quality transcripts were filtered and TMM Normalization was performed during data preprocessing. 3) Statistical analysis was performed using Fold Change, exactTest using edgeR per comparison pair. The significant results were selected on the conditions of |fc|>=2 & exactTest raw *p*-value<0.05. KEGG pathway mapping was carried out using KEGG Mapper (https://www.genome.jp/kegg/mapper.html). Hierarchical clustering and heat map generation was performed using R, version 3.4.4. ^40^

### Statistical analysis

Student’s *t*-test (two-tailed) was used to determine significant differences compared to the control cells in reticuline production, and plasmid stability analyses. Multiple comparisons were conducted using repeated analysis of variance with Bonferroni test in reticuline transport.

### Data availability statement

Data supporting the findings of this work are available within the paper and its Supplementary Information files. RNA-seq data are available in the DDBJ Sequenced Read Archive under the accession number DRA011247. All relevant data presented in this paper are available from the corresponding author upon request.

## Acknowledgments

We thank Ms. Yoko Nakahara (Kobe Pharmaceutical University, Japan) for the assistance with the experiments. We thank Dr. Y. Moriyama (Kurume University, Japan) for providing the anti-NtJAT1 antibodies. DNA sequences were analyzed by the Life Research Support Center of Akita Prefectural University. This work was supported by JSPS KAKENHI (grant number 17H05453 to N.S.) Grant-in-Aid for Scientific Research on Innovative Areas.

## Author contributions

Y.Y., A.N., F.S., H.M., and N.S. designed the experiments. Y.Y., M.U., H.O., K.I., H.M., Y.I., A.N., and N.S. performed the experiments. Y.Y., F.S., A.N., H.M., and N.S. analyzed the results and wrote the manuscript. All authors read and approved the final manuscript.

