## Supplemental Tables for "Transport engineering for improving production and secretion of valuable alkaloids in *Escherichia coli*"

**Supplementary Table 1. Plasmids used in this study**

| Plasmids | Description | Source |
|---|---|---|
| pCOLADuet-1 | Kanamycin resistance, expression vector | Novagen |
| pMW118tet | Tetracycline resistance, modified expression vector, pMW118 | This study |
| pMW-TyrOE | *tyrA*$^{fbr}$, *aroG*$^{fbr}$, *tktA*, and *ppsA* in pMW118tet | This study |
| pCDF-MPSdTH2op | *BsMtrAop*, *RnPTPSop*, *RnSPRop*, and *dTH2op* in pCDFPL | Matsumura et al., 2018 |
| pET-NMop | *NCSop* and *MAOop* in pET-23a | Matsumura et al., 2018 |
| pACYC-3MT-DDC | *6OMT*, *CNMT*, *DODC*, and *4'OMT* in pACYC184 | Matsumura et al., 2018 |
| pCOLA Duet-1 _AtDTX1 | *AtDTX1* in pCOLADuet-1 | This study |
| pCOLA Duet-1 _NtJAT1 | *NtJAT1* in pCOLADuet-1 | This study |

op: codon optimized

**Supplementary Table 2. The number of differentially expressed genes in KEGG pathway in AtDTX1-expressing cells**

| Metabolism | | (h) | 0 | 8 | 12 | 24 | 48 |
|---|---|---|---|---|---|---|---|
| Carbohydrate metabolism | 00010 Glycolysis / Gluconeogenesis | up | 0 | 2 | 8 | 4 | 1 |
| | | down | 0 | 0 | 2 | 4 | 3 |
| | 00020 Citrate cycle (TCA cycle) | up | 0 | 1 | 3 | 1 | 0 |
| | | down | 1 | 0 | 1 | 1 | 5 |
| | 00030 Pentose phosphate pathway | up | 0 | 0 | 6 | 2 | 1 |
| | | down | 0 | 1 | 0 | 3 | 1 |
| | 00040 Pentose and glucuronate interconversions | up | 0 | 0 | 2 | 2 | 2 |
| | | down | 0 | 2 | 0 | 3 | 1 |
| | 00051 Fructose and mannose metabolism | up | 1 | 0 | 5 | 3 | 1 |
| | | down | 1 | 1 | 3 | 6 | 11 |
| | 00052 Galactose metabolism | up | 0 | 4 | 3 | 3 | 0 |
| | | down | 1 | 0 | 1 | 3 | 8 |
| | 00053 Ascorbate and aldarate metabolism | up | 0 | 0 | 0 | 0 | 0 |
| | | down | 0 | 1 | 0 | 3 | 1 |
| | 00500 Starch and sucrose metabolism | up | 0 | 0 | 5 | 2 | 2 |
| | | down | 0 | 0 | 0 | 1 | 3 |
| | 00520 Amino sugar and nucleotide sugar metabolism | up | 1 | 0 | 2 | 4 | 2 |
| | | down | 1 | 0 | 5 | 5 | 11 |
| | 00620 Pyruvate metabolism | up | 0 | 1 | 5 | 3 | 2 |
| | | down | 1 | 0 | 1 | 2 | 0 |
| | 00630 Glyoxylate and dicarboxylate metabolism | up | 0 | 3 | 5 | 1 | 3 |
| | | down | 1 | 0 | 4 | 3 | 0 |
| | 00640 Propanoate metabolism | up | 0 | 1 | 1 | 1 | 2 |
| | | down | 0 | 1 | 1 | 6 | 2 |
| | 00650 Butanoate metabolism | up | 0 | 3 | 4 | 1 | 1 |
| | | down | 0 | 1 | 1 | 4 | 3 |
| | 00660 C5-Branched dibasic acid metabolism | up | 0 | 1 | 2 | 0 | 0 |
| | | down | 0 | 0 | 0 | 0 | 4 |
| | 00562 Inositol phosphate metabolism | up | 0 | 0 | 0 | 0 | 0 |
| | | down | 0 | 0 | 0 | 0 | 1 |
| Energy metabolism | 00190 Oxidative phosphorylation | up | 0 | 0 | 1 | 0 | 0 |

| | | | | | | | |
|---|---|---|---|---|---|---|---|
| | | down | 0 | 0 | 0 | 1 | 5 |
| | 00680 Methane metabolism | up | 0 | 1 | 4 | 1 | 2 |
| | | down | 0 | 1 | 6 | 7 | 2 |
| | 00910 Nitrogen metabolism | up | 0 | 1 | 1 | 2 | 3 |
| | | down | 0 | 2 | 4 | 4 | 4 |
| | 00920 Sulfur metabolism | up | 0 | 0 | 4 | 3 | 1 |
| | | down | 4 | 0 | 0 | 2 | 16 |
| Lipid metabolism | 00061 Fatty acid biosynthesis | up | 0 | 0 | 0 | 0 | 0 |
| | | down | 0 | 0 | 0 | 0 | 1 |
| | 00071 Fatty acid degradation | up | 0 | 0 | 2 | 0 | 1 |
| | | down | 0 | 0 | 1 | 3 | 0 |
| | 00121 Secondary bile acid biosynthesis | up | 0 | 0 | 1 | 0 | 0 |
| | | down | 0 | 0 | 0 | 0 | 0 |
| | 00561 Glycerolipid metabolism | up | 0 | 0 | 4 | 3 | 1 |
| | | down | 0 | 0 | 1 | 1 | 2 |
| | 00564 Glycerophospholipid metabolism | up | 0 | 0 | 1 | 1 | 1 |
| | | down | 0 | 0 | 2 | 2 | 3 |
| | 00565 Ether lipid metabolism | up | 0 | 0 | 0 | 0 | 0 |
| | | down | 0 | 0 | 0 | 0 | 0 |
| | 00600 Sphingolipid metabolism | up | 0 | 0 | 0 | 0 | 0 |
| | | down | 1 | 0 | 0 | 1 | 2 |
| | 00590 Arachidonic acid metabolism | up | 0 | 0 | 0 | 0 | 0 |
| | | down | 0 | 0 | 0 | 0 | 0 |
| | 00592 alpha-Linolenic acid metabolism | up | 0 | 0 | 0 | 0 | 0 |
| | | down | 0 | 0 | 0 | 0 | 0 |
| | 01040 Biosynthesis of unsaturated fatty acids | up | 0 | 0 | 1 | 0 | 2 |
| | | down | 1 | 0 | 1 | 0 | 0 |
| Nucleotide metabolism | 00230 Purine metabolism | up | 0 | 0 | 3 | 3 | 6 |
| | | down | 1 | 1 | 4 | 8 | 7 |
| | 00240 Pyrimidine metabolism | up | 0 | 1 | 4 | 14 | 4 |
| | | down | 1 | 0 | 4 | 1 | 4 |
| Amino acid metabolism | 00250 Alanine, aspartate and glutamate metabolism | up | 0 | 1 | 1 | 4 | 4 |
| | | down | 0 | 0 | 2 | 6 | 2 |
| | 00260 Glycine, serine and threonine metabolism | up | 0 | 5 | 5 | 1 | 1 |
| | | down | 0 | 0 | 1 | 3 | 7 |

| | | | | | | | |
|---|---|---|---|---|---|---|---|
| | 00270 Cysteine and methionine metabolism | up | 0 | 2 | 5 | 5 | 4 |
| | | down | 0 | 0 | 0 | 2 | 3 |
| | 00280 Valine, leucine and isoleucine degradation | up | 0 | 0 | 0 | 0 | 0 |
| | | down | 0 | 0 | 0 | 3 | 0 |
| | 00290 Valine, leucine and isoleucine biosynthesis | up | 0 | 2 | 3 | 0 | 0 |
| | | down | 0 | 0 | 0 | 0 | 2 |
| | 00300 Lysine biosynthesis | up | 0 | 0 | 1 | 0 | 0 |
| | | down | 0 | 0 | 0 | 0 | 1 |
| | 00310 Lysine degradation | up | 0 | 0 | 1 | 1 | 1 |
| | | down | 0 | 0 | 0 | 3 | 2 |
| | 00220 Arginine biosynthesis | up | 0 | 0 | 0 | 1 | 2 |
| | | down | 0 | 0 | 2 | 0 | 1 |
| | 00330 Arginine and proline metabolism | up | 0 | 0 | 1 | 3 | 3 |
| | | down | 0 | 0 | 0 | 0 | 0 |
| | 00340 Histidine metabolism | up | 0 | 0 | 0 | 0 | 0 |
| | | down | 0 | 0 | 0 | 8 | 5 |
| | 00350 Tyrosine metabolism | up | 0 | 0 | 2 | 0 | 1 |
| | | down | 0 | 0 | 1 | 3 | 1 |
| | 00360 Phenylalanine metabolism | up | 0 | 1 | 0 | 2 | 4 |
| | | down | 0 | 0 | 0 | 4 | 2 |
| | 00380 Tryptophan metabolism | up | 0 | 0 | 1 | 1 | 2 |
| | | down | 0 | 0 | 0 | 1 | 2 |
| | 00400 Phenylalanine, tyrosine and tryptophan biosynthesis | up | 0 | 0 | 1 | 0 | 0 |
| | | down | 0 | 0 | 0 | 2 | 8 |
| Metabolism of other amino acids | 00410 beta-Alanine metabolism | up | 0 | 1 | 0 | 0 | 0 |
| | | down | 0 | 0 | 2 | 3 | 3 |
| | 00430 Taurine and hypotaurine metabolism | up | 0 | 0 | 1 | 0 | 0 |
| | | down | 0 | 0 | 0 | 0 | 1 |
| | 00440 Phosphonate and phosphinate metabolism | up | 0 | 0 | 0 | 0 | 0 |
| | | down | 0 | 1 | 0 | 0 | 0 |
| | 00450 Selenocompound metabolism | up | 0 | 2 | 4 | 2 | 1 |
| | | down | 1 | 1 | 0 | 0 | 4 |
| | 00460 Cyanoamino acid metabolism | up | 0 | 0 | 0 | 0 | 0 |
| | | down | 0 | 0 | 0 | 1 | 0 |
| | | up | 0 | 0 | 0 | 0 | 0 |

| | | | | | | | |
|---|---|---|---|---|---|---|---|
| | 00471 D-Glutamine and D-glutamate metabolism | down | 0 | 0 | 1 | 0 | 1 |
| | 00473 D-Alanine metabolism | up | 0 | 0 | 0 | 0 | 0 |
| | | down | 0 | 0 | 0 | 0 | 0 |
| | 00480 Glutathione metabolism | up | 0 | 2 | 4 | 3 | 3 |
| | | down | 0 | 0 | 0 | 0 | 0 |
| Glycan biosynthesis and metabolism | 00540 Lipopolysaccharide biosynthesis | up | 0 | 0 | 0 | 2 | 0 |
| | | down | 0 | 0 | 3 | 3 | 6 |
| | 00541 O-Antigen nucleotide sugar biosynthesis | up | 1 | 0 | 0 | 1 | 0 |
| | | down | 1 | 0 | 3 | 3 | 2 |
| | 00550 Peptidoglycan biosynthesis | up | 0 | 0 | 0 | 0 | 0 |
| | | down | 0 | 0 | 0 | 1 | 8 |
| | 00511 Other glycan degradation | up | 0 | 0 | 0 | 0 | 0 |
| | | down | 1 | 0 | 0 | 1 | 1 |
| Metabolism of cofactors and vitamins | 00730 Thiamine metabolism | up | 0 | 0 | 0 | 0 | 1 |
| | | down | 0 | 0 | 0 | 4 | 3 |
| | 00740 Riboflavin metabolism | up | 0 | 0 | 0 | 1 | 1 |
| | | down | 0 | 0 | 0 | 0 | 1 |
| | 00750 Vitamin B6 metabolism | up | 0 | 0 | 0 | 0 | 1 |
| | | down | 0 | 0 | 0 | 0 | 1 |
| | 00760 Nicotinate and nicotinamide metabolism | up | 0 | 1 | 1 | 2 | 4 |
| | | down | 0 | 0 | 0 | 2 | 0 |
| | 00770 Pantothenate and CoA biosynthesis | up | 0 | 3 | 2 | 0 | 0 |
| | | down | 0 | 0 | 2 | 1 | 4 |
| | 00780 Biotin metabolism | up | 0 | 0 | 0 | 0 | 1 |
| | | down | 0 | 1 | 2 | 2 | 0 |
| | 00785 Lipoic acid metabolism | up | 0 | 0 | 0 | 0 | 0 |
| | | down | 0 | 0 | 0 | 0 | 0 |
| | 00790 Folate biosynthesis | up | 0 | 0 | 4 | 3 | 0 |
| | | down | 0 | 0 | 0 | 1 | 0 |
| | 00670 One carbon pool by folate | up | 0 | 2 | 2 | 0 | 0 |
| | | down | 0 | 0 | 1 | 2 | 1 |
| | 00860 Porphyrin and chlorophyll metabolism | up | 0 | 0 | 0 | 1 | 0 |
| | | down | 0 | 0 | 0 | 1 | 0 |
| | | up | 0 | 0 | 1 | 2 | 2 |

| | | | | | | | |
|---|---|---|---|---|---|---|---|
| | 00130 Ubiquinone and other terpenoid-quinone biosynthesis | down | 0 | 0 | 2 | 1 | 2 |
| Metabolism of terpenoids and polyketides | 00900 Terpenoid backbone biosynthesis | up | 0 | 0 | 0 | 0 | 0 |
| | | down | 0 | 0 | 0 | 0 | 2 |
| | 00903 Limonene and pinene degradation | up | 0 | 0 | 0 | 0 | 0 |
| | | down | 0 | 0 | 0 | 1 | 0 |
| | 00281 Geraniol degradation | up | 0 | 0 | 1 | 1 | 0 |
| | | down | 0 | 0 | 0 | 1 | 0 |
| | 00523 Polyketide sugar unit biosynthesis | up | 0 | 0 | 0 | 0 | 0 |
| | | down | 0 | 0 | 0 | 5 | 0 |
| | 01053 Biosynthesis of siderophore group nonribosomal peptides | up | 0 | 0 | 0 | 0 | 2 |
| | | down | 0 | 0 | 0 | 0 | 1 |
| Biosynthesis of other secondary metabolites | 00332 Carbapenem biosynthesis | up | 0 | 0 | 0 | 0 | 0 |
| | | down | 0 | 0 | 0 | 0 | 0 |
| | 00261 Monobactam biosynthesis | up | 0 | 0 | 0 | 0 | 0 |
| | | down | 1 | 0 | 0 | 0 | 2 |
| | 00521 Streptomycin biosynthesis | up | 0 | 0 | 0 | 0 | 0 |
| | | down | 0 | 0 | 0 | 0 | 1 |
| | 00525 Acarbose and validamycin biosynthesis | up | 0 | 0 | 0 | 0 | 0 |
| | | down | 0 | 0 | 0 | 0 | 0 |
| | 00401 Novobiocin biosynthesis | up | 0 | 0 | 0 | 0 | 0 |
| | | down | 0 | 0 | 0 | 1 | 1 |
| Xenobiotics biodegradation and metabolism | 00362 Benzoate degradation | up | 0 | 0 | 0 | 0 | 0 |
| | | down | 0 | 0 | 0 | 1 | 0 |
| | 00627 Aminobenzoate degradation | up | 0 | 0 | 0 | 0 | 0 |
| | | down | 0 | 1 | 1 | 0 | 0 |
| | 00364 Fluorobenzoate degradation | up | 0 | 0 | 0 | 0 | 0 |
| | | down | 0 | 0 | 0 | 0 | 0 |
| | 00625 Chloroalkane and chloroalkene degradation | up | 0 | 0 | 2 | 0 | 1 |
| | | down | 0 | 0 | 1 | 1 | 0 |
| | 00361 Chlorocyclohexane and chlorobenzene degradation | up | 0 | 0 | 0 | 0 | 0 |
| | | down | 0 | 0 | 0 | 0 | 0 |
| | 00623 Toluene degradation | up | 0 | 0 | 0 | 0 | 0 |
| | | down | 0 | 0 | 0 | 0 | 0 |
| | 00622 Xylene degradation | up | 0 | 0 | 0 | 0 | 0 |

| | | | | | | | |
|---|---|---|---|---|---|---|---|
| | | down | 0 | 0 | 0 | 0 | 0 |
| | 00633 Nitrotoluene degradation | up | 0 | 0 | 1 | 1 | 1 |
| | | down | 0 | 0 | 1 | 0 | 1 |
| | 00930 Caprolactam degradation | up | 0 | 0 | 0 | 0 | 0 |
| | | down | 0 | 0 | 0 | 1 | 0 |
| | 00621 Dioxin degradation | up | 0 | 0 | 0 | 0 | 0 |
| | | down | 0 | 0 | 0 | 0 | 0 |
| | 00626 Naphthalene degradation | up | 0 | 0 | 2 | 0 | 1 |
| | | down | 0 | 0 | 1 | 1 | 0 |

| **Genetic Information Processing** | | (h) | 0 | 8 | 12 | 24 | 48 |
|---|---|---|---|---|---|---|---|
| Transcription | 03020 RNA polymerase | up | 0 | 0 | 0 | 0 | 0 |
| | | down | 0 | 0 | 0 | 0 | 0 |
| Translation | 03010 Ribosome | up | 0 | 0 | 0 | 0 | 0 |
| | | down | 0 | 0 | 23 | 1 | 1 |
| | 00970 Aminoacyl-tRNA biosynthesis | up | 0 | 0 | 0 | 0 | 0 |
| | | down | 0 | 0 | 0 | 0 | 0 |
| Folding, sorting and degradation | 03060 Protein export | up | 0 | 0 | 0 | 0 | 0 |
| | | down | 0 | 0 | 0 | 1 | 3 |
| | 04122 Sulfur relay system | up | 0 | 0 | 3 | 0 | 1 |
| | | down | 0 | 0 | 1 | 1 | 2 |
| | 03018 RNA degradation | up | 0 | 0 | 0 | 0 | 0 |
| | | down | 0 | 0 | 2 | 0 | 1 |
| Replication and repair | 03030 DNA replication | up | 0 | 0 | 1 | 2 | 0 |
| | | down | 0 | 0 | 1 | 1 | 3 |
| | 03410 Base excision repair | up | 0 | 0 | 1 | 1 | 2 |
| | | down | 0 | 0 | 1 | 1 | 3 |
| | 03420 Nucleotide excision repair | up | 0 | 0 | 1 | 0 | 0 |
| | | down | 0 | 0 | 0 | 1 | 0 |
| | 03430 Mismatch repair | up | 0 | 0 | 0 | 2 | 0 |
| | | down | 0 | 0 | 1 | 1 | 3 |
| | 03440 Homologous recombination | up | 0 | 0 | 1 | 2 | 0 |
| | | down | 0 | 0 | 3 | 0 | 5 |

| **Environmental Information Processing** | | (h) | 0 | 8 | 12 | 24 | 48 |
|---|---|---|---|---|---|---|---|
| Membrane transport | 02010 ABC transporters | up | 0 | 3 | 8 | 11 | 16 |

| | | | | | | | |
|---|---|---|---|---|---|---|---|
| | | down | 1 | 10 | 12 | 25 | 30 |
| | 02060 Phosphotransferase system (PTS) | up | 0 | 3 | 5 | 6 | 0 |
| | | down | 0 | 0 | 2 | 2 | 13 |
| | 03070 Bacterial secretion system | up | 0 | 0 | 0 | 0 | 0 |
| | | down | 0 | 0 | 1 | 0 | 3 |
| Signal transduction | 02020 Two-component system | up | 0 | 1 | 3 | 15 | 15 |
| | | down | 1 | 5 | 11 | 23 | 15 |

| **Cellular Processes** | | (h) | 0 | 8 | 12 | 24 | 48 |
|---|---|---|---|---|---|---|---|
| Cellular community - prokaryotes | 02024 Quorum sensing | up | 0 | 0 | 0 | 2 | 3 |
| | | down | 0 | 2 | 4 | 3 | 8 |
| | 02026 Biofilm formation - Escherichia coli | up | 0 | 0 | 1 | 3 | 4 |
| | | down | 0 | 1 | 1 | 3 | 4 |
| Cell motility | 02030 Bacterial chemotaxis | up | 0 | 0 | 0 | 1 | 0 |
| | | down | 0 | 1 | 1 | 8 | 3 |
| | 02040 Flagellar assembly | up | 0 | 4 | 1 | 3 | 0 |
| | | down | 0 | 0 | 2 | 0 | 11 |

| **Human Diseases** | | (h) | 0 | 8 | 12 | 24 | 48 |
|---|---|---|---|---|---|---|---|
| Drug resistance: antimicrobial | 01501 beta-Lactam resistance | up | 0 | 0 | 0 | 1 | 0 |
| | | down | 0 | 0 | 1 | 0 | 3 |
| | 01502 Vancomycin resistance | up | 0 | 0 | 0 | 1 | 1 |
| | | down | 0 | 0 | 0 | 0 | 1 |
| | 01503 Cationic antimicrobial peptide (CAMP) resistance | up | 0 | 0 | 0 | 3 | 3 |
| | | down | 0 | 0 | 0 | 0 | 2 |
