## Supplementary Figures for "Transport engineering for improving production and secretion of valuable alkaloids in *Escherichia coli*"

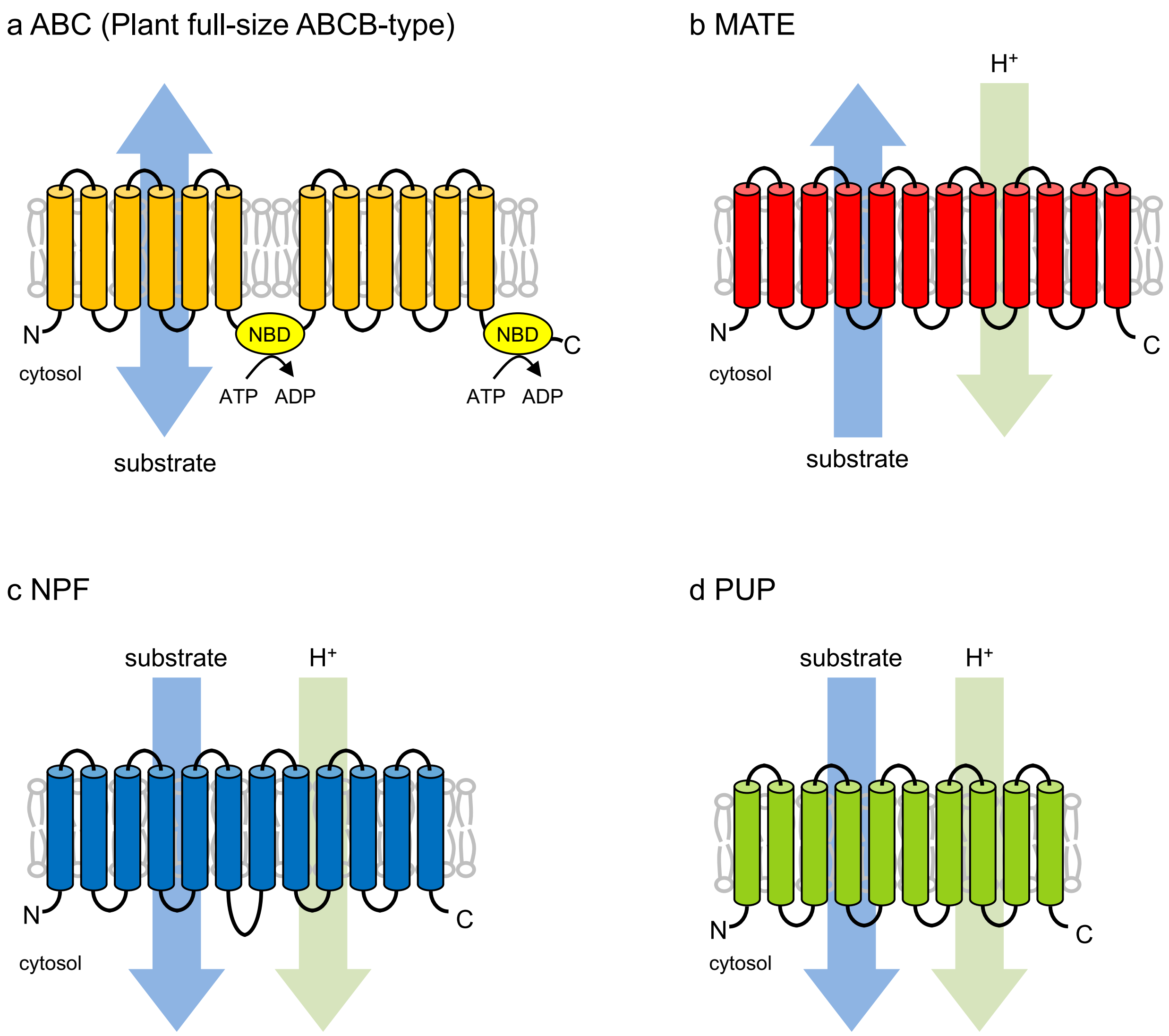


**Supplementary Fig. 1 Representative structure and the direction of transport of plant transporters involved in transporting specialized metabolites.** (a) ABC (plant full-size ABCB-type) transporter. (b) MATE transporter. (c) NPF transporter. (d) PUP transporter.

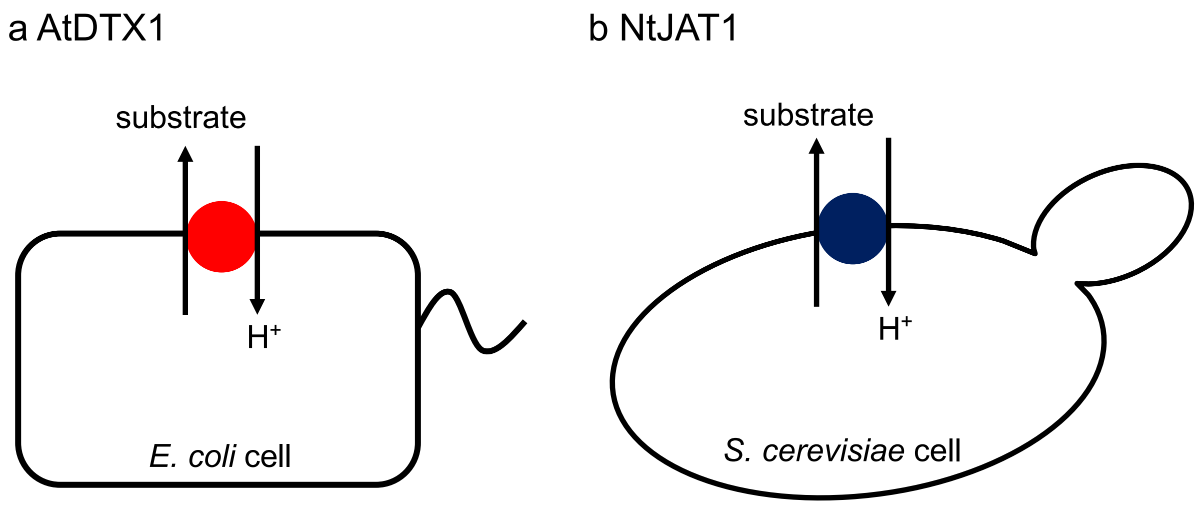


| | AtDTX1 | NtJAT1 |
|---|---|---|
| Organism | *Arabidopsis thaliana* | *Nicotiana tabacum* |
| Subcellular localization | Plasma membrane | Vacuolar membrane (*N. tabacum*)<br>Plasma membrane (yeast cells) |
| Functional characterization | Heterologous expression and transport analysis using *E. coli* cells | Heterologous expression and transport analysis using yeast cells or insect cells (proteoliposeome assay) |
| Substrates | Berberine, palmatine, norfloxacin, ethidium bromide, $Cd^{2+}$ | Nicotine, berberine, hyoscyamine, anabasine, rhodamine, ethidium bromide, verapamil |
| Reference | Li et al., *Journal of Biological Chemistry,* 277, 5360-5368 (2002) | Morita et al., *Proceedings of the National Academy of Sciences of the United States of America* 106, 2447-2452 (2009) |

**Supplementary Fig. 2 Model showing AtDTX1 (a) and NtJAT1 (b) function in microorganisms.**

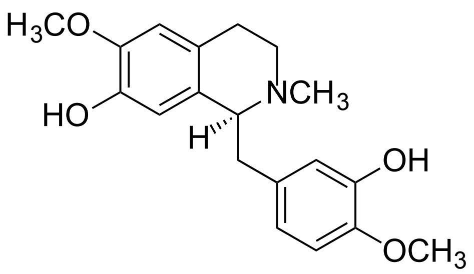


(*S*)-Reticuline

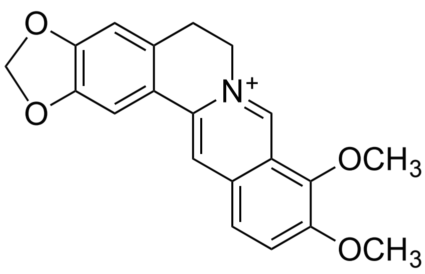


Berberine

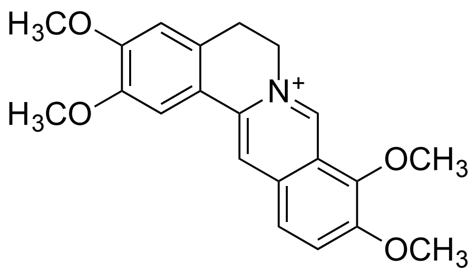


Palmatine

**Supplementary Fig. 3 Structures of reticuline, berberine, and palmatine.**

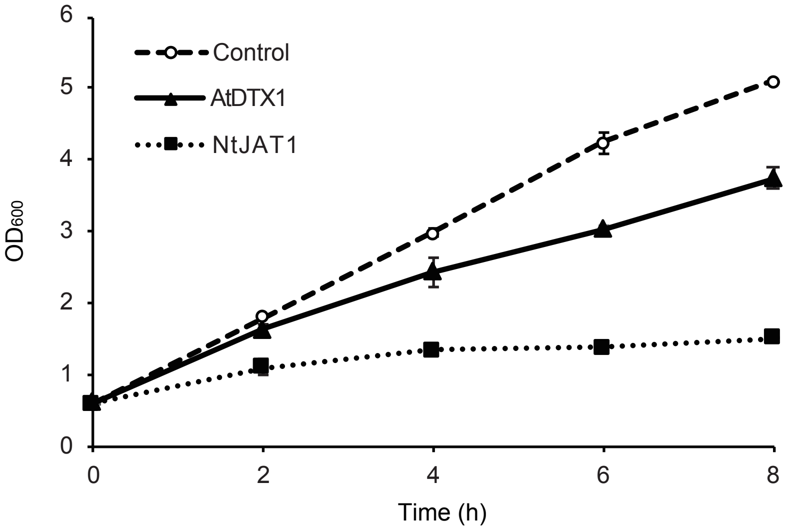


**Supplementary Fig. 4 Growth of *E. coli* BL21(DE3) cells expressing AtDTX1 or NtJAT1.** Growth was evaluated by measurement of the optical density at 600 nm.

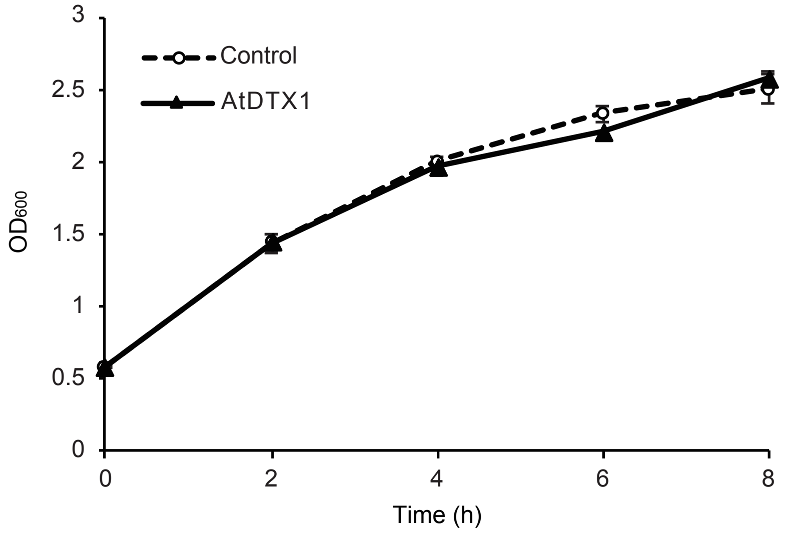


**Supplementary Fig. 5 Growth of *E. coli* reticuline-producing cells expressing AtDTX1.**

Growth was evaluated by measurement of the optical density at 600 nm.

**a Reticuline (authentic standard)** **b Reticuline (medium)**

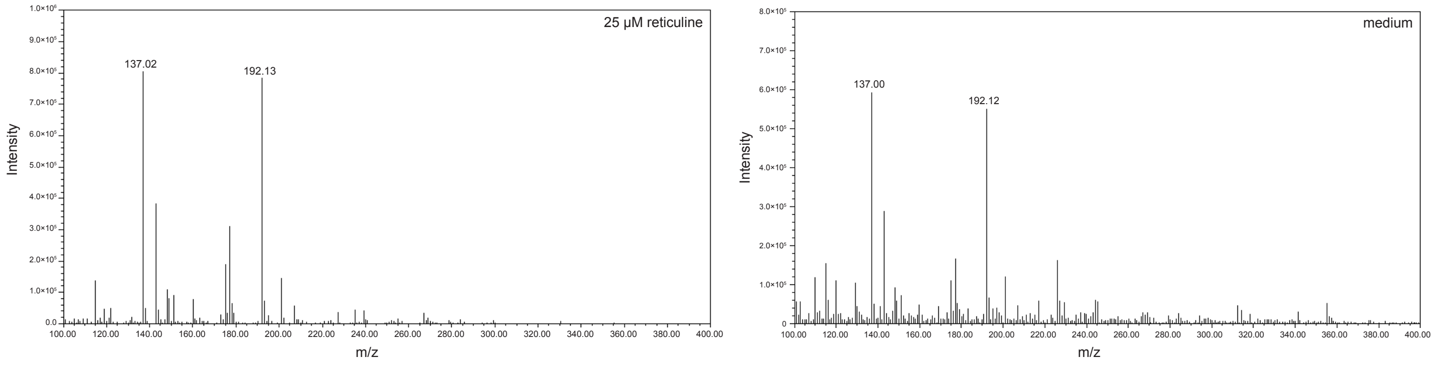


**Supplementary Fig. 6 Fragmentation patterns of authentic reticuline (a) and reticuline extracted from medium (b).**

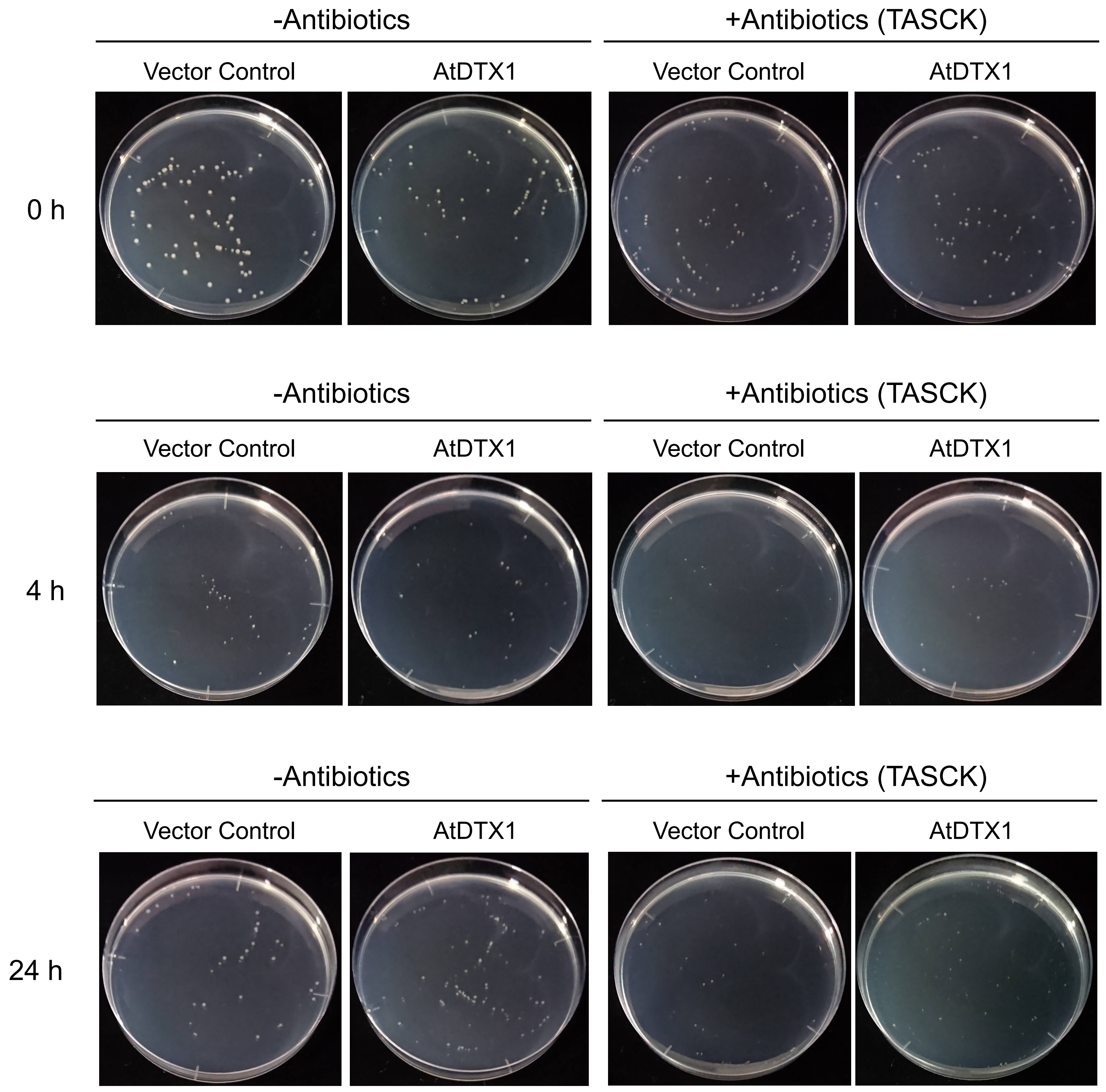


**Supplementary Fig. 7 Representative image showing colonies on LB medium with or without antibiotics.** TASCK denotes tetracycline, ampicillin, spectinomycin, chloramphenicol, and kanamycin.

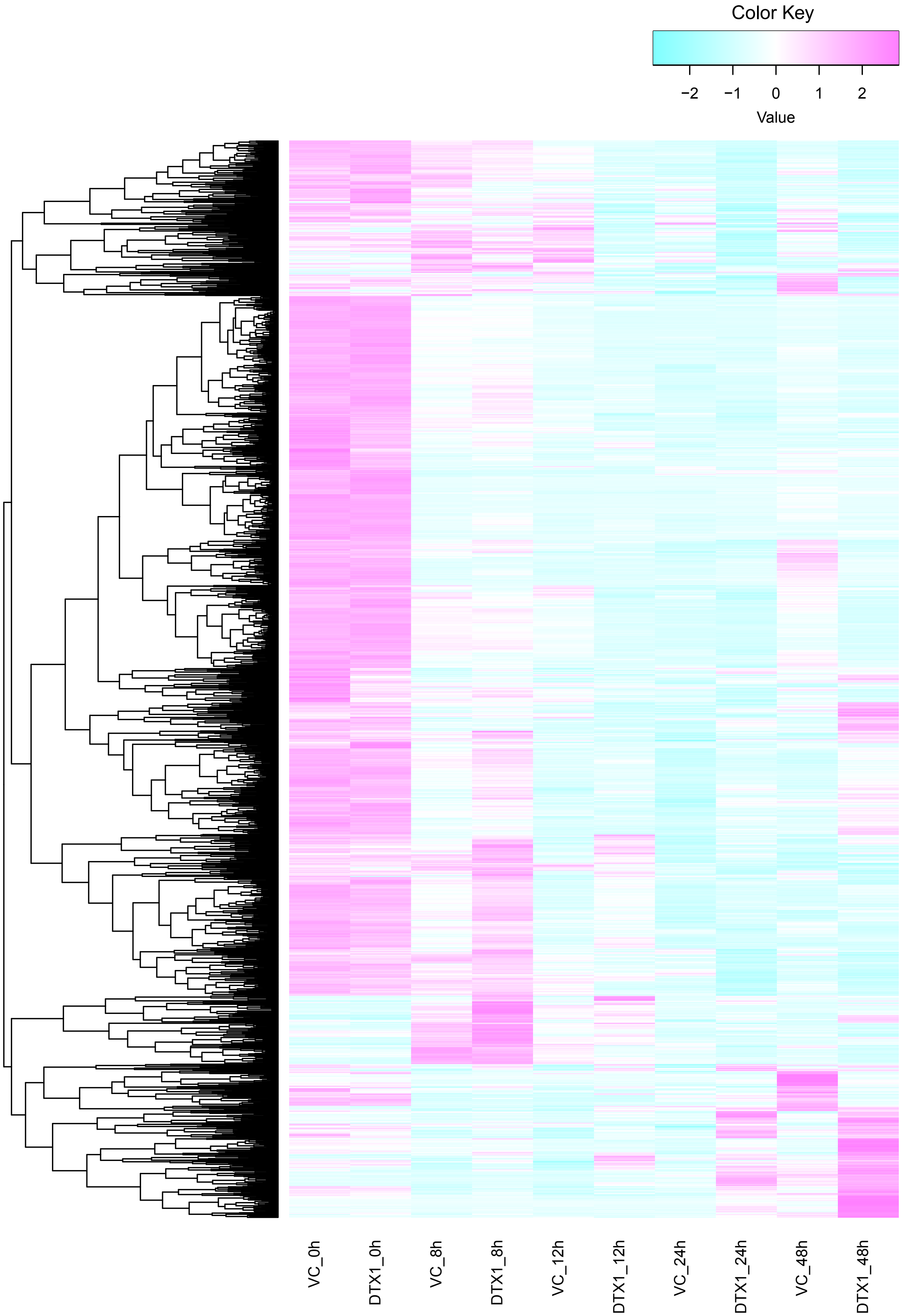

Color Key
−2
−1
0
1
2
Value
VC_0h
DTX1_0h
VC_8h
DTX1_8h
VC_12h
DTX1_12h
VC_24h
DTX1_24h
VC_48h
DTX1_48h

**Supplementary Fig. 8 Heatmap of 1,393 genes induced or suppressed in AtDTX1-expressing cells, with |fold change| ≥ 2.** Red indicates the upregulation and blue indicates the downregulation of each gene.

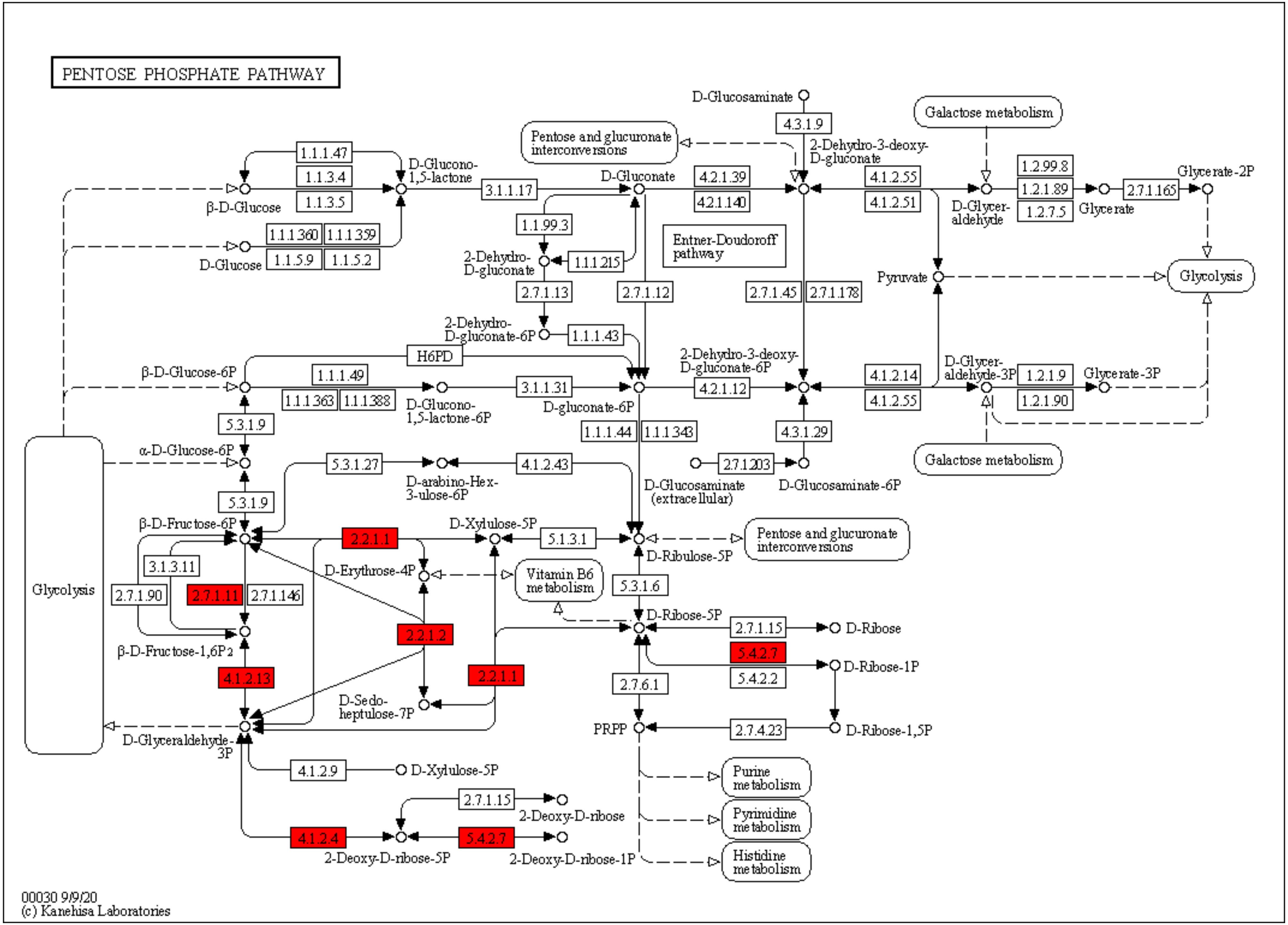
PENTOSE PHOSPHATE PATHWAY
D-Glucosaminate
4.3.1.9
2-Dehydro-3-deoxy-D-gluconate
Galactose metabolism
Pentose and glucuronate interconversions
1.1.1.47
1.1.3.4
1.1.3.5
D-Glucono-1,5-lactone
β-D-Glucose
3.1.1.17
D-Gluconate
4.2.1.39
4.2.1.140
4.1.2.55
4.1.2.51
D-Glyceraldehyde
1.2.99.8
1.2.1.89
1.2.7.5
Glycerate
2.7.1.165
Glycerate-2P
1.1.1.360
1.1.1.359
1.1.5.9
1.1.5.2
D-Glucose
1.1.99.3
2-Dehydro-D-gluconate
1.1.1.215
Entner-Doudoroff pathway
Pyruvate
Glycolysis
2.7.1.13
2.7.1.12
2.7.1.45
2.7.1.178
2-Dehydro-D-gluconate-6P
1.1.1.43
H6PD
β-D-Glucose-6P
1.1.1.49
1.1.1.363
1.1.1.388
D-Glucono-1,5-lactone-6P
3.1.1.31
D-gluconate-6P
2-Dehydro-3-deoxy-D-gluconate-6P
4.2.1.12
4.1.2.14
4.1.2.55
D-Glyceraldehyde-3P
1.2.1.9
1.2.1.90
Glycerate-3P
5.3.1.9
1.1.1.44
1.1.1.343
4.3.1.29
α-D-Glucose-6P
5.3.1.27
D-arabino-Hex-3-ulose-6P
4.1.2.43
2.7.1.203
D-Glucosamine (extracellular)
D-Glucosamine-6P
Galactose metabolism
Glycolysis
5.3.1.9
β-D-Fructose-6P
2.2.1.1
D-Xylulose-5P
5.1.3.1
D-Ribulose-5P
Pentose and glucuronate interconversions
3.1.3.11
2.7.1.90
2.7.1.11
2.7.1.146
D-Erythrose-4P
Vitamin B6 metabolism
5.3.1.6
D-Ribose-5P
2.7.1.15
D-Ribose
2.2.1.2
β-D-Fructose-1,6P2
5.4.2.7
D-Ribose-1P
5.4.2.2
4.1.2.13
2.2.1.1
2.7.6.1
D-Sedo-heptulose-7P
D-Glyceraldehyde-3P
PRPP
2.7.4.23
D-Ribose-1,5P
4.1.2.9
D-Xylulose-5P
Purine metabolism
2.7.1.15
2-Deoxy-D-ribose
Pyrimidine metabolism
4.1.2.4
5.4.2.7
2-Deoxy-D-ribose-5P
2-Deoxy-D-ribose-1P
Histidine metabolism
00030 9/9/20
(c) Kanehisa Laboratories

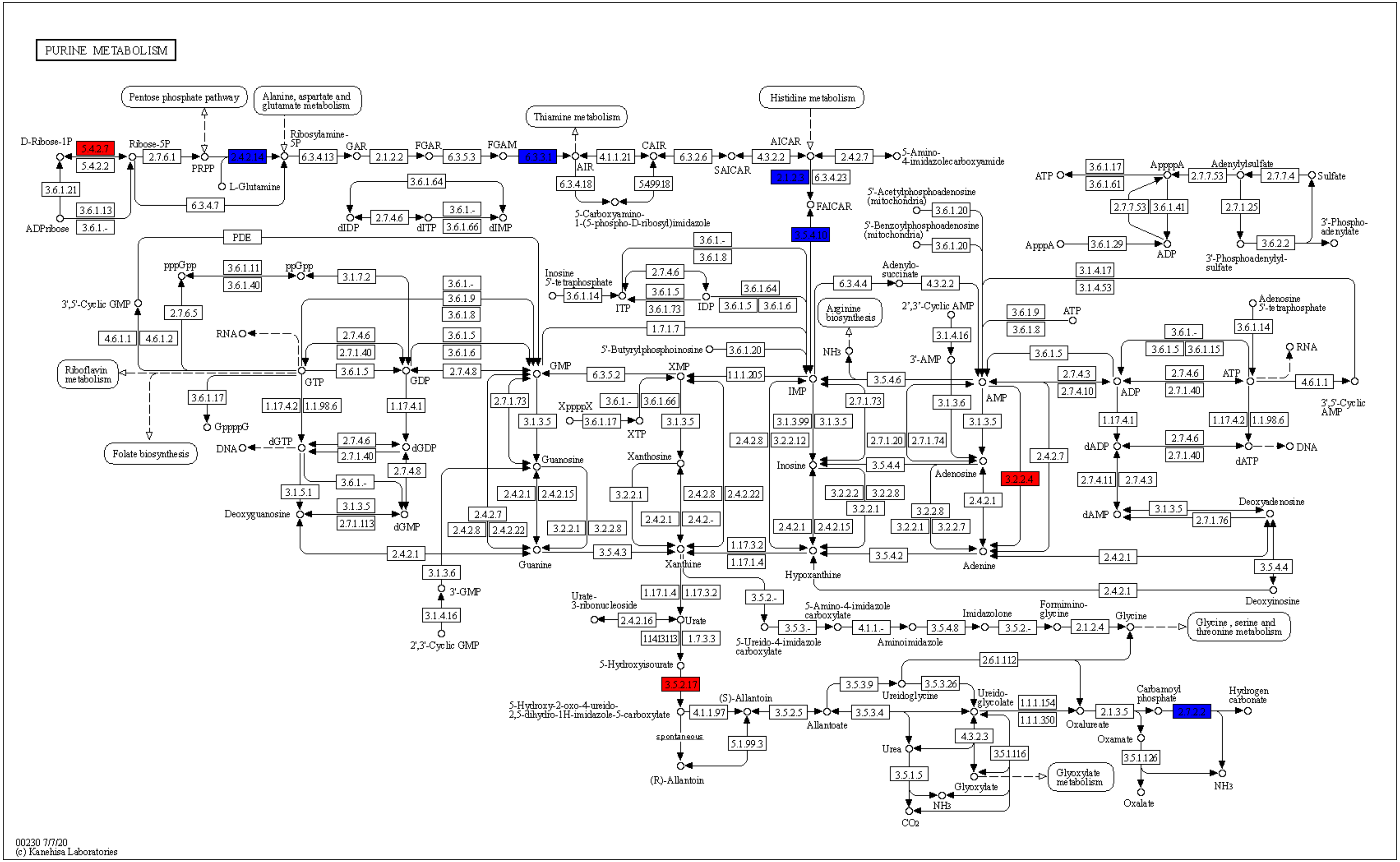

PURINE METABOLISM
Pentose phosphate pathway
Alanine, aspartate and glutamate metabolism
Thiamine metabolism
Histidine metabolism
D-Ribose-1P
5.4.2.7
5.4.2.2
Ribose-5P
2.7.6.1
PRPP
2.4.2.14
Ribosylamine-5P
L-Glutamine
6.3.4.13
GAR
2.1.2.2
FGAR
6.3.5.3
FGAM
6.3.3.1
AIR
4.1.1.21
CAIR
6.3.2.6
SAICAR
4.3.2.2
AICAR
2.4.2.7
5-Amino-4-imidazolecarboxyamide
3.6.1.21
3.6.1.13
3.6.1.-
ADPribose
6.3.4.7
3.6.1.64
2.7.4.6
dIDP
dITP
3.6.1.66
dIMP
6.3.4.18
5.4.99.18
5-Carboxyamino-1-(5-phospho-D-ribosyl)imidazole
2.1.2.3
6.3.4.23
FAICAR
3.5.4.10
5'-Acetylphosphoadenosine (mitochondria)
5'-Benzoylphosphoadenosine (mitochondria)
3.6.1.20
ATP
3.6.1.17
3.6.1.61
ApppppA
2.7.7.53
3.6.1.41
Adenylylsulfate
2.7.7.4
Sulfate
2.7.1.25
3'-Phosphoadenylate
3.6.2.2
3'-Phosphoadenylyl-sulfate
ApppA
3.6.1.29
ADP
PDE
3.6.1.8
Inosine 5'-tetraphosphate
3.6.1.14
ITP
IDP
3.6.1.5
3.6.1.73
3.6.1.6
6.3.4.4
Adenylo-succinate
4.3.2.2
3.1.4.17
3.1.4.53
Arginine biosynthesis
2',3'-Cyclic AMP
3.1.4.16
3'-AMP
NH3
3.6.1.9
Adenosine 5'-tetraphosphate
pppGpp
3.6.1.11
3.6.1.40
ppGpp
3.1.7.2
2.7.6.5
3',5'-Cyclic GMP
4.6.1.1
4.6.1.2
RNA
Riboflavin metabolism
GTP
GDP
2.7.1.40
2.7.4.8
GMP
6.3.5.2
XMP
1.1.1.205
IMP
3.5.4.6
AMP
2.7.4.3
2.7.4.10
3'-5'-Cyclic AMP
1.7.1.7
5'-Butyrylphosphoinosine
3.6.1.15
3.6.1.17
GppppG
Folate biosynthesis
1.17.4.2
1.1.98.6
1.17.4.1
DNA
dGTP
dGDP
3.1.5.1
Deoxyguanosine
3.1.3.5
2.7.1.113
dGMP
2.7.1.73
XppppX
XTP
Guanosine
Xanthosine
3.1.3.99
2.4.2.8
3.2.2.12
2.7.1.20
2.7.1.74
Inosine
3.5.4.4
Adenosine
3.2.2.4
dADP
dATP
2.7.4.11
dAMP
Deoxyadenosine
2.7.1.76
2.4.2.1
2.4.2.15
2.4.2.22
3.2.2.1
3.2.2.8
3.2.2.2
3.2.2.7
2.4.2.-
Guanine
3.5.4.3
Xanthine
1.17.3.2
1.17.1.4
Hypoxanthine
3.5.4.2
Adenine
Deoxyinosine
3.1.3.6
3'-GMP
2',3'-Cyclic GMP
Urate-3-ribonucleoside
2.4.2.16
Urate
1.14.13.113
1.7.3.3
3.5.2.-
5-Amino-4-imidazole carboxylate
5-Ureido-4-imidazole carboxylate
3.5.3.-
4.1.1.-
Aminoimidazole
3.5.4.8
Imidazolone
Formimino-glycine
2.1.2.4
Glycine
Glycine, serine and threonine metabolism
2.6.1.112
5-Hydroxyisourate
3.5.2.17
5-Hydroxy-2-oxo-4-ureido-2,5-dihydro-1H-imidazole-5-carboxylate
4.1.1.97
(S)-Allantoin
3.5.2.5
Allantoate
3.5.3.9
Ureidoglycine
3.5.3.26
Ureido-glycolate
1.1.1.154
1.1.1.350
Oxalureate
2.1.3.5
Carbamoyl phosphate
2.7.2.2
Hydrogen carbonate
spontaneous
5.1.99.3
(R)-Allantoin
3.5.3.4
4.3.2.3
Urea
3.5.1.116
3.5.1.5
Glyoxylate
Glyoxylate metabolism
NH3
CO2
Oxamate
3.5.1.126
Oxalate
00230 7/7/20
(c) Kanehisa Laboratories

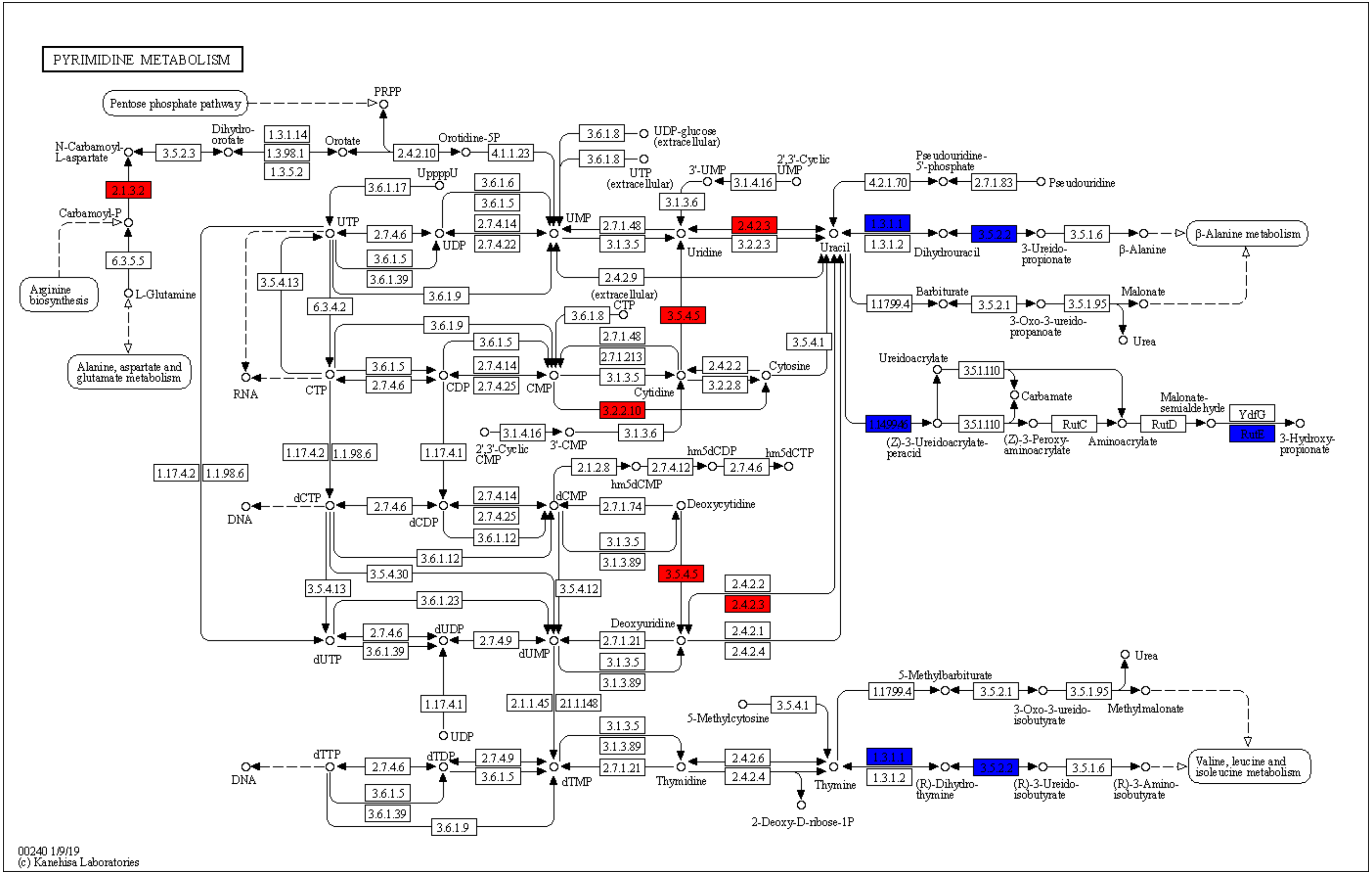
PYRIMIDINE METABOLISM
Pentose phosphate pathway
PRPP
N-Carbamoyl-L-aspartate
Dihydro-orotate
Orotate
Orotidine-5P
UDP-glucose (extracellular)
UTP (extracellular)
3'-UMP
2',3'-Cyclic UMP
Pseudouridine-5'-phosphate
Pseudouridine
Carbamoyl-P
L-Glutamine
Arginine biosynthesis
Alanine, aspartate and glutamate metabolism
UpppU
UTP
UDP
UMP
Uridine
Uracil
Dihydrouracil
3-Ureido-propionate
β-Alanine
β-Alanine metabolism
Barbiturate
3-Oxo-3-ureido-propanoate
Malonate
Urea
CTP (extracellular)
Cytosine
Cytidine
RNA
CTP
CDP
CMP
2',3'-Cyclic CMP
3'-CMP
Ureidoacrylate
Carbamate
(Z)-3-Ureidoacrylate-peracid
(Z)-3-Peroxy-aminoacrylate
Aminoacrylate
Malonate-semialdehyde
3-Hydroxy-propionate
hm5dCDP
hm5dCTP
hm5dCMP
DNA
dCTP
dCDP
dCMP
Deoxycytidine
Deoxyuridine
dUTP
dUDP
dUMP
UDP
5-Methylbarbiturate
3-Oxo-3-ureido-isobutyrate
Methylmalonate
5-Methylcytosine
dTTP
dTDP
dTMP
Thymidine
Thymine
(R)-Dihydro-thymine
(R)-3-Ureido-isobutyrate
(R)-3-Amino-isobutyrate
Valine, leucine and isoleucine metabolism
2-Deoxy-D-ribose-1P
2.1.3.2
3.5.2.3
1.3.1.14
1.3.98.1
1.3.5.2
2.4.2.10
4.1.1.23
6.3.5.5
3.6.1.17
3.6.1.6
3.6.1.5
2.7.4.14
2.7.4.22
2.7.4.6
3.6.1.39
3.6.1.9
3.5.4.13
6.3.4.2
3.6.1.8
2.7.1.48
3.1.3.5
3.1.3.6
3.1.4.16
2.4.2.3
3.2.2.3
2.4.2.9
3.5.4.5
4.2.1.70
2.7.1.83
1.3.1.1
1.3.1.2
3.5.2.2
3.5.1.6
1.17.99.4
3.5.2.1
3.5.1.95
2.7.1.213
2.4.2.2
3.2.2.8
3.5.4.1
3.2.2.10
2.7.4.25
3.5.1.110
1.14.99.46
RutC
RutD
YdfG
RutE
1.17.4.2
1.1.98.6
1.17.4.1
2.1.2.8
2.7.4.12
2.7.1.74
3.6.1.12
3.1.3.89
3.5.4.30
3.5.4.12
3.6.1.23
2.4.2.1
2.4.2.4
2.7.4.9
2.7.1.21
2.1.1.45
2.1.1.148
2.4.2.6
00240 1/9/19
(c) Kanehisa Laboratories

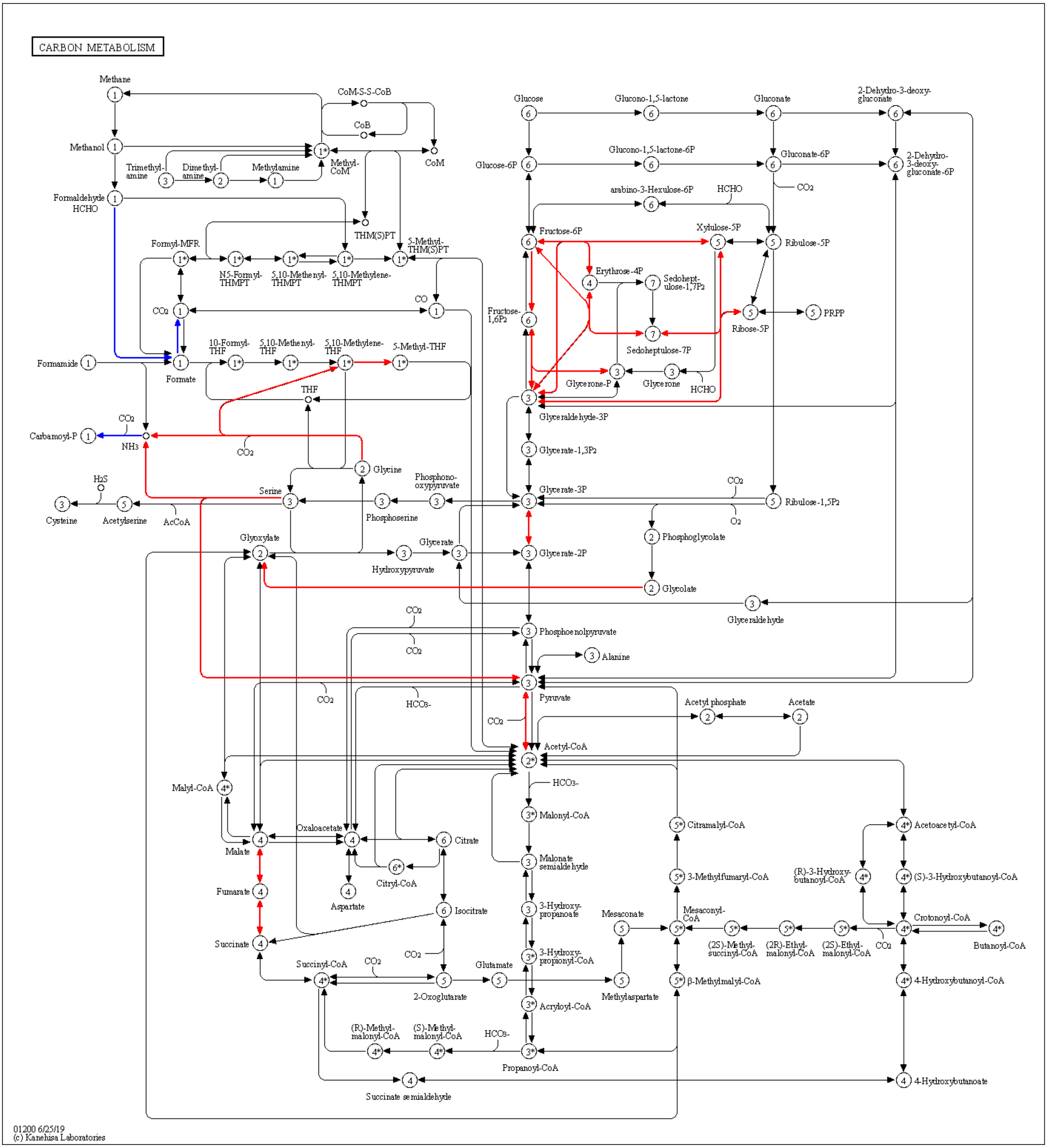
CARBON METABOLISM
Methane
CoM-S-S-CoB
CoB
CoM
Methanol
Trimethyl-amine
Dimethyl-amine
Methylamine
Methyl-CoM
Formaldehyde HCHO
THM(S)PT
5-Methyl-THM(S)PT
Formyl-MFR
N5-Formyl-THMPT
5,10-Methenyl-THMPT
5,10-Methylene-THMPT
CO2
CO
Formamide
Formate
10-Formyl-THF
5,10-Methenyl-THF
5,10-Methylene-THF
5-Methyl-THF
THF
CO2
Carbamoyl-P
NH3
CO2
Glycine
H2S
Cysteine
Acetylserine
AcCoA
Serine
Phosphoserine
Phosphono-oxypyruvate
Glyoxylate
Hydroxypyruvate
Glycerate
Glucose
Glucono-1,5-lactone
Gluconate
2-Dehydro-3-deoxy-gluconate
Glucose-6P
Glucono-1,5-lactone-6P
Gluconate-6P
2-Dehydro-3-deoxy-gluconate-6P
CO2
arabino-3-Hexulose-6P
HCHO
Fructose-6P
Xylulose-5P
Ribulose-5P
Erythrose-4P
Sedohept-ulose-1,7P2
Fructose-1,6P2
Ribose-5P
PRPP
Sedoheptulose-7P
Glycerone-P
Glycerone
HCHO
Glyceraldehyde-3P
Glycerate-1,3P2
Glycerate-3P
CO2
Ribulose-1,5P2
O2
Phosphoglycolate
Glycerate-2P
Glycolate
Glyceraldehyde
CO2
Phosphoenolpyruvate
CO2
Alanine
Pyruvate
CO2
HCO3-
CO2
Acetyl phosphate
Acetate
Acetyl-CoA
HCO3-
Malyl-CoA
Malonyl-CoA
Citramalyl-CoA
Acetoacetyl-CoA
Oxaloacetate
Malate
Citrate
Citryl-CoA
Malonate semialdehyde
3-Methylfumaryl-CoA
(R)-3-Hydroxy-butanoyl-CoA
(S)-3-Hydroxybutanoyl-CoA
Fumarate
Aspartate
Isocitrate
3-Hydroxy-propanoate
Mesaconate
Mesaconyl-CoA
Crotonoyl-CoA
Succinate
3-Hydroxy-propionyl-CoA
(2S)-Methyl-succinyl-CoA
(2R)-Ethyl-malonyl-CoA
(2S)-Ethyl-malonyl-CoA
CO2
Butanoyl-CoA
Succinyl-CoA
CO2
CO2
Glutamate
2-Oxoglutarate
Methylaspartate
β-Methylmalyl-CoA
4-Hydroxybutanoyl-CoA
Acryloyl-CoA
(R)-Methyl-malonyl-CoA
(S)-Methyl-malonyl-CoA
HCO3-
Propanoyl-CoA
Succinate semialdehyde
4-Hydroxybutanoate
01200 6/25/19
(c) Kanehisa Laboratories

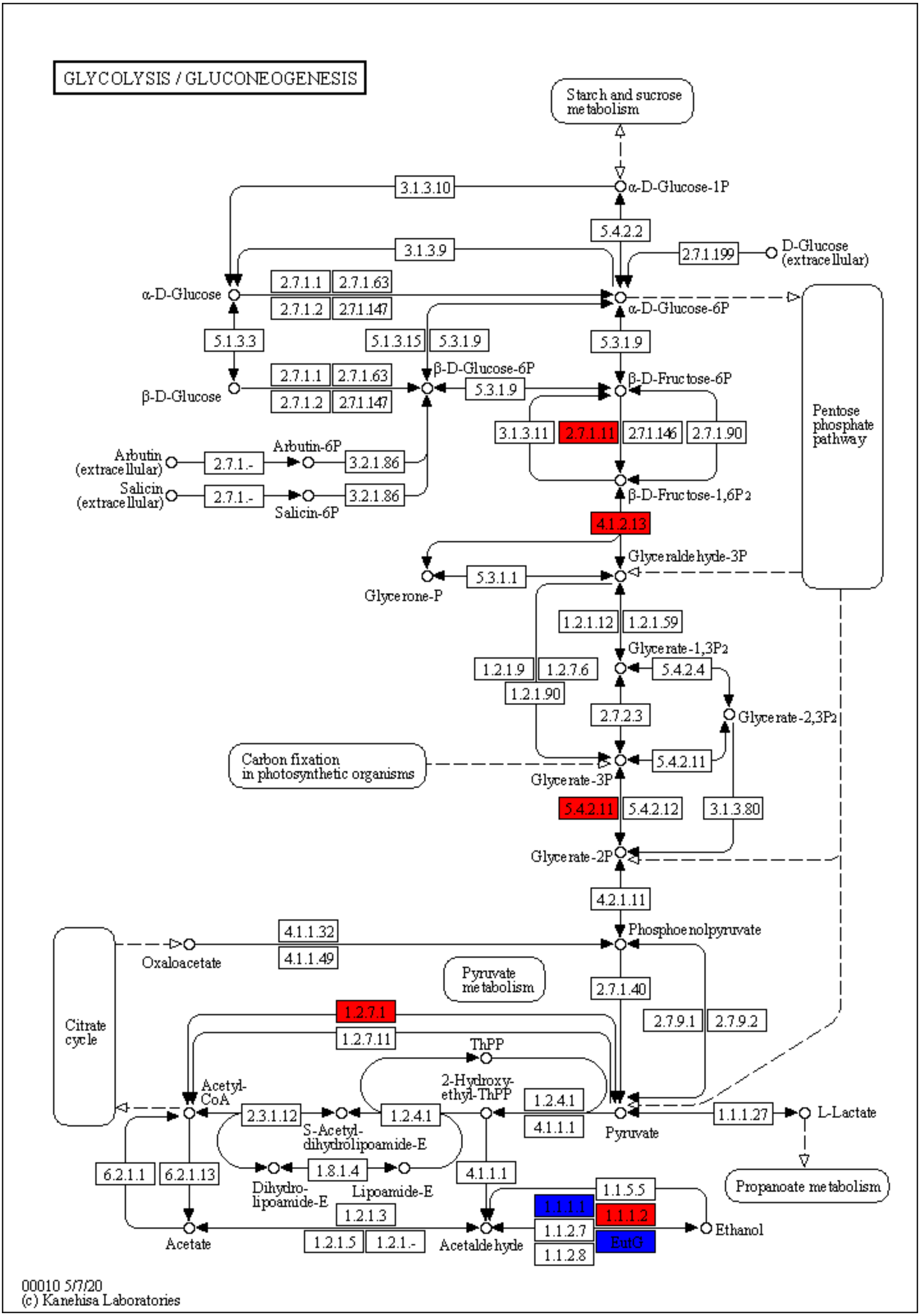
GLYCOLYSIS / GLUCONEOGENESIS
Starch and sucrose metabolism
α-D-Glucose-1P
3.1.3.10
5.4.2.2
3.1.3.9
2.7.1.199
D-Glucose (extracellular)
α-D-Glucose
2.7.1.1
2.7.1.63
2.7.1.2
2.7.1.147
α-D-Glucose-6P
5.1.3.3
5.1.3.15
5.3.1.9
β-D-Glucose-6P
β-D-Glucose
β-D-Fructose-6P
Pentose phosphate pathway
3.1.3.11
2.7.1.11
2.7.1.146
2.7.1.90
Arbutin (extracellular)
2.7.1.-
Arbutin-6P
3.2.1.86
Salicin (extracellular)
Salicin-6P
β-D-Fructose-1,6P2
4.1.2.13
Glyceraldehyde-3P
5.3.1.1
Glycerone-P
1.2.1.12
1.2.1.59
Glycerate-1,3P2
1.2.1.9
1.2.7.6
1.2.1.90
5.4.2.4
2.7.2.3
Glycerate-2,3P2
Carbon fixation in photosynthetic organisms
5.4.2.11
Glycerate-3P
5.4.2.12
3.1.3.80
Glycerate-2P
4.2.1.11
Phosphoenolpyruvate
4.1.1.32
4.1.1.49
Oxaloacetate
Pyruvate metabolism
2.7.1.40
Citrate cycle
1.2.7.1
1.2.7.11
2.7.9.1
2.7.9.2
ThPP
2-Hydroxy-ethyl-ThPP
Acetyl-CoA
2.3.1.12
1.2.4.1
4.1.1.1
Pyruvate
1.1.1.27
L-Lactate
S-Acetyl-dihydrolipoamide-E
6.2.1.1
6.2.1.13
1.8.1.4
Dihydro-lipoamide-E
Lipoamide-E
Propanoate metabolism
1.1.5.5
1.1.1.1
1.1.1.2
1.2.1.3
1.1.2.7
ExaG
1.1.2.8
Ethanol
Acetate
1.2.1.5
1.2.1.-
Acetaldehyde
00010 5/7/20
(c) Kanehisa Laboratories

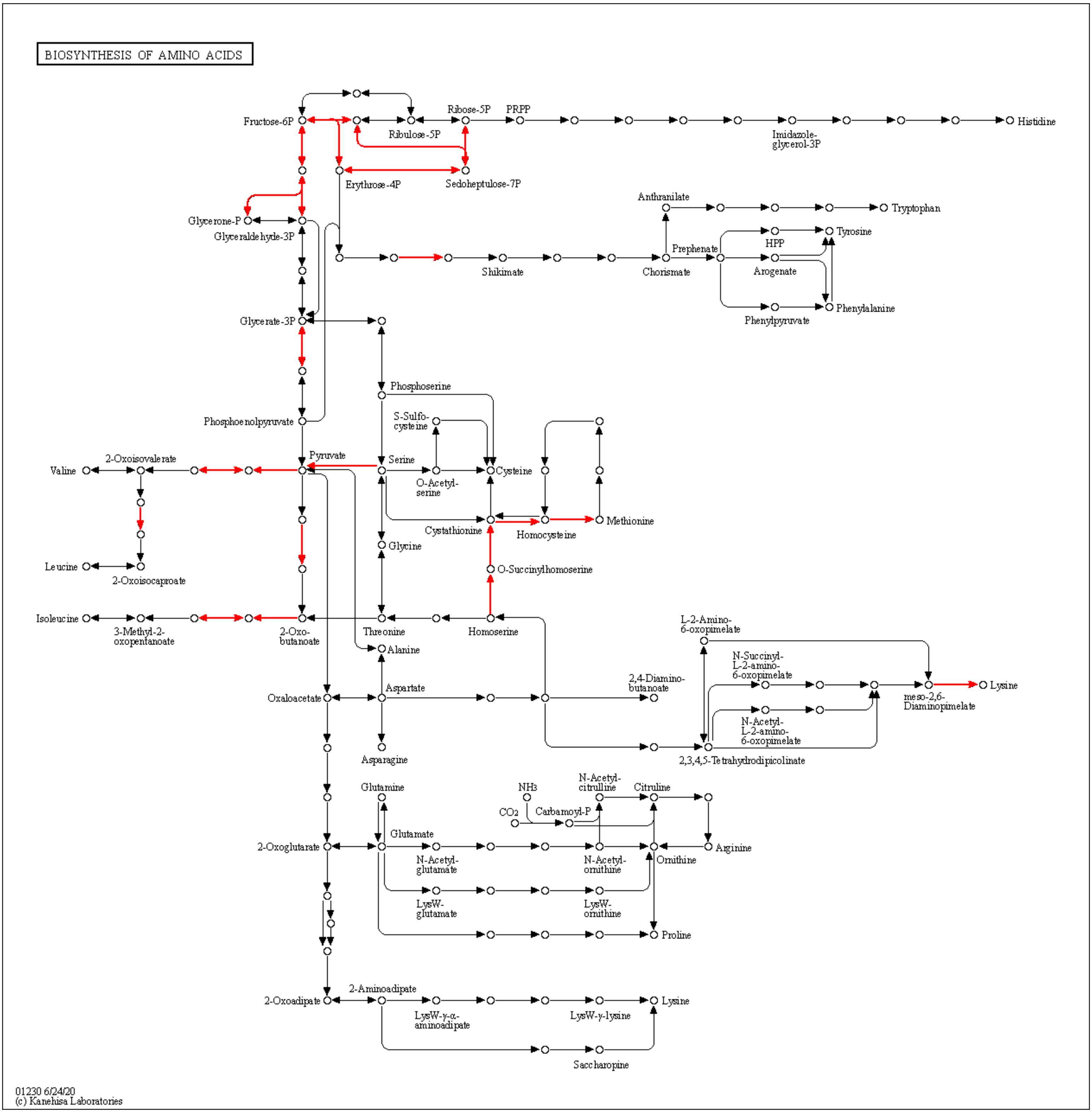
BIOSYNTHESIS OF AMINO ACIDS
Fructose-6P
Ribose-5P
PRPP
Ribulose-5P
Histidine
Imidazole-glycerol-3P
Erythrose-4P
Sedoheptulose-7P
Glycerone-P
Glyceraldehyde-3P
Anthranilate
Tryptophan
Tyrosine
HPP
Prephenate
Shikimate
Chorismate
Arogenate
Phenylalanine
Phenylpyruvate
Glycerate-3P
Phosphoserine
S-Sulfo-cysteine
Phosphoenolpyruvate
2-Oxoisovalerate
Valine
Pyruvate
Serine
Cysteine
O-Acetyl-serine
Cystathionine
Homocysteine
Methionine
Glycine
Leucine
2-Oxoisocaproate
O-Succinylhomoserine
Isoleucine
3-Methyl-2-oxopentanoate
2-Oxo-butanoate
Threonine
Homoserine
Alanine
L-2-Amino-6-oxopimelate
N-Succinyl-L-2-amino-6-oxopimelate
Lysine
meso-2,6-Diaminopimelate
2,4-Diamino-butanoate
Oxaloacetate
Aspartate
N-Acetyl-L-2-amino-6-oxopimelate
Asparagine
2,3,4,5-Tetrahydrodipicolinate
Glutamine
$NH_3$
N-Acetyl-citrulline
Citrulline
$CO_2$
Carbamoyl-P
Glutamate
2-Oxoglutarate
N-Acetyl-glutamate
N-Acetyl-ornithine
Ornithine
Arginine
LysW-glutamate
LysW-ornithine
Proline
2-Aminoadipate
2-Oxoadipate
LysW-γ-α-aminoadipate
LysW-γ-lysine
Lysine
Saccharopine
01230 6/24/20
(c) Kanehisa Laboratories

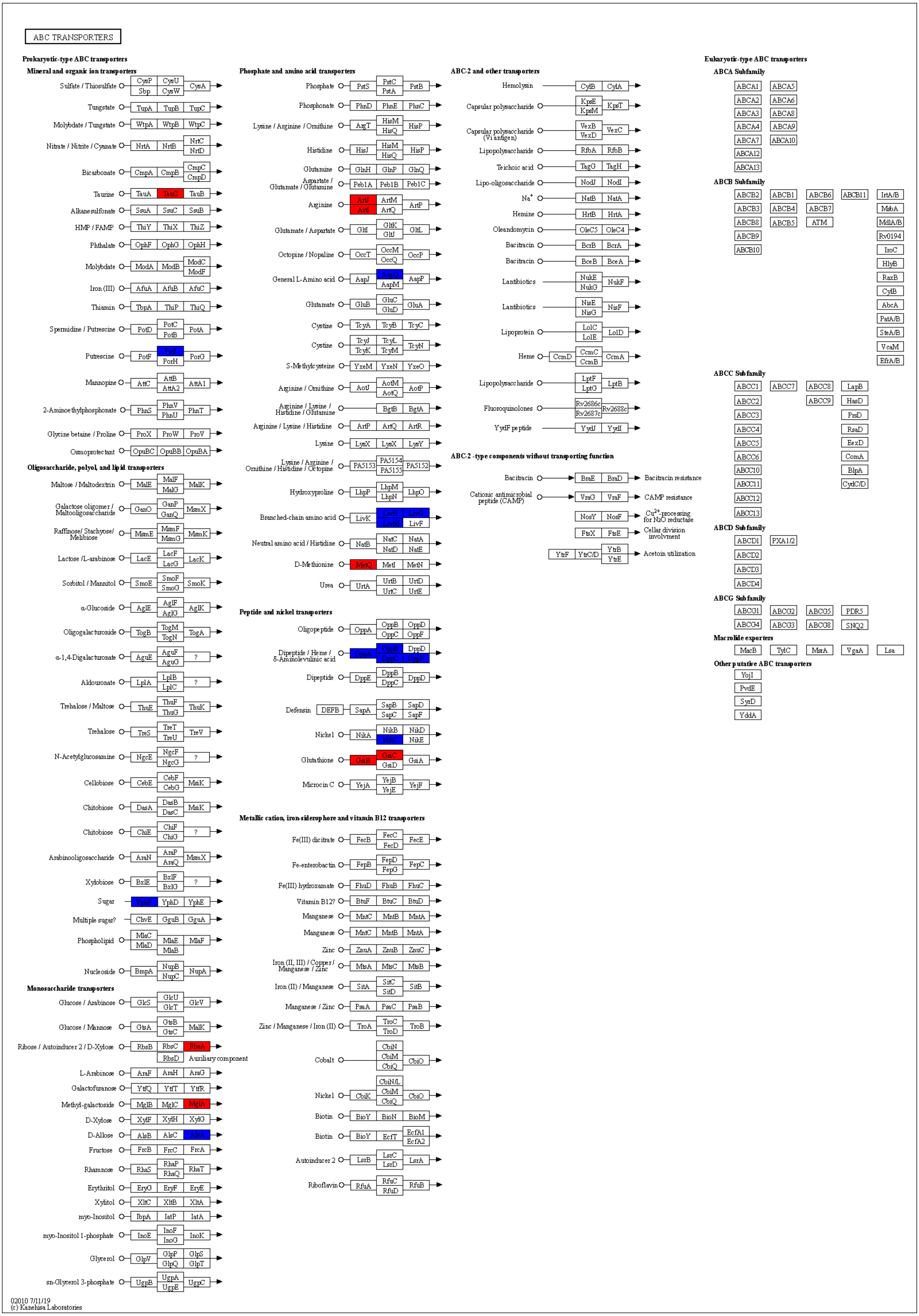
ABC TRANSPORTERS
Prokaryotic-type ABC transporters
Mineral and organic ion transporters
Sulfate / Thiosulfate
CysP
Sbp
CysU
CysW
CysA
Tungstate
TupA
TupB
TupC
Molybdate / Tungstate
WtpA
WtpB
WtpC
Nitrate / Nitrite / Cyanate
NrtA
NrtB
NrtC
NrtD
Bicarbonate
CmpA
CmpB
CmpC
CmpD
Taurine
TauA
TauC
TauB
Alkanesulfonate
SsuA
SsuC
SsuB
HMP / FAMP
ThiY
ThiX
ThiZ
Phthalate
OphF
OphG
OphH
Molybdate
ModA
ModB
ModC
ModF
Iron (III)
AfuA
AfuB
AfuC
Thiamin
TbpA
ThiP
ThiQ
Spermidine / Putrescine
PotD
PotC
PotB
PotA
Putrescine
PotF
PotH
PotG
Mannopine
AttC
AttB
AttA2
AttA1
2-Aminoethylphosphonate
PhnS
PhnV
PhnU
PhnT
Glycine betaine / Proline
ProX
ProW
ProV
Osmoprotectant
OpuBC
OpuBB
OpuBA
Oligosaccharide, polyol, and lipid transporters
Maltose / Maltodextrin
MalE
MalF
MalG
MalK
Galactose oligomer / Maltooligosaccharide
GanO
GanP
GanQ
MsmX
Raffinose/ Stachyose/ Melibiose
MsmE
MsmF
MsmG
MsmK
Lactose /L-arabinose
LacE
LacF
LacG
LacK
Sorbitol / Mannitol
SmoE
SmoF
SmoG
SmoK
α-Glucoside
AglE
AglF
AglG
AglK
Oligogalacturonide
TogB
TogM
TogN
TogA
α-1,4-Digalacturonate
AguE
AguF
AguG
?
Aldouronate
LplA
LplB
LplC
?
Trehalose / Maltose
ThuE
ThuF
ThuG
ThuK
Trehalose
TreS
TreT
TreU
TreV
N-Acetylglucosamine
NgcE
NgcF
NgcG
?
Cellobiose
CebE
CebF
CebG
MsiK
Chitobiose
DasA
DasB
DasC
MsiK
Chitobiose
ChiE
ChiF
ChiG
?
Arabinooligosaccharide
AraN
AraP
AraQ
MsmX
Xylobiose
BxlE
BxlF
BxlG
?
Sugar
YphD
YphE
Multiple sugar?
ChvE
GguB
GguA
Phospholipid
MlaC
MlaD
MlaE
MlaB
MlaF
Nucleoside
BmpA
NupB
NupC
NupA
Monosaccharide transporters
Glucose / Arabinose
GlcS
GlcU
GlcT
GlcV
Glucose / Mannose
GtsA
GtsB
GtsC
MalK
Ribose / Autoinducer 2 / D-Xylose
RbsB
RbsC
RbsA
RbsD
Auxiliary component
L-Arabinose
AraF
AraH
AraG
Galactofuranose
YtfQ
YtfT
YtfR
Methyl-galactoside
MglB
MglC
MglA
D-Xylose
XylF
XylH
XylG
D-Allose
AlsB
AlsC
AlsA
Fructose
FrcB
FrcC
FrcA
Rhamnose
RhaS
RhaP
RhaQ
RhaT
Erythritol
EryG
EryF
EryE
Xylitol
XltC
XltB
XltA
myo-Inositol
IbpA
IatP
IatA
myo-Inositol 1-phosphate
InoE
InoF
InoG
InoK
Glycerol
GlpV
GlpP
GlpQ
GlpS
GlpT
sn-Glycerol 3-phosphate
UgpB
UgpA
UgpE
UgpC
Phosphate and amino acid transporters
Phosphate
PstS
PstC
PstA
PstB
Phosphonate
PhnD
PhnE
PhnC
Lysine / Arginine / Ornithine
ArgT
HisM
HisQ
HisP
Histidine
HisJ
HisM
HisQ
HisP
Glutamine
GlnH
GlnP
GlnQ
Aspartate / Glutamate / Glutamine
Peb1A
Peb1B
Peb1C
Arginine
ArtJ
ArtI
ArtM
ArtQ
ArtP
Glutamate / Aspartate
GltI
GltK
GltJ
GltL
Octopine / Nopaline
OccT
OccM
OccQ
OccP
General L-Amino acid
AapJ
AapM
AapP
Glutamate
GluB
GluC
GluD
GluA
Cystine
TcyA
TcyB
TcyC
Cystine
TcyJ
TcyK
TcyL
TcyM
TcyN
S-Methylcysteine
YxeM
YxeN
YxeO
Arginine / Ornithine
AotJ
AotM
AotQ
AotP
Arginine / Lysine / Histidine / Glutamine
BgtB
BgtA
Arginine / Lysine / Histidine
ArtP
ArtQ
ArtR
Lysine
LysX
LysX
LysY
Lysine / Arginine / Ornithine / Histidine / Octopine
PA5153
PA5154
PA5155
PA5152
Hydroxyproline
LhpP
LhpM
LhpN
LhpO
Branched-chain amino acid
LivK
LivH
LivM
LivG
LivF
Neutral amino acid / Histidine
NatB
NatC
NatD
NatA
NatE
D-Methionine
MetQ
MetI
MetN
Urea
UrtA
UrtB
UrtC
UrtD
UrtE
Peptide and nickel transporters
Oligopeptide
OppA
OppB
OppC
OppD
OppF
Dipeptide / Heme / δ-Aminolevulinic acid
DppA
DppB
DppC
DppD
DppF
Dipeptide
DppE
DppB
DppC
DppD
Defensin
DEFB
SapA
SapB
SapC
SapD
SapF
Nickel
NikA
NikB
NikC
NikD
NikE
Glutathione
GsiB
GsiC
GsiD
GsiA
Microcin C
YejA
YejB
YejE
YejF
Metallic cation, iron-siderophore and vitamin B12 transporters
Fe(III) dicitrate
FecB
FecC
FecD
FecE
Fe-enterobactin
FepB
FepD
FepG
FepC
Fe(III) hydroxamate
FhuD
FhuB
FhuC
Vitamin B12?
BtuF
BtuC
BtuD
Manganese
MntC
MntB
MntA
Manganese
MntC
MntB
MntA
Zinc
ZnuA
ZnuB
ZnuC
Iron (II, III) / Copper / Manganese / Zinc
MtsA
MtsC
MtsB
Iron (II) / Manganese
SitA
SitC
SitD
SitB
Manganese / Zinc
PsaA
PsaC
PsaB
Zinc / Manganese / Iron (II)
TroA
TroC
TroD
TroB
Cobalt
CbiN
CbiM
CbiQ
CbiO
Nickel
CbiK
CbiN/L
CbiM
CbiQ
CbiO
Biotin
BioY
BioN
BioM
Biotin
BioY
EcfT
EcfA1
EcfA2
Autoinducer 2
LsrB
LsrC
LsrD
LsrA
Riboflavin
RfuA
RfuC
RfuD
RfuB
ABC-2 and other transporters
Hemolysin
CylB
CylA
Capsular polysaccharide
KpsE
KpsM
KpsT
Capsular polysaccharide (Vi antigen)
VexB
VexD
VexC
Lipopolysaccharide
RfbA
RfbB
Teichoic acid
TagG
TagH
Lipo-oligosaccharide
NodJ
NodI
Na+
NatB
NatA
Hemine
HrtB
HrtA
Oleandomycin
OleC5
OleC4
Bacitracin
BcrB
BcrA
Bacitracin
BceB
BceA
Lantibiotics
NukE
NukG
NukF
Lantibiotics
NisE
NisG
NisF
Lipoprotein
LolC
LolE
LolD
Heme
CcmD
CcmC
CcmB
CcmA
Lipopolysaccharide
LptF
LptG
LptB
Fluoroquinolones
Rv2686c
Rv2687c
Rv2688c
YydF peptide
YydJ
YydI
ABC-2 -type components without transporting function
Bacitracin
BraE
BraD
Bacitracin resistance
Cationic antimicrobial peptide (CAMP)
VraG
VraF
CAMP resistance
NosY
NosF
Cu2+-processing for N2O reductase
FtsX
FtsE
Cellar division involvment
YbtF
YtrC/D
YtrB
YtrE
Acetoin utilization
Eukaryotic-type ABC transporters
ABCA Subfamily
ABCA1
ABCA5
ABCA2
ABCA6
ABCA3
ABCA8
ABCA4
ABCA9
ABCA7
ABCA10
ABCA12
ABCA13
ABCB Subfamily
ABCB2
ABCB1
ABCB6
ABCB11
IrtA/B
ABCB3
ABCB4
ABCB7
MsbA
ABCB8
ABCB5
ATM
MdlA/B
ABCB9
Rv0194
ABCB10
IrtC
HlyB
RaxB
CylB
AbcA
PatA/B
SteA/B
VcaM
EfrA/B
ABCC Subfamily
ABCC1
ABCC7
ABCC8
LapB
ABCC2
ABCC9
HasD
ABCC3
PrsD
ABCC4
RsaD
ABCC5
EexD
ABCC6
ComA
ABCC10
BlpA
ABCC11
CydC/D
ABCC12
ABCC13
ABCD Subfamily
ABCD1
PXA1/2
ABCD2
ABCD3
ABCD4
ABCG Subfamily
ABCG1
ABCG2
ABCG5
PDR5
ABCG4
ABCG3
ABCG8
SNQ2
Macrolide exporters
MacB
TylC
MsrA
VgaA
Lsa
Other putative ABC transporters
YojI
PvdE
SyrD
YddA
02010 7/11/19
(c) Kanehisa Laboratories

**Supplementary Fig. 9 Representative metabolic pathway maps at 12 h from KEGG mapper.** Red indicates the upregulation and blue indicates the downregulation of each gene.

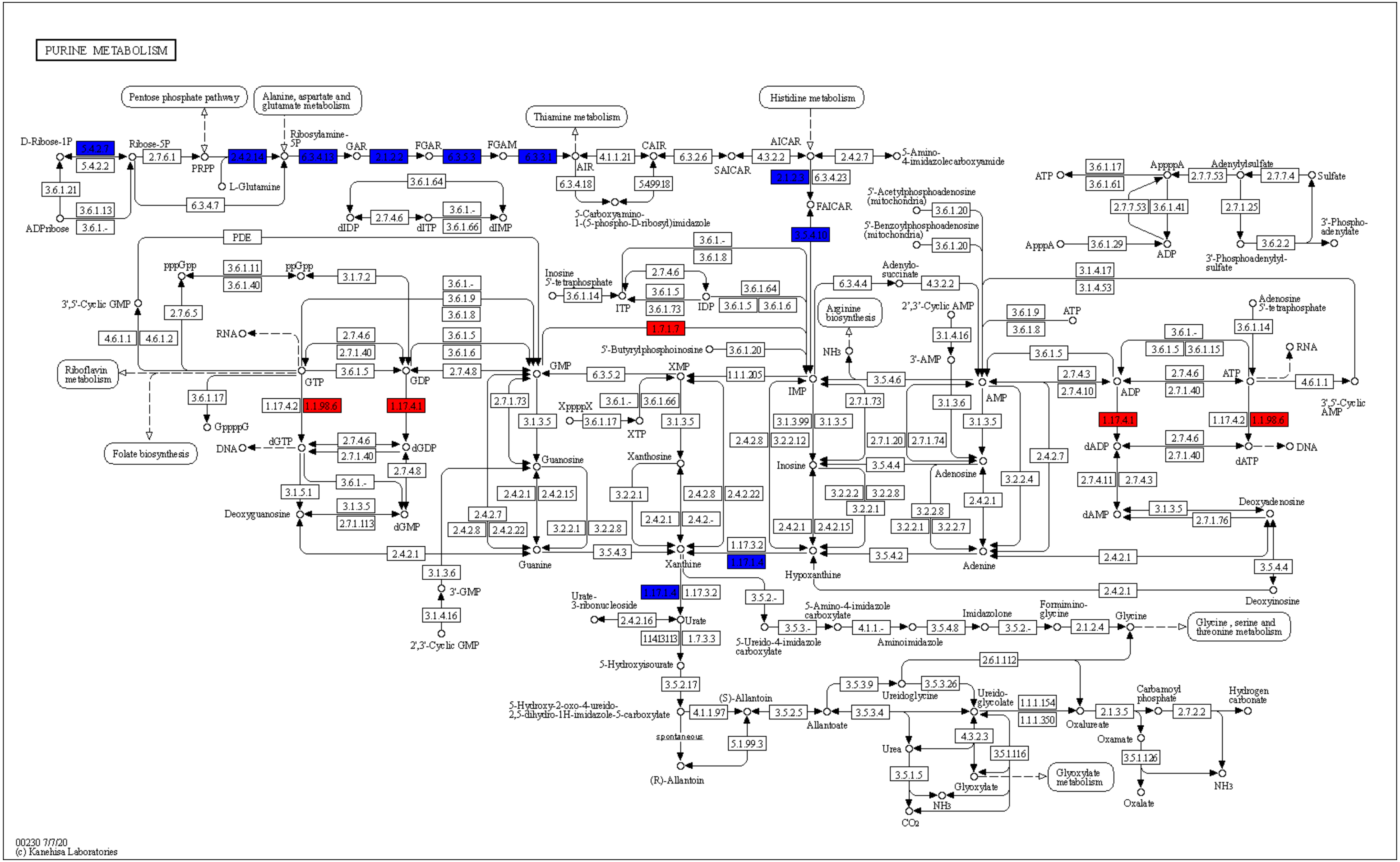
PURINE METABOLISM
Pentose phosphate pathway
Alanine, aspartate and glutamate metabolism
Thiamine metabolism
Histidine metabolism
D-Ribose-1P
Ribose-5P
PRPP
L-Glutamine
Ribosylamine-5P
GAR
FGAR
FGAM
AIR
CAIR
SAICAR
AICAR
5-Amino-4-imidazolecarboxyamide
FAICAR
5-Carboxyamino-1-(5-phospho-D-ribosyl)imidazole
ADPribose
dIDP
dITP
dIMP
PDE
pppGpp
ppGpp
3',5'-Cyclic GMP
RNA
Riboflavin metabolism
Folate biosynthesis
GTP
GDP
GMP
GppppG
DNA
dGTP
dGDP
dGMP
Deoxyguanosine
Guanosine
Guanine
3'-GMP
2',3'-Cyclic GMP
Inosine 5'-tetraphosphate
ITP
IDP
IMP
XppppX
XTP
XMP
Xanthosine
Xanthine
5'-Butyrylphosphoinosine
Inosine
Hypoxanthine
Urate-3-ribonucleoside
Urate
5-Hydroxyisourate
5-Hydroxy-2-oxo-4-ureido-2,5-dihydro-1H-imidazole-5-carboxylate
(S)-Allantoin
spontaneous
(R)-Allantoin
5-Ureido-4-imidazole carboxylate
5-Amino-4-imidazole carboxylate
Aminoimidazole
Imidazolone
Formimino-glycine
Glycine
Glycine, serine and threonine metabolism
Allantoate
Ureidoglycine
Ureido-glycolate
Urea
Glyoxylate
Glyoxylate metabolism
NH3
CO2
Oxalureate
Carbamoyl phosphate
Oxamate
Oxalate
Hydrogen carbonate
Arginine biosynthesis
Adenylo-succinate
5'-Acetylphosphoadenosine (mitochondria)
5'-Benzoylphosphoadenosine (mitochondria)
2',3'-Cyclic AMP
3'-AMP
AMP
Adenosine
Adenine
ATP
ApppA
ApppA
Adenylylsulfate
Sulfate
3'-Phosphoadenylyl-sulfate
3'-Phospho-adenylate
Adenosine 5'-tetraphosphate
ADP
dADP
dATP
dAMP
Deoxyadenosine
Deoxyinosine
3',5'-Cyclic AMP
00230 7/7/20
(c) Kanehisa Laboratories

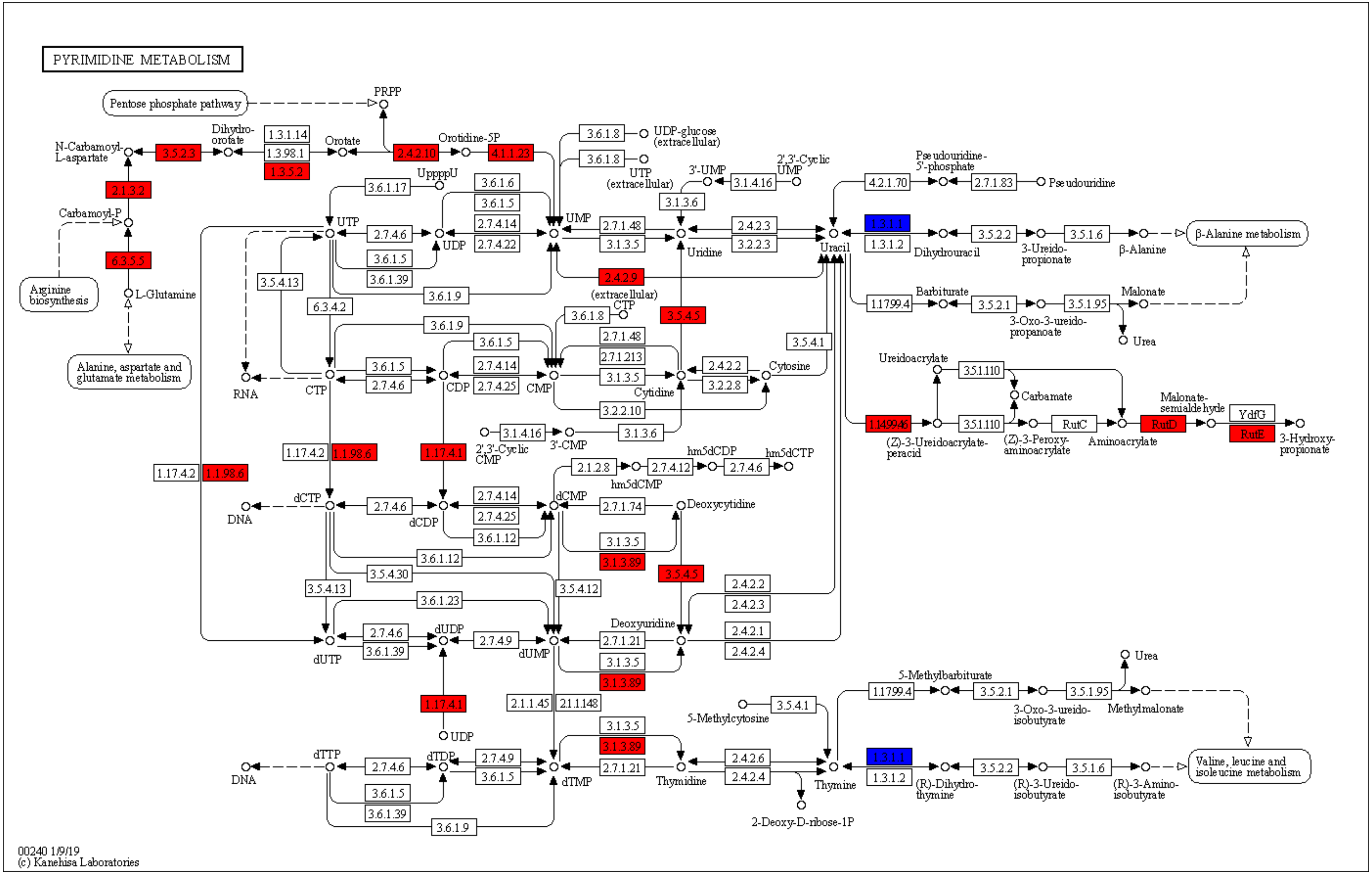
PYRIMIDINE METABOLISM
Pentose phosphate pathway
PRPP
N-Carbamoyl-L-aspartate
3.5.2.3
Dihydro-orotate
1.3.1.14
1.3.98.1
1.3.5.2
Orotate
2.4.2.10
Orotidine-5P
4.1.1.23
2.1.3.2
Carbamoyl-P
6.3.5.5
L-Glutamine
Arginine biosynthesis
Alanine, aspartate and glutamate metabolism
3.6.1.8
UDP-glucose (extracellular)
3.6.1.8
UTP (extracellular)
3'-UMP
3.1.4.16
2',3'-Cyclic UMP
3.1.3.6
UppppU
3.6.1.17
3.6.1.6
3.6.1.5
2.7.4.14
2.7.4.22
UTP
2.7.4.6
UDP
UMP
2.7.1.48
3.1.3.5
2.4.2.3
3.2.2.3
Uridine
Uracil
3.6.1.5
3.6.1.39
3.5.4.13
3.6.1.9
6.3.4.2
2.4.2.9
(extracellular) CTP
3.6.1.8
3.5.4.5
3.6.1.9
3.6.1.5
2.7.1.48
2.7.1.213
2.7.4.14
2.7.4.25
3.6.1.5
2.7.4.6
RNA
CTP
CDP
CMP
3.1.3.5
Cytidine
2.4.2.2
3.2.2.8
Cytosine
3.5.4.1
3.2.2.10
3.1.4.16
2',3'-Cyclic CMP
3'-CMP
3.1.3.6
1.17.4.2
1.1.98.6
1.17.4.1
1.17.4.2
1.1.98.6
2.1.2.8
hm5dCMP
2.7.4.12
hm5dCDP
2.7.4.6
hm5dCTP
dCTP
DNA
2.7.4.6
dCDP
2.7.4.14
2.7.4.25
3.6.1.12
dCMP
2.7.1.74
Deoxycytidine
3.1.3.5
3.1.3.89
3.5.4.5
3.6.1.12
3.5.4.30
3.5.4.13
3.5.4.12
3.6.1.23
2.4.2.2
2.4.2.3
Deoxyuridine
2.4.2.1
2.4.2.4
2.7.4.6
3.6.1.39
dUTP
dUDP
2.7.4.9
dUMP
2.7.1.21
3.1.3.5
3.1.3.89
1.17.4.1
UDP
2.1.1.45
2.1.1.148
3.1.3.5
3.1.3.89
5-Methylcytosine
3.5.4.1
dTTP
DNA
2.7.4.6
dTDP
2.7.4.9
3.6.1.5
dTMP
2.7.1.21
Thymidine
2.4.2.6
2.4.2.4
Thymine
2-Deoxy-D-ribose-1P
3.6.1.5
3.6.1.39
3.6.1.9
Pseudouridine-5'-phosphate
4.2.1.70
2.7.1.83
Pseudouridine
1.3.1.1
1.3.1.2
Dihydrouracil
3.5.2.2
3-Ureido-propionate
3.5.1.6
β-Alanine
β-Alanine metabolism
1.17.99.4
Barbiturate
3.5.2.1
3-Oxo-3-ureido-propanoate
3.5.1.95
Malonate
Urea
Ureidoacrylate
3.5.1.110
Carbamate
1.14.99.46
3.5.1.110
(Z)-3-Ureidoacrylate-peracid
(Z)-3-Peroxy-aminoacrylate
RutC
Aminoacrylate
Malonate-semialdehyde
RutD
YdfG
RutE
3-Hydroxy-propionate
Urea
5-Methylbarbiturate
1.17.99.4
3.5.2.1
3-Oxo-3-ureido-isobutyrate
3.5.1.95
Methylmalonate
Valine, leucine and isoleucine metabolism
1.3.1.1
1.3.1.2
(R)-Dihydro-thymine
3.5.2.2
(R)-3-Ureido-isobutyrate
3.5.1.6
(R)-3-Amino-isobutyrate
00240 1/9/19
(c) Kanehisa Laboratories

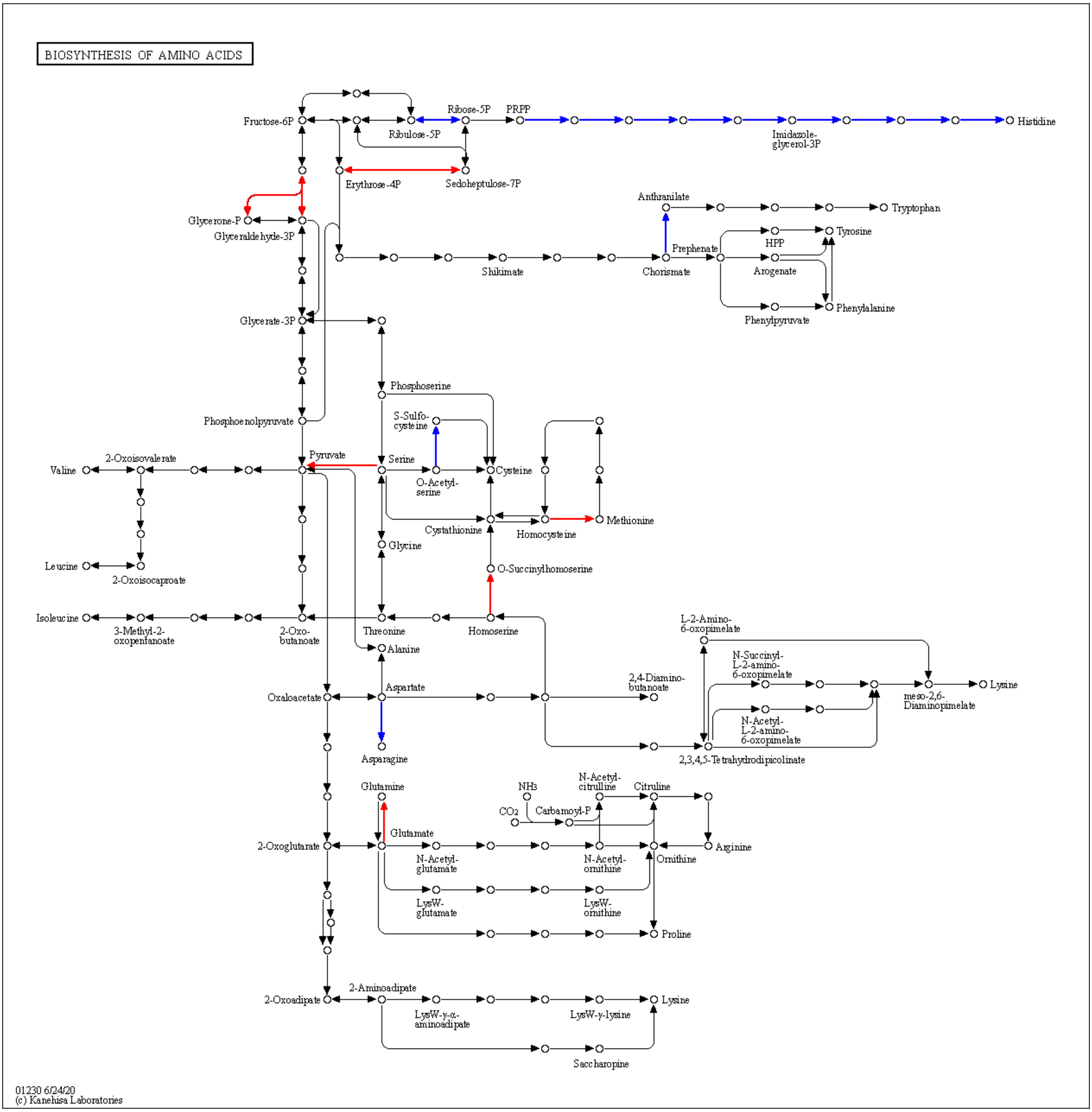
BIOSYNTHESIS OF AMINO ACIDS
Fructose-6P
Ribose-5P
PRPP
Ribulose-5P
Histidine
Imidazole-glycerol-3P
Erythrose-4P
Sedoheptulose-7P
Anthranilate
Tryptophan
Glycerone-P
Glyceraldehyde-3P
Tyrosine
HPP
Prephenate
Arogenate
Shikimate
Chorismate
Phenylalanine
Phenylpyruvate
Glycerate-3P
Phosphoserine
S-Sulfo-cysteine
Phosphoenolpyruvate
2-Oxoisovalerate
Pyruvate
Serine
Valine
Cysteine
O-Acetyl-serine
Methionine
Cystathionine
Homocysteine
Glycine
Leucine
2-Oxoisocaproate
O-Succinylhomoserine
Isoleucine
3-Methyl-2-oxopentanoate
2-Oxo-butanoate
Threonine
Homoserine
L-2-Amino-6-oxopimelate
Alanine
N-Succinyl-L-2-amino-6-oxopimelate
Aspartate
2,4-Diamino-butanoate
Oxaloacetate
Lysine
meso-2,6-Diaminopimelate
N-Acetyl-L-2-amino-6-oxopimelate
Asparagine
2,3,4,5-Tetrahydrodipicolinate
Glutamine
NH3
N-Acetyl-citrulline
Citruline
CO2
Carbamoyl-P
Glutamate
2-Oxoglutarate
Arginine
N-Acetyl-glutamate
N-Acetyl-ornithine
Ornithine
LysW-glutamate
LysW-ornithine
Proline
2-Aminoadipate
2-Oxoadipate
Lysine
LysW-γ-α-aminoadipate
LysW-γ-lysine
Saccharopine
01230 6/24/20
(c) Kanehisa Laboratories

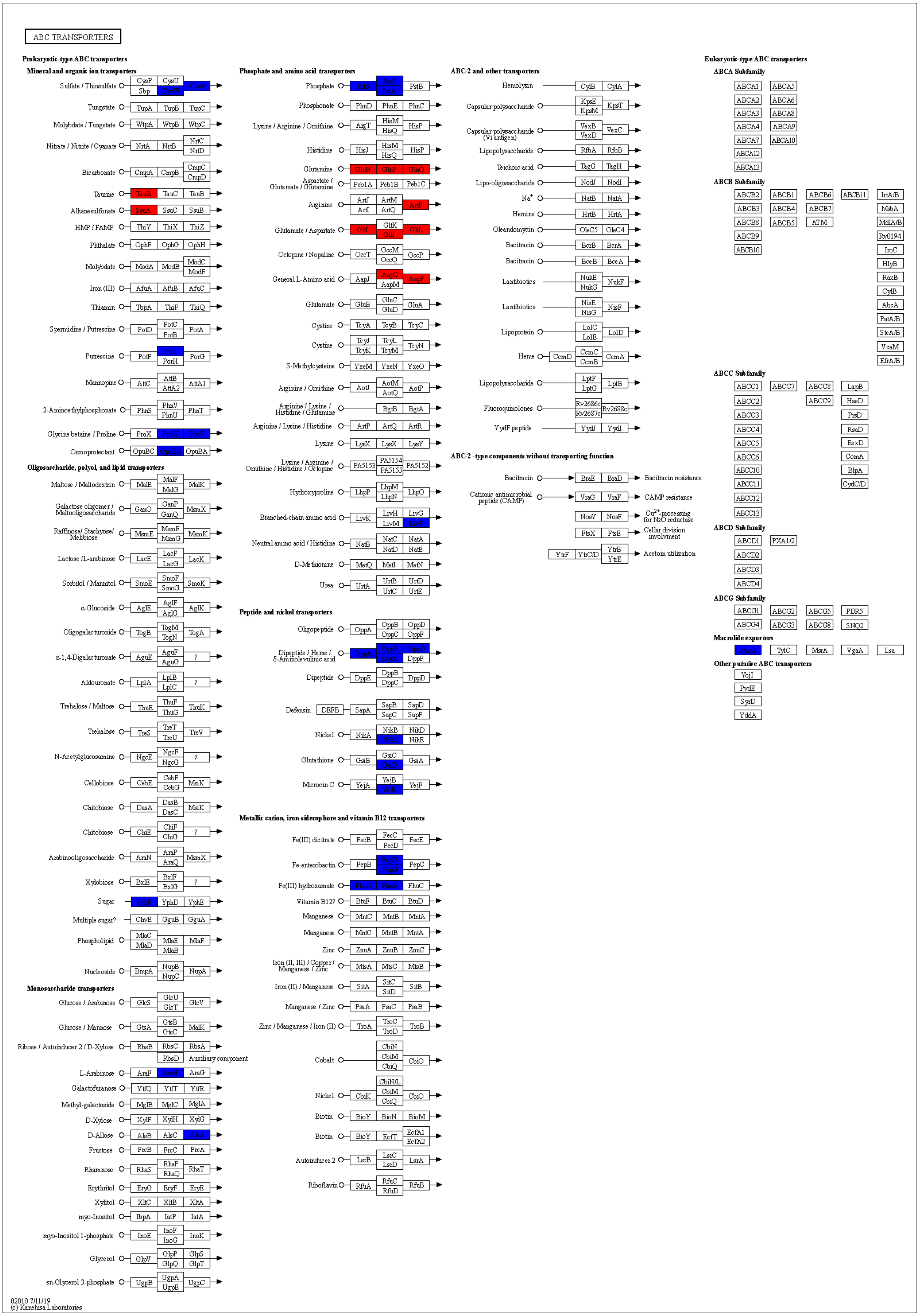

ABC TRANSPORTERS
Prokaryotic-type ABC transporters
Mineral and organic ion transporters
Sulfate / Thiosulfate CysP CysU Sbp CysW CysA
Tungstate TupA TupB TupC
Molybdate / Tungstate WtpA WtpB WtpC
Nitrate / Nitrite / Cyanate NrtA NrtB NrtC NrtD
Bicarbonate CmpA CmpB CmpC CmpD
Taurine TauA TauC TauB
Alkanesulfonate SsuA SsuC SsuB
HMP / FAMP ThiY ThiX ThiZ
Phthalate OphF OphG OphH
Molybdate ModA ModB ModC ModF
Iron (III) AfuA AfuB AfuC
Thiamin TbpA ThiP ThiQ
Spermidine / Putrescine PotD PotC PotB PotA
Putrescine PotF PotI PotH PotG
Mannopine AttC AttB AttA2 AttA1
2-Aminoethylphosphonate PhnS PhnV PhnU PhnT
Glycine betaine / Proline ProX ProW ProV
Osmoprotectant OpuBC OpuBB OpuBA
Oligosaccharide, polyol, and lipid transporters
Maltose / Maltodextrin MalE MalF MalG MalK
Galactose oligomer / Maltooligosaccharide GanO GanP GanQ MsmX
Raffinose/ Stachyose/ Melibiose MsmE MsmF MsmG MsmK
Lactose /L-arabinose LacE LacF LacG LacK
Sorbitol / Mannitol SmoE SmoF SmoG SmoK
α-Glucoside AglE AglF AglG AglK
Oligogalacturonide TogB TogM TogN TogA
α-1,4-Digalacturonate AguE AguF AguG ?
Aldouronate LplA LplB LplC ?
Trehalose / Maltose ThuE ThuF ThuG ThuK
Trehalose TreS TreT TreU TreV
N-Acetylglucosamine NgcE NgcF NgcG ?
Cellobiose CebE CebF CebG MsiK
Chitobiose DasA DasB DasC MsiK
Chitobiose ChiE ChiF ChiG ?
Arabinooligosaccharide AraN AraP AraQ MsmX
Xylobiose BxlE BxlF BxlG ?
Sugar YphF YphD YphE
Multiple sugar? ChvE GguB GguA
Phospholipid MlaC MlaD MlaE MlaB MlaF
Nucleoside BmpA NupB NupC NupA
Monosaccharide transporters
Glucose / Arabinose GlcS GlcU GlcT GlcV
Glucose / Mannose GtsA GtsB GtsC MalK
Ribose / Autoinducer 2 / D-Xylose RbsB RbsC RbsA RbsD Auxiliary component
L-Arabinose AraF AraH AraG
Galactofuranose YtfQ YtfT YtfR
Methyl-galactoside MglB MglC MglA
D-Xylose XylF XylH XylG
D-Allose AlsB AlsC AlsA
Fructose FrcB FrcC FrcA
Rhamnose RhaS RhaP RhaQ RhaT
Erythritol EryG EryF EryE
Xylitol XltC XltB XltA
myo-Inositol IbpA IatP IatA
myo-Inositol 1-phosphate InoE InoF InoG InoK
Glycerol GlpV GlpP GlpQ GlpS GlpT
sn-Glycerol 3-phosphate UgpB UgpA UgpE UgpC
Phosphate and amino acid transporters
Phosphate PstS PstC PstA PstB
Phosphonate PhnD PhnE PhnC
Lysine / Arginine / Ornithine ArgT HisM HisQ HisP
Histidine HisJ HisM HisQ HisP
Glutamine GlnH GlnP GlnQ
Aspartate / Glutamate / Glutamine Peb1A Peb1B Peb1C
Arginine ArtJ ArtI ArtM ArtQ ArtP
Glutamate / Aspartate GltI GltK GltJ GltL
Octopine / Nopaline OccT OccM OccQ OccP
General L-Amino acid AapJ AapQ AapM AapP
Glutamate GluB GluC GluD GluA
Cystine TcyA TcyB TcyC
Cystine TcyJ TcyK TcyL TcyM TcyN
S-Methylcysteine YxeM YxeN YxeO
Arginine / Ornithine AotJ AotM AotQ AotP
Arginine / Lysine / Histidine / Glutamine BgtB BgtA
Arginine / Lysine / Histidine ArtP ArtQ ArtR
Lysine LysX LysX LysY
Lysine / Arginine / Ornithine / Histidine / Octopine PA5153 PA5154 PA5155 PA5152
Hydroxyproline LhpP LhpM LhpN LhpO
Branched-chain amino acid LivK LivH LivM LivG LivF
Neutral amino acid / Histidine NatB NatC NatD NatA NatE
D-Methionine MetQ MetI MetN
Urea UrtA UrtB UrtC UrtD UrtE
Peptide and nickel transporters
Oligopeptide OppA OppB OppC OppD OppF
Dipeptide / Heme / δ-Aminolevulinic acid DppA DppB DppC DppD DppF
Dipeptide DppE DppB DppC DppD
Defensin DEFB SapA SapB SapC SapD SapF
Nickel NikA NikB NikC NikD NikE
Glutathione GsiB GsiC GsiD GsiA
Microcin C YejA YejB YejE YejF
Metallic cation, iron-siderophore and vitamin B12 transporters
Fe(III) dicitrate FecB FecC FecD FecE
Fe-enterobactin FepB FepD FepG FepC
Fe(III) hydroxamate FhuD FhuB FhuC
Vitamin B12? BtuF BtuC BtuD
Manganese MntC MntB MntA
Manganese MntC MntB MntA
Zinc ZnuA ZnuB ZnuC
Iron (II, III) / Copper / Manganese / Zinc MtsA MtsC MtsB
Iron (II) / Manganese SitA SitC SitD SitB
Manganese / Zinc PsaA PsaC PsaB
Zinc / Manganese / Iron (II) TroA TroC TroD TroB
Cobalt CbiN CbiM CbiQ CbiO
Nickel CbiK CbiN/L CbiM CbiQ CbiO
Biotin BioY BioN BioM
Biotin BioY EcfT EcfA1 EcfA2
Autoinducer 2 LsrB LsrC LsrD LsrA
Riboflavin RfuA RfuC RfuD RfuB
ABC-2 and other transporters
Hemolysin CylB CylA
Capsular polysaccharide KpsE KpsM KpsT
Capsular polysaccharide (Vi antigen) VexB VexD VexC
Lipopolysaccharide RfbA RfbB
Teichoic acid TagG TagH
Lipo-oligosaccharide NodJ NodI
Na+ NatB NatA
Hemine HrtB HrtA
Oleandomycin OleC5 OleC4
Bacitracin BcrB BcrA
Bacitracin BceB BceA
Lantibiotics NukE NukG NukF
Lantibiotics NisE NisG NisF
Lipoprotein LolC LolE LolD
Heme CcmD CcmC CcmB CcmA
Lipopolysaccharide LptF LptG LptB
Fluoroquinolones Rv2686c Rv2687c Rv2688c
YydF peptide YydJ YydI
ABC-2-type components without transporting function
Bacitracin BraE BraD Bacitracin resistance
Cationic antimicrobial peptide (CAMP) VraG VraF CAMP resistance
NosY NosF Cu2+-processing for N2O reductase
FtsX FtsE Cellular division involvment
YtrF YtrC/D YtrB YtrE Acetoin utilization
Eukaryotic-type ABC transporters
ABCA Subfamily
ABCA1 ABCA5 ABCA2 ABCA6 ABCA3 ABCA8 ABCA4 ABCA9 ABCA7 ABCA10 ABCA12 ABCA13
ABCB Subfamily
ABCB2 ABCB1 ABCB6 ABCB11 IrtA/B ABCB3 ABCB4 ABCB7 MsbA ABCB8 ABCB5 ATM MdlA/B ABCB9 Rv0194 ABCB10 IroC HlyB RaxB CydB AbcA PatA/B SteA/B VcaM EfrA/B
ABCC Subfamily
ABCC1 ABCC7 ABCC8 LapB ABCC2 ABCC9 HasD ABCC3 PrsD ABCC4 RsaD ABCC5 EexD ABCC6 ComA ABCC10 BlpA ABCC11 CydC/D ABCC12 ABCC13
ABCD Subfamily
ABCD1 PXA1/2 ABCD2 ABCD3 ABCD4
ABCG Subfamily
ABCG1 ABCG2 ABCG5 PDR5 ABCG4 ABCG3 ABCG8 SNQ2
Macrolide exporters
MacB TylC MsrA VgaA Lsa
Other putative ABC transporters
YojI PvdE SyrD YddA
02010 7/11/19
(c) Kanehisa Laboratories

**Supplementary Fig. 10 Representative metabolic pathway maps at 24 h from KEGG mapper.** Red indicates the upregulation and blue indicates the downregulation of each gene.

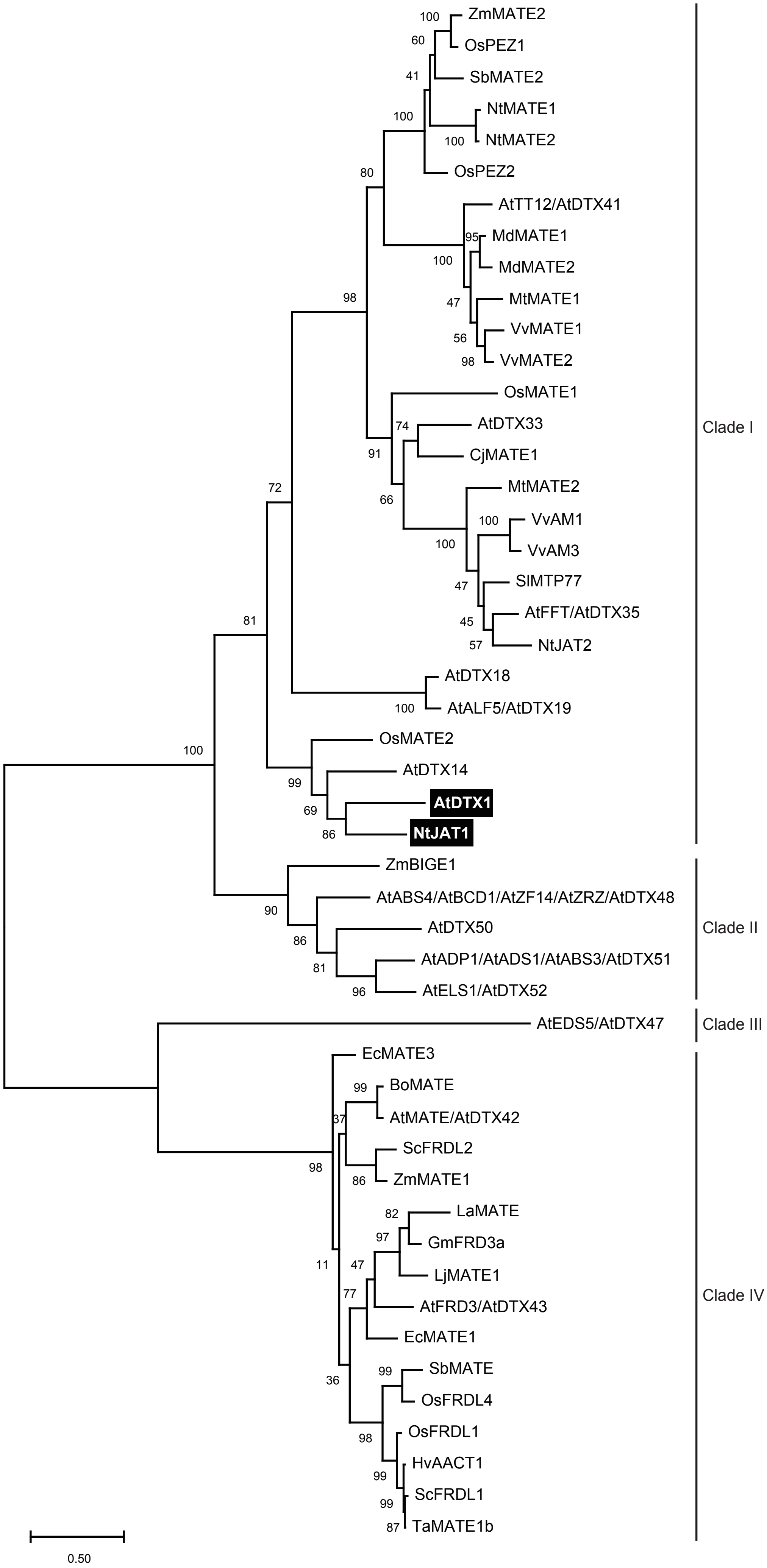
ZmMATE2
OsPEZ1
SbMATE2
NtMATE1
NtMATE2
OsPEZ2
AtTT12/AtDTX41
MdMATE1
MdMATE2
MtMATE1
VvMATE1
VvMATE2
OsMATE1
AtDTX33
CjMATE1
MtMATE2
VvAM1
VvAM3
SlMTP77
AtFFT/AtDTX35
NtJAT2
AtDTX18
AtALF5/AtDTX19
OsMATE2
AtDTX14
AtDTX1
NtJAT1
ZmBIGE1
AtABS4/AtBCD1/AtZF14/AtZRZ/AtDTX48
AtDTX50
AtADP1/AtADS1/AtABS3/AtDTX51
AtELS1/AtDTX52
AtEDS5/AtDTX47
EcMATE3
BoMATE
AtMATE/AtDTX42
ScFRDL2
ZmMATE1
LaMATE
GmFRD3a
LjMATE1
AtFRD3/AtDTX43
EcMATE1
SbMATE
OsFRDL4
OsFRDL1
HvAACT1
ScFRDL1
TaMATE1b
Clade I
Clade II
Clade III
Clade IV
0.50

**Supplementary Fig. 11 Phylogenetic relationship of plant MATE family members.** Plant MATE transporter sequences were aligned using ClustalW and subjected to phylogenetic analysis conducted using the Maximum Likelihood method and Le_Gascuel_2008 model [41] of the MEGAX software [42] with 1,000 bootstraps. The numbers on the branches represent bootstrap values. The scale bar shows the number of amino acid substitutions per site. Many proteins belonging to clade I transport secondary metabolites, such as cyanidin-3-*O*-glucoside, epicatechin 3′-*O*-glucoside, apigenin 7-*O*-glucoside, coumarolyagmatine, nicotine, berberine, palmatine, dhurrin, and protocatechuic acid. Some MATE transporters of clade I transport xenobiotics. MATE transporters of clade II are related to morphogenesis, disease resistance, leaf senescence, and ABA transport. AtEDS5/AtDTX47 (clade III) is involved in disease resistance via salicylic acid transport. Almost all clade IV proteins transport citrate and are implicated in $Al^{3+}$ detoxification or Fe translocation.

The *Arabidopsis* MATE members are as follows: AtTT12/AtDTX41, At3g59030; AtDTX33, At1G47530; AtFFT/AtDTX35, At4g25640; AtDTX18, At3g23550; AtALF5/AtDTX19, At3g23560; AtDTX14, At1g71140; AtDTX1, At2g04070; AtABS4/AtBCD1/AtZF14/AtZRZ/AtDTX48, At1g58340; AtDTX50, At5g52050; AtADP1/AtADS1/AtABS3/AtDTX51, At4g29140; AtELS1/AtDTX52, At5g19700; AtEDS5/AtDTX47, At4g39030; AtMATE/AtDTX42, At1g51340; AtFRD3/AtDTX43, At3g08040. The MATE members of other plant species and their accession numbers are as follows: BoMATE (*Brassica oleracea*), KF031944; CjMATE1 (*Coptis japonica*), BAX73926; EcMATE1 (*Eucalyptus camaldulensis*), BAM68465; EcMATE3, BAM68467; GmFRD3a (*Glycine max*), ACE89001; HvAACT1 (barley), BAF75822; LaMATE (*Lupinus albus*), AAW30732; LjMATE1 (*Lotus japonicus*), BAN59993; MdMATE1 (*Malus domestica*), ADO22710; MdMATE2, ADO22712; MtMATE1 (*Medicago truncatula*),

ACX37118; MtMATE2, ADV04045; NtJAT1 (*Nicotiana tabacum*), CAQ51477; NtJAT2, BAP40098; NtMATE1, BAF47751; NtMATE2, BAF47752; OsFRDL1 (*Oryza sativa*), BAG95121; OsFRDL4, BAL41687; OsMATE1, Os03g08900; OsMATE2, Os05g48040; OsPEZ1, AK243209; OsPEZ2, Os03g0572900; SbMATE (*Sorghum bicolor*), ABS89149; SbMATE2, Sobic.001G012600; ScFRDL1 (rye), BAJ61741; ScFRDL2, BAJ61742; SlMTP77 (tomato), AAQ55183; TaMATE1b (*Triticum aestivum*), AFZ61900; VvAM1 (*Vitis vinifera*), ACN91542; VvAM3, ACN88706; VvMATE1, XP_002282907; VvMATE2, XP_002282932; ZmBIGE1 (*Zea mays*), KT310084; ZmMATE1, ACM47309; ZmMATE2, ACZ55931.

41. Kumar S., Stecher G., Li M., Knyaz C., and Tamura K. (**2018**). MEGA X: Molecular Evolutionary Genetics Analysis across computing platforms. *Molecular Biology and Evolution* **35**:1547-1549.

42. Le S.Q. and Gascuel O. (**2008**). An Improved General Amino Acid Replacement Matrix. *Mol Biol Evol* **25(7)**:1307-1320.

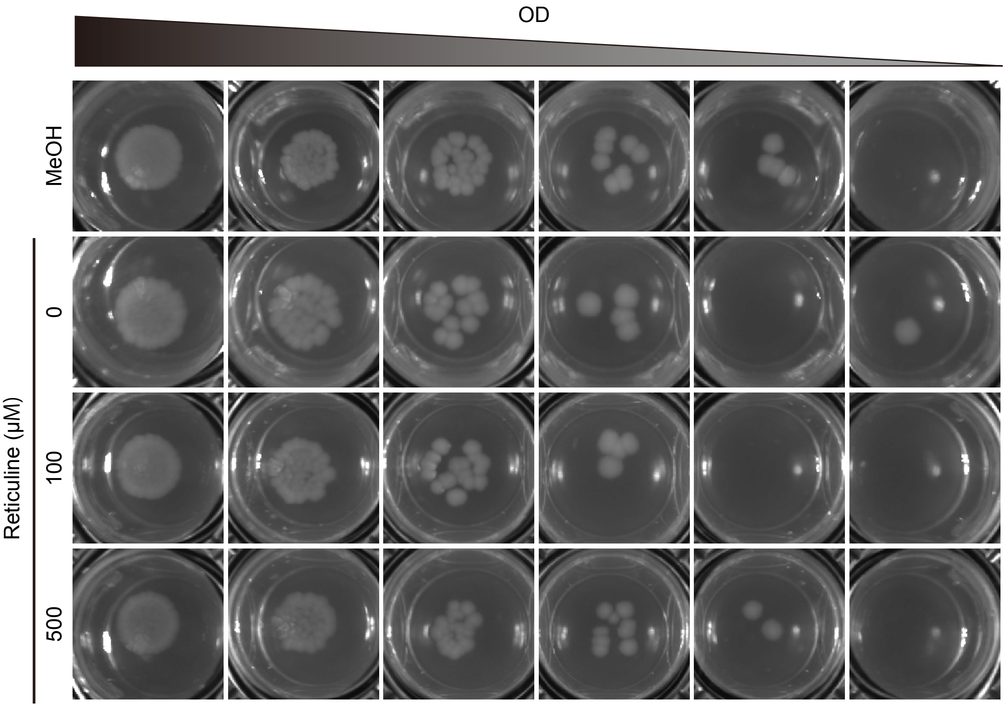


**Supplementary Fig. 12 Growth of *E. coli* in LB medium containing reticuline.** *E. coli* BL21(DE3) harboring pCOLADuet-1 vector were precultured in LB medium at 37°C with shaking at 200 rpm. When the $OD_{600}$ of the cultures reached 1.0, they were diluted to $OD_{600}$ = 0.001. Five microliters of the serially diluted samples were spotted onto a half-strength LB plate containing 0, 100 and 500 μM reticuline and incubated for 36 h at 25°C; 5% (v/v) MeOH was used as the control.
