## Supplementary data legends for "Transport engineering for improving production and secretion of valuable alkaloids in *Escherichia coli*"

**Supplementary Data 1. Summary of RNA-seq data.**

**Supplementary Data 2. Summary of genes induced or suppressed in AtDTX1-expressing cells, with |fold change| ≥ 2.** Statistical analysis was performed using Fold Change. exactTest using edgeR was performed per comparison pair. Significant results were selected based on |fc| ≥ 2 and exactTest raw $P$-value < 0.05.

**Supplementary Data 3. Gene Ontology analysis of significantly altered genes at 0 h (AtDTX1 vs. vector control).** Statistical analysis was performed using Fold Change. exactTest using edgeR was performed per comparison pair. Significant results were selected based on |fc| ≥ 2 and exactTest raw $P$-value < 0.05.

**Supplementary Data 4. Gene Ontology analysis of significantly altered genes at 8 h (AtDTX1 vs. vector control).** Statistical analysis was performed using Fold Change; exactTest was performed using edgeR per comparison pair. The significant results were selected based on |fc| ≥ 2 and exactTest raw $P$-value < 0.05.

**Supplementary Data 5. Gene Ontology analysis of significantly altered genes at 12 h (AtDTX1 vs. vector control).** Statistical analysis was performed using Fold Change. exactTest was performed using edgeR per comparison pair. The significant results were selected based on |fc| ≥ 2 and exactTest raw $P$-value < 0.05.

**Supplementary Data 6 . Gene Ontology analysis of significantly altered genes at 24 h (AtDTX1 vs. vector control).** Statistical analysis was performed using Fold Change. exactTest was performed using edgeR per comparison pair. The significant results were selected based on |fc| ≥2 and exactTest raw *P*-value < 0.05.

**Supplementary Data 7. Gene Ontology analysis of significantly altered genes at 48 h (AtDTX1 vs. vector control).** Statistical analysis was performed using Fold Change. exactTest was performed using edgeR per comparison pair. The significant results were selected based on |fc| ≥ 2 and exactTest raw *P*-value < 0.05.
